# Structural and functional insights into the Rcs phosphorelay

**DOI:** 10.64898/2026.05.08.723598

**Authors:** Melesse Nune, Anushya Petchiappan, Istvan Botos, Nadim Majdalani, Sydney H. Shapiro, Rodolfo Ghirlando, Chin-Hsien Tai, Amila Abeykoon, Ann Marie Stanley, Bridgette M. Beach, Susan Gottesman, Susan K. Buchanan

## Abstract

The Rcs phosphorelay regulates gene expression in response to cell envelope stress and is critical for the virulence of pathogenic bacteria, including *Klebsiella pneumoniae,* due to its regulation of genes related to extracellular capsule, cell division, and motility. The RcsC histidine kinase, RcsD phosphotransfer protein and RcsB response regulator, which form the core of the Rcs phosphorelay, are negatively regulated by the unique inner membrane protein IgaA via interaction with RcsD. An outer membrane lipoprotein, RcsF, activates signaling by interaction with IgaA, but the precise activation mechanisms remain unclear. In this study, we determined the structures of IgaA and the IgaA/RcsF complex using Cryo-electron microscopy (Cryo-EM). We also determined the structures of RcsC and RcsD, which both form homodimers stabilized by hydrophobic interactions, creating ladder-like structures. Combining the Cryo-EM structures, AlphaFold3 structure predictions of IgaA/RcsD and RcsF/IgaA/RcsD, and genetic studies, we describe a model for how RcsF modifies the IgaA/RcsD interaction, lifting negative regulation and activating the Rcs phosphorelay. Our findings provide a high-resolution depiction of the Rcs stress response system and suggest potential targets for small molecule inhibitors.

## INTRODUCTION

The Gram-negative cell envelope is important for withstanding environmental stress and is therefore a primary target for several antimicrobial compounds. Bacteria utilize multiple envelope stress response mechanisms including two-component signaling systems and phosphorelays to monitor and respond to any damage to their membrane^1,2^. One of these mechanisms is the Regulator of Capsule Synthesis (Rcs) phosphorelay^3,4^. The Rcs stress response is critical for maintaining cell wall integrity and is highly conserved across enterobacteria, including pathogens like *Klebsiella pneumoniae*, *Salmonella enterica*, and *Yersinia pestis*^3,5^. This pathway responds to envelope stress from antimicrobial peptides, beta-lactam compounds, and changes in pH and osmolarity^6–8^. It also senses defects in membrane and peptidoglycan biogenesis, LPS alterations, and mechanical stress^5,9,10^. The Rcs cascade regulates transcription of genes related to capsule synthesis, biofilm formation, motility, cell division, and virulence^3,5,11–13^.

A prototypical bacterial two-component system contains a sensor kinase for sensing stress and a response regulator to modulate the cellular response^14^. The Rcs phosphorelay is notably more sophisticated than many of these systems; it comprises the hybrid histidine kinase RcsC and a phosphotransferase RcsD in the inner membrane (IM) along with the cytoplasmic response regulator RcsB (Fig. 1A-B). RcsC possesses two transmembrane domains, a periplasmic domain, and a cytoplasmic region with an active histidine kinase domain (HisKA) and a C-terminal response regulator (Rec) domain (Fig. 1A). RcsD has a similar domain architecture to RcsC (Fig. 1A). It contains two alpha helical transmembrane domains, a periplasmic domain, and a large cytoplasmic domain that harbors, in addition to an inactive histidine kinase domain, the ABL domain and the Hpt phosphotransferase domain^4,15^ (Fig. 1A). The Hpt domain contains the reactive histidine where phosphate is received from RcsC and transferred to the cytoplasmic transcriptional regulator RcsB^16^. Upon induction, the phosphorelay begins with the autophosphorylation of a histidine in RcsC (Fig. 1) followed by transfer of a phosphoryl group to an aspartate in the Rec domain. The phosphate is subsequently transferred to a catalytic histidine in RcsD. Finally, RcsD transfers the phosphate to an aspartate in RcsB. Phosphorylated RcsB triggers homodimerization to regulate genes in the Rcs regulon. RcsB can also heterodimerize with other proteins such as RcsA (in *E. coli*) or RmpA (in *Klebsiella*) to regulate transcription^4,17,18^. Most but not all bacteria with the Rcs system show similar overall activation mechanisms^3,5,19^.

**Figure 1:**
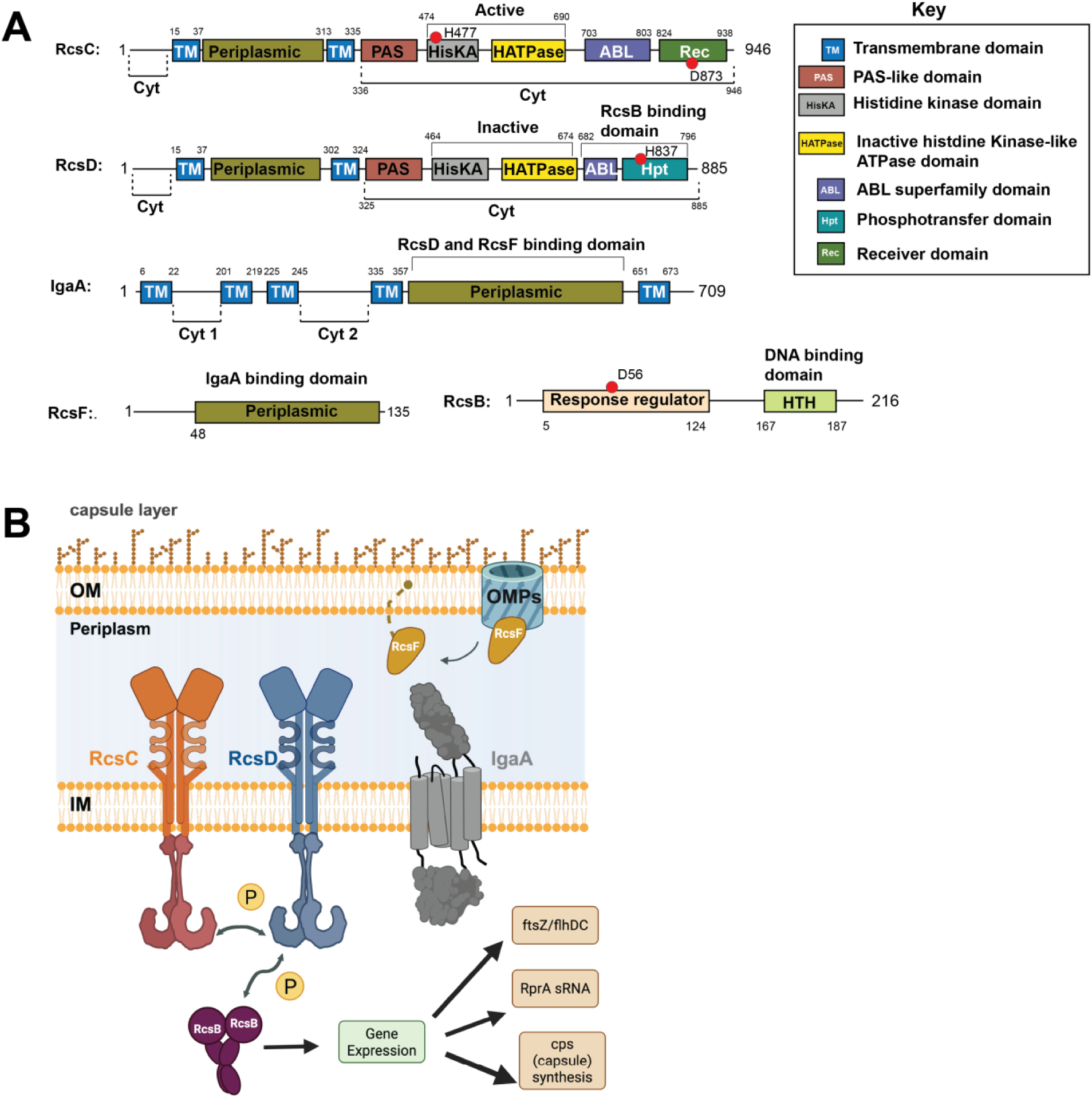
The Rcs phosphorelay system. A) Domain architecture of *Klebsiella pneumoniae* Rcs proteins; not drawn to scale. *E. coli* Rcs proteins are similar in organization and only slightly differ in size and domain borders (**s**ee Fig. S1A). Important residues for the phosphorelay are indicated by red dots. RcsC and RcsD have similar domain architectures, but RcsD contains a Hpt phosphotransfer domain along with an inactive histidine kinase domain. RcsB contains the aspartate residue which is phosphorylated during signaling. RcsF is lipidated, contains a disordered linker and a C-terminal region that interacts with IgaA. The periplasmic domain of IgaA interacts with RcsD/RcsF while the cytoplasmic domains interact only with RcsD. B) Snapshot of the Rcs phosphorelay signaling cascade at start of this project.

In addition to these phosphorelay proteins, this system uniquely involves an essential inner membrane (IM) protein IgaA and an outer membrane (OM) lipoprotein RcsF^20,21^. IgaA is a ∼710 amino acid protein with five predicted transmembrane alpha helices, a cytoplasmic domain, and a large periplasmic domain implicated in binding to RcsF/RcsD^4,22^ (Fig. 1, S1A). During normal growth, IgaA interacts with RcsD to repress signaling, allowing RcsC to function as a phosphatase^4^. IgaA serves as the traffic controller to turn the Rcs phosphorelay OFF/ON; unregulated activation of Rcs is lethal, and thus IgaA is essential^4,21^. RcsF is a small lipoprotein that is anchored in the outer membrane via an N-terminal unstructured region and has a C-terminal periplasmic domain that recognizes IgaA^4,23^ (Fig. 1). RcsF acts as the major sensor for this pathway and induces signaling in response to LPS/peptidoglycan defects or antimicrobial products such as polymyxin B (which likely disrupts LPS) and A22 (MreB inhibitor)^8^.

Previous studies have suggested that RcsF interacts with OM proteins at the cell surface (such as BamA and OmpA) and remains unavailable to interact with IgaA, therefore keeping the Rcs cascade in a “repressed” state^23^. When disruptions occur at the cell surface or peptidoglycan layer, RcsF is released to interact with IgaA in the periplasm through an unknown mechanism^4,23^. This is believed to weaken/alter the IgaA/RcsD interaction allowing the RcsC/RcsD/RcsB phosphorelay to be activated. It is this switch in IgaA binding partners that is the focus of this study; despite years of investigation, the molecular basis for signal transduction is not known. Basic unanswered questions include how the various protein-protein interactions are mediated within the Rcs complex and how signals modulate the switch between the kinase/phosphatase activity. Increased capsule synthesis is a defining trait of hypervirulent *Klebsiella* strains^24^ and RcsB is among the three response regulators in *E. coli* necessary for colonization of mouse intestine^25^, making this pathway a potential drug target for pathogenic enterobacteria.

Here we report the structural and functional characterization of the Rcs proteins from *K. pneumoniae* and *E. coli*. We reveal the full-length structure of IgaA in isolation and in complex with RcsF, identifying the key residues in IgaA critical for RcsF-dependent signaling. Furthermore, using AlphaFold3 modeling we predicted and then tested critical IgaA/RcsD/RcsF contacts necessary for repression and signaling phosphorelay. We also elucidated the structures of RcsD and RcsC homodimers, including their periplasmic and transmembrane regions, providing insights into the importance of dimerization for signaling. We used genetic and biochemical approaches to assay Rcs signaling and define the importance of these various protein interaction networks in Rcs signal transduction. Our results provide a detailed structural picture of the Rcs signaling cascade and suggest targets for small molecule interventions.

## RESULTS

### Characterization of the IgaA/RcsF complex

The Rcs system is found in a range of enterobacterial species, and for the most part, the components and the signaling appear to be conserved. The components of this phosphorelay have been best studied in *E. coli*; however, *K. pneumoniae* Rcs proteins have the same domain architecture and high sequence similarity for RcsF, IgaA, RcsC, and RcsD (Fig. 1, S1A).

To determine the oligomeric states of the Rcs proteins, we performed analytical ultracentrifugation (AUC) analysis on full-length *K. pneumoniae* IgaA (IgaA_Kp_) purified in LMNG detergent (Fig. S1 panel B-C). The data revealed that IgaA predominantly exists as a monomer, with a sedimentation coefficient of 5.33 S and an estimated protein contribution of 82 ± 14 kDa (Fig. S1C). For RcsF_Kp_, AUC analysis yielded a sedimentation coefficient of 0.97 S and an estimated molar mass of 8.4 kDa, consistent with a monomeric state. We also analyzed the IgaA_Kp_/RcsF_Kp_ complex and found that it exists predominantly as a 1:1 complex with a sedimentation coefficient of 5.34 S and an estimated protein mass of 120 ± 18 kDa (Fig. S1 panel C).

IgaA is anchored in the inner membrane by five transmembrane helices (Fig. 1) and contains two cytoplasmic domains (cyt 1 and cyt 2), in addition to a large periplasmic domain. The crystal structure of the IgaA_Ec_ periplasmic domain in complex with RcsF_Ec_ was recently reported^26^, but there is currently no available structure of full-length IgaA, either in isolation or in complex with other Rcs proteins. We used Isothermal Titration Calorimetry (ITC) to show that full-length IgaA_Kp_ interacts strongly with RcsF_Kp_, with a dissociation constant (kD) of approximately 47.5 nM (Fig. S1D).

To visualize interactions between IgaA and RcsF, we determined the Cryo-EM structure of the full-length IgaA_Kp_/RcsF_Kp_ complex at an overall resolution of approximately 5.44 Å (Fig. 2A, S2-3, Supplementary Table 1). Although the moderate resolution precluded detailed interpretation, we were able to confidently assign the major domains of IgaA_Kp_, including the cytoplasmic region, the majority of transmembrane alpha helices (TMs), and the periplasmic region, by fitting Alphafold3-predicted models into the Cryo-EM density map (Fig. 2A-B, S3). In this structure, RcsF_Kp_ binds to the periplasmic domain of IgaA_Kp_ at the periplasmic “crest”.

**Figure 2:**
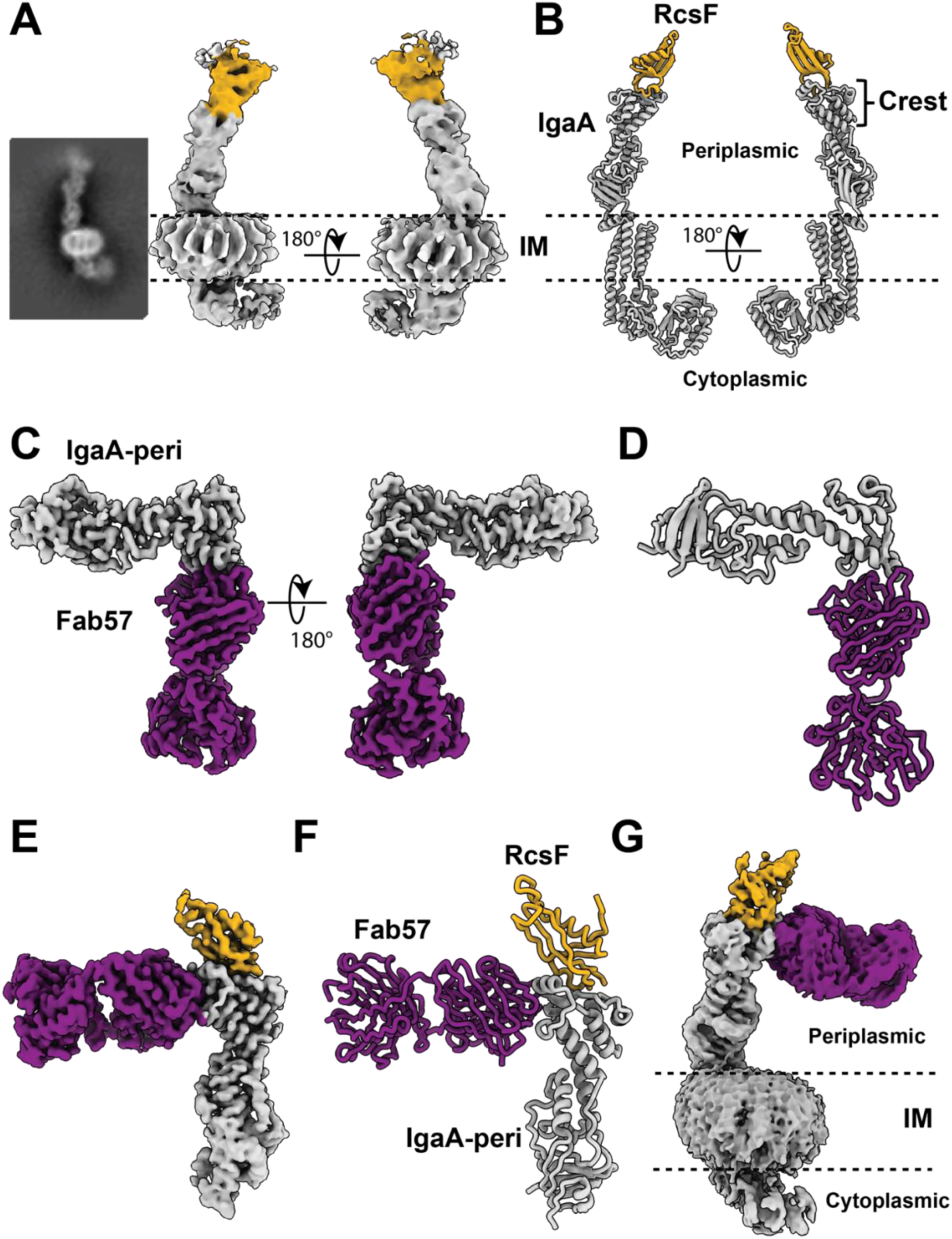
Structural basis for IgaA recognition of RcsF. A) Overall architecture of the IgaA_Kp_/RcsF_Kp_ complex from Cryo-EM reconstruction. RcsF is highlighted in gold. B) AlphaFold3 model of IgaA/RcsF fitted into the Cryo-EM density shown in panel (A). C) High-resolution Cryo-EM map of Apo IgaA-bound Fab57 (purple). Mask excluding the detergent micelle was applied to obtain well-resolved features of the periplasmic domain of IgaA. D) Refined structure of IgaA_Kp_/Fab57 complex. E) High-resolution Cryo-EM map of IgaA_Kp_/Fab57/RcsF_Kp_ complex. Masking was applied to improve IgaA and RcsF periplasmic features. F) Refined structure of IgaA_Kp_/Fab57/RcsF_Kp_ complex. G) Cryo-EM map of full-length IgaA_Kp_ in complex with Fab57.

### A monoclonal antibody improves structural resolution for Apo IgaA_Kp_ and IgaA_Kp_/RcsF_Kp_

To improve the resolution of the IgaA_Kp_/RcsF_Kp_ complex structure, we screened and identified a monoclonal antibody that specifically recognizes purified IgaA_Kp_. We then isolated the antigen-binding fragment (Fab), hereafter referred to as Fab57. We successfully determined the Cryo-EM structure of the IgaA_Kp_/Fab57 complex at a resolution of 2.87 Å, which provided high-resolution details of Fab57 and the periplasmic domain of IgaA_Kp_ (Fig. 2C-D, S4, Supplementary Table 2). This map enabled us to build an accurate model of the periplasmic domain of IgaA_Kp_ (Fig. 2D). Additionally, a separate subset of particles was used to elucidate a moderate-resolution structure of full-length IgaA_Kp_ in complex with Fab57 (Fig. S5A-B, Supplementary Table 1). This structure includes not only the periplasmic domain, but also the cytoplasmic and TM regions. However, electron density around the TMs and cytoplasmic regions is relatively weak. This may be attributed to the inherent flexibility of the cytoplasmic domain, which is characterized by numerous loops and folded regions, resulting in considerable conformational variability.

Fab57 engages the crest of the periplasmic domain of IgaA through the complementarity-determining regions (CDRs) of the Fab. Notably, IgaA_Kp_ periplasmic domain interacts with Fab57 via hydrogen bond interactions and salt bridges (Fig. S5C). Furthermore, we compared the crystal structure of the periplasmic domain of IgaA_Kp_ (9BIZ)^26^ with our Cryo-EM structure of IgaA_Kp_ in complex with Fab57, and observed a strong structural agreement, with an overall root mean deviation square (RMSD) of 0.89 Å (Fig. S5D). This indicates that Fab57 does not disrupt the IgaA_Kp_ structure.

Once we had determined the structure of IgaA_Kp_/Fab57, we next aimed to elucidate the molecular details of IgaA_Kp_ recognition of RcsF_Kp_ using Fab57 as a fiducial marker. We purified the IgaA_Kp_/RcsF_Kp_/Fab57 complex by incubating pre-formed IgaA_Kp_/RcsF_Kp_ with Fab57 and analyzed the complex by Cryo-EM. A subset of particles from this preparation enabled us to resolve a structure at 2.8 Å resolution, consisting of the periplasmic region of IgaA_Kp_ (IgaA-peri), RcsF_Kp_, and Fab57 (Fig. 2E-F, S6, Supplementary Table S2). This high-resolution map allowed us to build an accurate structure for both IgaA_Kp_-peri and RcsF_Kp_. In addition, another set of particles was used to reconstruct a structure that included the full IgaA_Kp_ domains (Fig. 2G, S6-7). Although the resolution for the cytoplasmic and TM domains is modest, it allowed us to fit RcsF_Kp_ and the different domains of IgaA_Kp_ representing the full-length architecture (Fig. S7A-B).

We compared the Apo IgaA_Kp_ and RcsF-bound IgaA_Kp_ structures to identify potential conformational changes occurring in the IgaA_Kp_ periplasmic domain upon RcsF_Kp_ binding. The overall architecture of the two structures exhibits remarkable similarity, with an RMSD of 0.9 Å (Fig. S7C). We next compared our Cryo-EM structure with the previously reported crystal structure of the *E. coli* IgaA_Ec_-peri/RcsF_Ec_ complex (PDB-ID 9BIY)^26^. The two structures exhibit an overall rmsd of 1.3 Å (Fig. S7D). This high degree of similarity suggests that the binding of RcsF does not significantly alter the overall conformation of the IgaA_Kp_ periplasmic domain. However, the resolution of the full-length structure is not adequate to determine whether conformational changes occur within the five-pass TM helices or the cytoplasmic domains when IgaA_Kp_ binds RcsF_Kp_.

We examined the key residues that mediate the interaction between IgaA_Kp_ and RcsF_Kp_, finding multiple points of contact that collectively stabilize the binding interface. A strong interaction is formed by the RcsF_Kp_ loop spanning residues 57-66, which packs against the β-sheet region (β5, see^26^) of IgaA_Kp_ comprising residues 530-539 (Fig. 3A-B). Additional notable contacts include the small IgaA_Kp_ helix consisting of residues 469-478 (α3, see^26^). Overall, the binding interactions bury a surface area of 672 Å^2^ characterized by a salt bridge, hydrogen bonds, and hydrophobic contacts (Fig. 3A-B). A salt bridge is formed between IgaA_Kp_ residue D532 and RcsF_Kp_ R65 (Fig. 3A). A cluster of hydrogen bonds also contributes to the interface stability, involving residues G61/L537, F64/N535, R65/K530, G61/N515 between RcsF_Kp_ and IgaA_Kp_ respectively (Fig. 3B). Other residues participate in hydrophobic interactions, with the strongest contributions coming from IgaA_Kp_ residues L471, I475, M480, L482, V534, R525, and L513 and RcsF_Kp_ residue D66 (Fig. 3B). Together, these diverse and cooperative interactions suggest that the IgaA/RcsF binding interface is finely tuned for both affinity and specificity, likely playing an important role in Rcs signal transduction.

**Figure 3:**
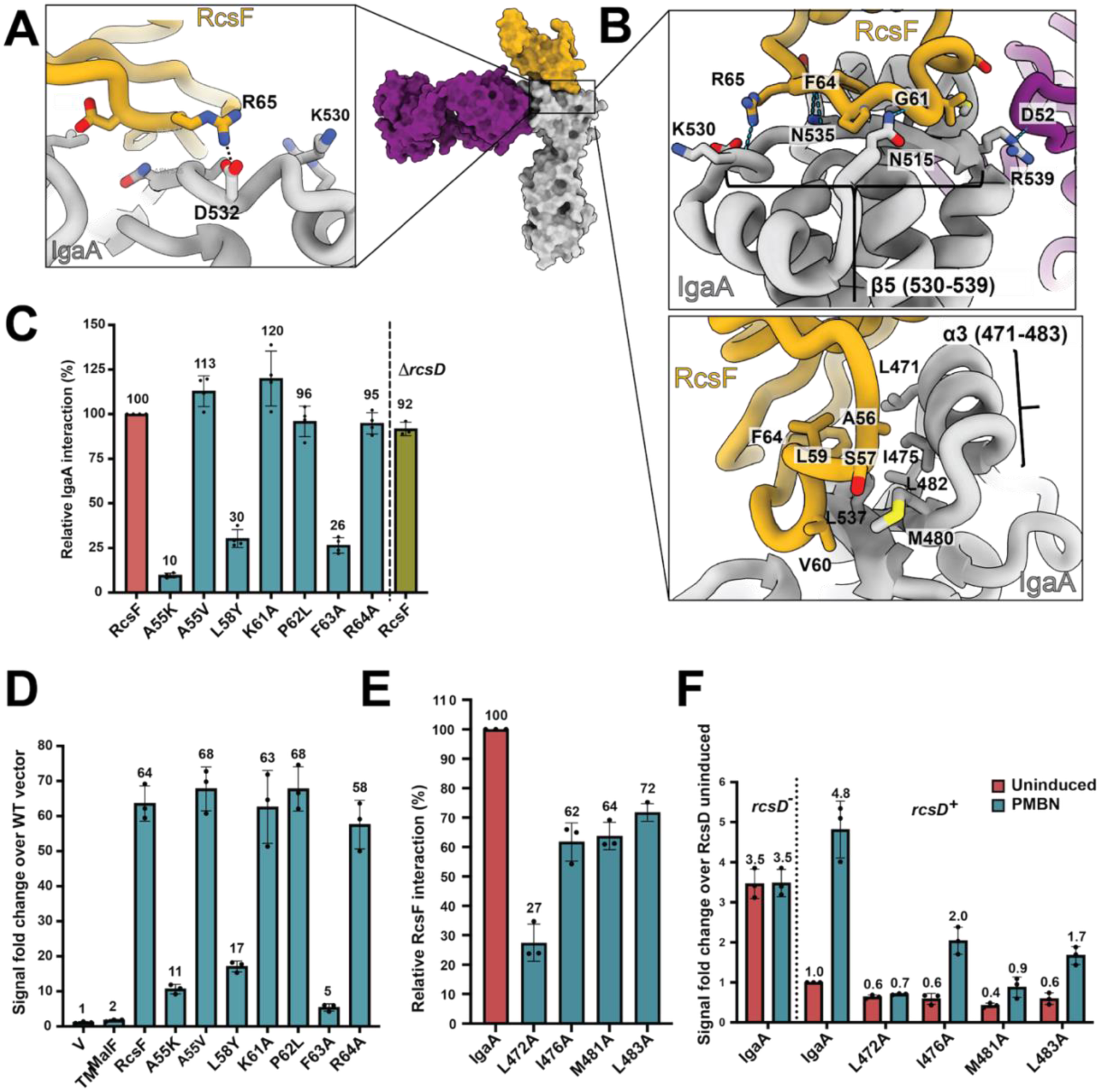
Mutations in IgaA/RcsF interface impact binding and signaling. A-B) Detailed views of IgaA_Kp_/RcsF_Kp_ interaction interface and the residues involved in binding. IgaA_Kp_ is shown in grey, RcsF_Kp_ in gold and Fab in purple. Note that in the assays in this figure and supplemental figures, experiments are all done with *E. coli* mutants and thus the *E. coli* numbering is used (1 residue less than *K. pneumoniae* numbering for both RcsF and IgaA). C) Interaction of RcsF mutants with IgaA. The interaction of *E. coli* IgaA-T18 with RcsF^MalF^-T25 or RcsF^MalF^ point mutants was tested in the strains BTH 101 and BTH101 Δ*rcsD* (EAW12). The interaction of IgaA with RcsF^MalF^ is set to 100 and all other interactions are plotted relative to this. The amino acid residue numbers are for *E. coli* IgaA and RcsF and hence the numbering differs by 1 from those of *K. pneumoniae* above. D) Signaling by RcsF mutants in Δ*rcsF*. RcsF^MalF^-T25 and its point mutants (used in Fig. 3C) were overexpressed in Δ*rcsF* (AP52) and the strain grown in MOPS glycerol minimal medium containing IPTG. The *rprA*-mCherry fluorescence signal at OD 0.4 is depicted here and the signal of pKNT25 vector (V) was set to 1. E) BACTH for IgaA mutants defective in RcsF interaction. The interaction of RcsF^MalF^-T25 tested with IgaA-T18 and IgaA point mutants is plotted here. All interactions are plotted relative to the RcsF^MalF^-IgaA interaction (set to 100). F) Signaling by RcsF-binding defective IgaA mutants. The *igaA* alleles are in the chromosome in a Δ*rcsD* strain carrying a P_rpra_:mCherry reporter fusion. These strains were transformed with either pBAD24 vector (devoid of RcsD) or pEAW11 (*rcsD^+^*) and their fluorescence assayed during growth in MOPS minimal glucose medium (containing 100 g/ml ampicillin). PMBN (20 µg/ml) was added in the medium at the beginning for Rcs activation. The RFU at OD 0.4 for each strain relative to the uninduced *igaA*^+^ *rcsD^+^* strain (set to 1) is plotted.

### Mutations in the IgaA/RcsF binding interface disrupt Rcs signaling

Upon signaling, RcsF is thought to alter its interactions with IgaA, likely loosening the interactions of IgaA with RcsD. Because IgaA acts as a negative regulator of RcsD, this induces signaling. One question we hoped to answer is whether the periplasmic interactions of RcsF with IgaA and RcsD with IgaA are distinct or overlapping. We assayed mutants in RcsF and IgaA for effects on signaling as well as interactions of IgaA with RcsF and RcsD. We previously developed a robust assay for monitoring system-wide Rcs phosphorelay signaling using an P_rpra_::mCherry reporter^4^, a transcriptional fusion of the *rprA* promoter (an RcsB target) to the fluorescent mCherry protein^4^. Signaling is dependent upon RcsC and RcsD as well as RcsB and is repressed by IgaA. Deletion of *rcsF* lowers the basal level of signaling and the cells are no longer capable of activating Rcs in response to signals like polymyxin B nonapeptide (PMBN)^20,27,28^.

We first confirmed that *K. pneumoniae* Rcs proteins are functional in *E. coli* by testing them for complementation of *rcs* deletions (Fig. S8A-C), further supporting that structural and sequence similarities between these proteins are also reflected in functional comparisons. Overexpression of RcsF_Ec_ or RcsF_Kp_ in an *ΔrcsF* strain restored the Rcs response to PMBN (Fig. S8A). The basal signals of *ΔrcsC* and *ΔrcsD* strains are higher than WT, as previously seen^4^, and overexpression of RcsC and RcsD respectively from either *E. coli* or *K. pneumoniae* lowered this signal and also made the strain responsive to PMBN (Fig. S8B). The Rcs homologues from the two species do show minor differences in the basal and induced levels of signaling, as reported earlier for RcsC^29^. To test that IgaA is also functionally conserved, we first replaced the *E. coli* chromosomal *igaA* allele with either *igaA_Kp_* or *igaA_Ec_* in a strain deleted of *rcsD* (to avoid lethality if IgaA is not functional) and then tested for IgaA function by complementing with RcsD expressed from a plasmid; while the uninduced level of expression is higher for IgaA_Kp_ than IgaA_Ec_, it supports both cell growth and PMBN inducibility (Fig. S8C).

We also validated the interaction between the various Rcs proteins using the bacterial two hybrid assay (BACTH)^4,30,31^(Fig. S8D). We previously demonstrated the interaction of *E. coli* IgaA and RcsD using this assay^4^. To test the IgaA/RcsF interaction using this assay we could not use the normally periplasmic RcsF anchored to the outer membrane. Instead we generated a normally periplasmic variant tethered to the transmembrane helix of inner membrane protein MalF; this allows RcsF to be functional at the inner membrane^32^ (see schematic at top of Fig. S8D). This MalF-RcsF variant efficiently interacted with full-length IgaA and IgaA Dcyt but not IgaA^MalF^ MalF transmembrane helix 1 fused to a T25 tag, was used as a control and did not show a high interaction with IgaA. RcsD interacted well with IgaA but not with RcsF. RcsC is non-functional in all of our BACTH assays^4^, so we could not test its interaction with Rcs proteins.

We next sought to evaluate the impact of mutations in the IgaA/RcsF binding interface. All the interfacial amino acid residues in IgaA and RcsF are conserved in *E. coli*, suggesting an identical interaction interface. The Cryo-EM structure of the RcsF_Kp_/IgaA_Kp_ complex suggests a small but strong binding interface. We validated this interaction by mutating the *E. coli* IgaA and RcsF residues at the interface, first measuring the interaction using the BACTH assay as above and then testing appropriate mutants for effects on the activity of the phosphorelay, using the P*_rpra_*::mCherry reporter. We first tested the ability of RcsF_Ec_ mutant in the RcsF_Ec_/IgaA_Ec_ contact region defined by the structure to interact with IgaA_Ec_ (Fig. 3A-C). A55K, L58Y, and F63A had decreased interaction with IgaA_Ec_, while A55V, K61A, P62L, and R64A did not show any significant changes, consistent with previous results^32^. The absence of RcsD_Ec_ had no effect on the wild-type IgaA_Ec_/RcsF_Ec_ interaction determined using BACTH (Fig. 3C). The effects of these RcsF_Ec_ mutants, expressed from a plasmid, on Rcs signaling were tested in a Δ*rcsF* strain carrying the P_rprA_-mCherry reporter (Fig. 3D). RcsF_Ec_ mutants that interacted well with IgaA_Ec_ in the BACTH assay were effective for signaling, while mutants A55K, L58Y, and F63A showed very low levels of activation, consistent with impaired IgaA_Ec_ interaction (Fig. 3D). It is to be noted that for these Rcs signaling assays, RcsF_Ec_ was replaced with the same RcsF^MalF^-T25 variants used in the BACTH. Therefore, we are quantifying the Rcs activation at the inner membrane, which is reported to be high and constitutive (note the 60-fold change over the vector control for the RcsF_Ec_ wild-type derivative in Fig. 3D, compared to the 6-fold change seen when WT RcsF_Ec_ was overexpressed in the absence of PMBN, Fig. S8A).

Next, we used the BACTH assay to interrogate the amino acid residues in IgaA_Ec_, generating structure-guided point mutations of IgaA residues at the binding interface and other conserved IgaA periplasmic residues as controls (Fig. 3E and Fig. S8E). Most of the single *igaA* mutations decreased interaction only slightly or not at all (Fig. S8E), suggesting more flexibility for the IgaA surface than was seen for RcsF. The residues that were most critical for the IgaA_Ec_/RcsF_Ec_ interaction were primarily hydrophobic in nature; alanine mutations in the highly conserved L472 and I476 present in the alpha-helix (α3, see^26^), L483 in the beta-strand, and M481 in the loop had the maximum effects on RcsF interaction (Fig. 3E). These alleles were introduced into the chromosomal copy of *igaA* in an *E. coli* strain deleted for *rcsD* and carrying the P_rprA_:mCherry reporter. In a *ΔrcsD* strain, the basal signal is higher than a strain containing RcsD but there is no Rcs activation by known Rcs activators such as polymyxin B nonapeptide (PMBN)^4^ (Fig. 3F). Expression of RcsD_Ec_ from a plasmid in the *igaA*^+^ Δ*rcsD* strain lowered the basal signal and this strain became PMBN-inducible (compare first two columns to columns three and four, Fig. 3F). However, the IgaA_Ec_ mutant strains, particularly L472A and M481A, had a lower level of basal fluorescence, similar to a Δ*rcsF* strain, and were not activated by PMBN, consistent with loss of RcsF interaction. Because these have a low basal level, they must still be able to “repress” the phosphorelay, presumably by interacting with RcsD, but have become blind to RcsF and therefore are uninducible. It is worth noting that even a partial defect in the BACTH interaction (as for M481A) was very defective for induction, consistent with the idea that the RcsF^MalF^, used to measure interaction but not present for the activity assays in Fig. 3F, may suppress some defects in the IgaA/RcsF interaction interface. Finally, our structure suggests that these IgaA residues define a hydrophobic interaction interface with RcsF residues like F63A and are consistent with the RcsF/IgaA periplasmic domain crystal structure^26^.

Other mutants in IgaA had modest or negligible effects on IgaA interaction and signaling (Fig. S8E-F). Alanine mutations in the conserved beta-sheet residues V535, N536, V537, and L538, present at the interface and defined as a hydrogen binding network by Watanabe and Savchenko had only modest effects on the RcsF interaction as did the residues N516, K531, and D533 (Fig. S8E). Combining L538A with L483A led to loss of interaction (Fig. S8); activity of the phosphorelay was higher but RcsF-blind as was a L472A L483 mutant. Other IgaA residues such as R506, N513, and R540, which are further away from the RcsF binding interface, also did not show any changes in RcsF interaction (Fig. S8E) but were found to seriously perturb RcsD interaction and activity (discussed below). A previously studied mucoid IgaA mutant L643P (present in a beta-sheet distant from the RcsF binding surface) and IgaA mutated in all four periplasmic cysteine residues (C404S, C425S, C498S, C504S-referred to as ‘C4S’) still interacted well with RcsF (Fig. S8E). Both IgaA L643P and C4S have previously been shown to be defective for RcsD interaction^4,33,34^. None of the mutants showed complete disruption of IgaA/RcsF interaction. This may be in part because we measure the interaction using the inner-membrane anchored RcsF, quite likely leading to a stronger interaction than found with the outer-membrane anchored RcsF, and these single alanine mutations are insufficient to abrogate this strong interaction.

### Structures and functions of RcsD and RcsC

The structures and data discussed above establish the interactions of RcsF and IgaA. Work from us and others suggests that the RcsF interaction with IgaA affects the way in which IgaA interacts with RcsD, which in turn suppresses signaling via its interactions with the histidine kinase RcsC. Relatively little is known about these downstream aspects of the signaling pathway. Here, we determined the nature of RcsC and RcsD dimer formation and found that while these proteins have similar overall structures, they form dimers in different ways.

#### RcsC and RcsD form homodimers

Structurally, RcsC and RcsD are predicted to be very similar, with the notable distinction that RcsD contains an inactive histidine kinase domain and a Hpt domain^4^ (Fig. 1A). Most membrane associated histidine kinases of this broad family function as homodimers but no full-length structures exist for any members of this family^35^. Due to their predicted structural and sequence homology it has been hypothesized that both RcsC and RcsD form homodimers, although this had not been experimentally validated^4^. To investigate the oligomeric states of RcsC and RcsD, we purified full-length proteins from *E. coli* and *K. pneumoniae* and subjected them to size-exclusion chromatography and analytical ultracentrifugation. As shown in Fig. S9, SEC analysis revealed the presence of multiple species for both RcsC and RcsD, eluting at distinct volumes. Subsequent AUC analysis of these fractions indicated significant differences in molecular weights across the peaks. For RcsD (both Ec and Kp), the early fractions (P1, boxed green in Fig. S9A and C) contained predominantly aggregated and polydisperse species.

However, the later fractions (P2, boxed green in Fig. S9A and C) revealed a more stable RcsD species with a sedimentation coefficient of 7.77 S that corresponds to a protein contribution of approximately 190 kDa, consistent with a homodimeric form (Fig. S9E). Similarly, for RcsC_Ec_, earlier fractions displayed aggregation and polydispersity, while later fractions yielded a clean dimeric species with a sedimentation coefficient of 7.82 S, corresponding to a protein mass of 231 kDa (Fig. S9D, S9F). In contrast, the AUC for RcsC_Kp_ was mostly aggregated and did not yield discrete species (Fig. S9F). These findings confirm that both RcsC and RcsD form stable homodimers under the experimental conditions, supporting their proposed role in signal transduction and further highlighting the structural parallels between the two proteins.

#### Structures of RcsD from K. pneumoniae and E. coli

Previous structural studies of RcsD and RcsC have been limited to NMR analysis of the ABL-Hpt domain of RcsD_Ec_ (residues 688–890) and phosphoreceiver domain of RcsC_Ec_ (residues 700-949, Fig. 1A)^36,37^. These structures provided limited information regarding the overall architecture and function of RcsD and RcsC. To gain further structural insights, we investigated the full-length RcsD and RcsC structures using Cryo-EM. For the first time ever, we have determined the structures of RcsD_Kp_ at 3.16 Å and RcsD_Ec_ at 2.6 Å resolution, respectively (Fig. 4 and Fig. S10-11, Supplementary Table S2). Although the full-length protein was vitrified during sample preparation, we observed clear density only for the periplasmic domains and the two transmembrane (TM1/TM2) domains (Fig. 4A-B), covering residues ∼9-329 for RcsD_Kp_ and ∼18-335 for RcsD_Ec_ (Fig 4C-D). The cytoplasmic domains of both proteins were not visible in our structures. Consistent with our AUC results, the Cryo-EM structures confirmed that RcsD_Kp_ forms symmetrical homodimers with an overall interface area of 1684 A^2^ (1613 A^2^ for RcsD_Ec_), stabilized by key interacting residues between the monomers. Overall, the two structures share architectural similarity, and the residues responsible for mediating dimerization are well conserved between the two organisms, suggesting a preserved structural and functional mechanism (Fig. 4A-D). Notably, an alpha helix (**α1**, Fig. S12A) from each monomer extends from the beginning of the TM region to the end of the periplasmic domain, spanning residues R15 to I72 (Fig. 4A-D, S18 to T81 for *E. coli*). This long helix plays a critical role in the overall stability and organization of both the transmembrane and periplasmic regions. In addition, another important helix (**α2**, Fig. S12A) from each chain (Kp S113 to L129 and Ec residues N119 to L135) packs against one another, stabilizing the dimer interface. The transmembrane domains come face-to-face to form a four-helix bundle (Fig 4A-D).

**Figure 4:**
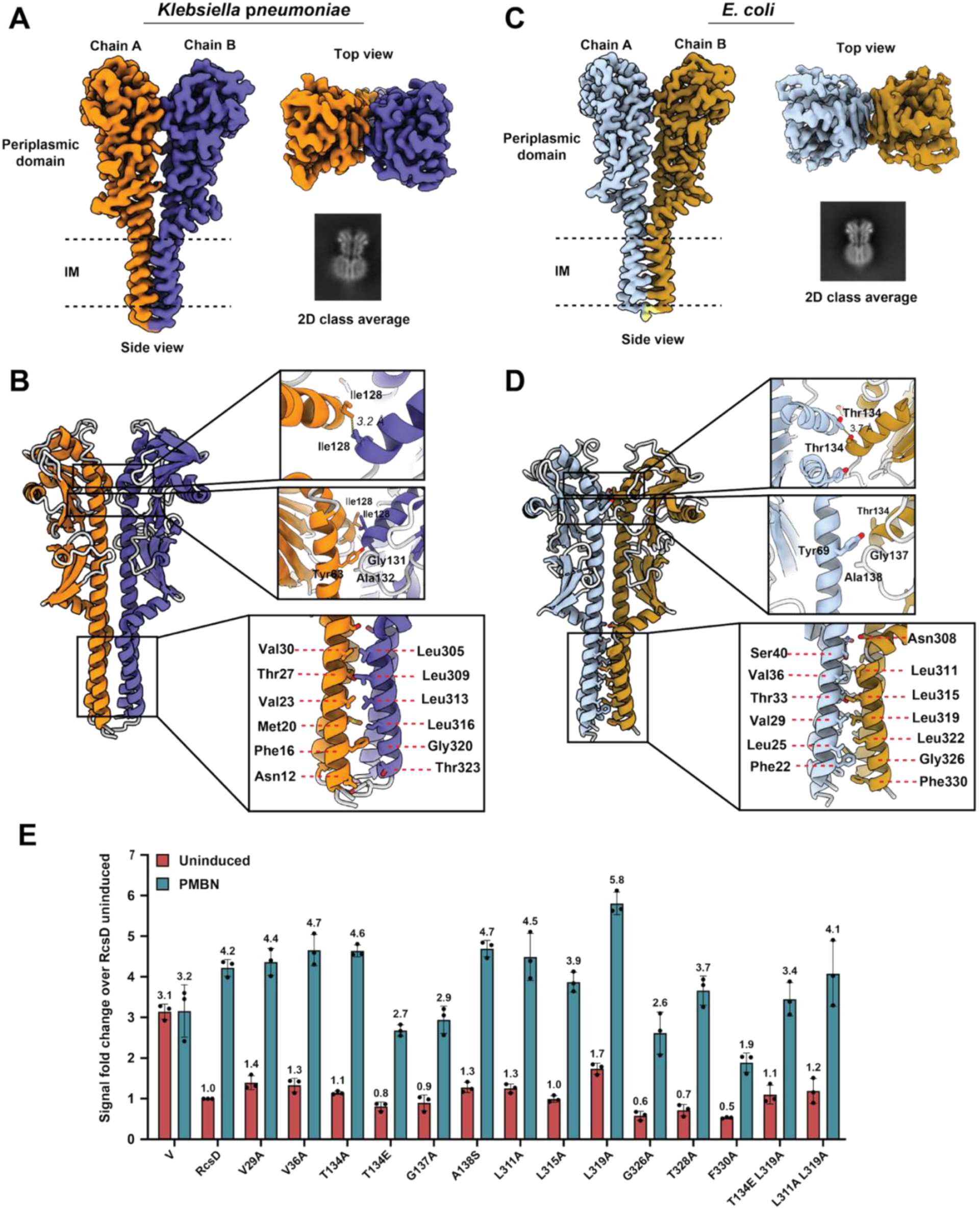
Overview of RcsD dimers and structure-guided mutational analysis. A) Multiple views of the RcsD_Kp_ Cryo-EM structure, highlighting the overall architecture. The detergent micelle has been removed for clarity. B) Refined structure of RcsD_Kp_ with detailed structural features of the protein and close-up views of the dimerization interface, showing key residues and interactions involved in dimer stabilization. C) Multiple views of the RcsD_Ec_ Cryo-EM structure, highlighting the overall architecture. The detergent micelle has been removed for clarity. D) Refined structure of RcsD_Ec_ with detailed structural features of the protein and close-up views of key residues and interactions involved in dimer stabilization. E) Signaling by RcsD_Ec_ dimer interface mutants. Strain EAW19 (Δ*rcsD*) was transformed with either pBAD24 vector, or RcsD_Ec_ (pEAW11) or its mutants and the fluorescence assayed in MOPS minimal glucose medium. PMBN was used for Rcs induction.

Analysis of the side chains at the interface revealed that dimerization in the transmembrane and periplasmic domains is primarily stabilized via hydrophobic interactions. Specifically, Y63 (Y69 in *E. coli*) from one chain forms critical contacts with G131 and A132 (G137 and A138 in *E. coli*) of the opposing chain. Importantly, a hydrophobic bridge formed by I128 (T134 in *E. coli*) from both chains appears to make the strongest contribution to dimer stability (Fig. 4B and 4D). Similarly, within the transmembrane regions, dimerization is mediated by a combination of conserved hydrophobic interactions and van der Waals contacts between residues on the two chains (see Fig. 4B and 4D). These interactions help anchor the two monomers in the membrane, ensuring the structural integrity of the dimer across both the membrane and periplasmic regions. Additionally, although only the TMs and periplasmic domains are resolved, based on AlphaFold3 modeling and our structure, we anticipate that the stable dimer interface extends continuously from the periplasmic domain to the cytoplasmic domain (Fig. S12B-C).

#### RcsD dimer interface mutants show minor impacts in signaling

The structures determined for RcsD as well as the AlphaFold3 prediction (for cytoplasmic domains) all suggest that the dimer interface may be very stable, and thus difficult to disrupt with point mutants. This was directly tested by asking if mutations in the RcsD periplasmic and transmembrane dimer interface residues impact RcsD-dependent signaling (Fig. 4E). WT RcsD_Ec_ complementation of the Δ*rcsD* strain leads to a decrease in basal level expression, with PMBN able to increase the signal 4-fold. As shown in Fig. 4E and Fig. S12D, we did not see major differences in signaling upon testing most mutant derivatives at the dimer interfaces, including periplasmic mutants and transmembrane four-helix bundle mutants. L319A had a somewhat increased Rcs basal and induced signal, although it seems somewhat unlikely that this is due to altered dimerization given that a double mutant (L311A/L319A) is very similar to WT. The mutants T134E (I128 in Kp) and G137A (G132 in Kp) were less inducible than WT RcsD, suggesting a possible role for this region in signal transmission. T134A was more like WT, as was a double mutant of T134E and L319A, making it difficult to interpret the modest change in inducibility for T134E. A set of mutations in adjacent residues in the adjacent transmembrane helix (Chain B) of RcsD_Ec_, G326A, T328A and F330A, all had lower basal levels of signaling, and varied in their inducibility, suggesting a structural role for these residues in the conformation of RcsD. None of the RcsD point mutants tested showed any change in IgaA interaction (Fig. S12E).

We could measure the dimerization of RcsD_Ec_ and various domain deletion constructs in our BACTH assay, although we observed a relatively low beta-galactosidase activity, high heterogeneity and slow growth upon overexpression of these constructs (Fig. S12F). The cytoplasmic constructs alone did not form stable dimers in BACTH although such constructs are capable of unregulated signaling from RcsC to RcsB^4^. Somewhat surprisingly, the interaction increased significantly in the absence of the periplasmic domain, as well as for Δ*peri* derivatives also carrying the RcsD V29A mutant at the four-helix bundle interface (Fig. S12F, Fig. 4D). RcsD L319A Δperi, on the other hand, did not dimerize well (7-fold less than the RcsD Δperi construct). RcsD Δperi and the mutants V29A and L319A did not show any changes in stability under these conditions (Fig. S12G).

These results support a robust dimerization interface, perturbed in a complex way by the periplasmic region and its interaction with the trans-membrane portions of the protein. The high degree of conservation in the types of residues and the overall structure of the RcsD dimer interface in *E. coli* and *K. pneumoniae* supports a likely critical role for this region. Under normal signaling conditions, the periplasmic domain of RcsD is likely interacting with the periplasmic region of IgaA. We speculate that signals from the periplasm may be transmitted to the cytoplasm via subtle changes in this important TM interface^4,33^.

#### Structures of RcsC from K. pneumoniae and E. coli

In our hands, full-length RcsC_Kp_ proved to be relatively unstable for structural studies. Therefore, we purified the stable region of RcsC_Kp_, encompassing the periplasmic domain and TM1/TM2 domains (residues 1-363, Fig. 1A) and determined its structure at 3.74 Å resolution using Cryo-EM (Fig. 5A-B, S13, Supplementary Table S2). In contrast, the full-length RcsC_Ec_ was well-behaved and we were able to purify it without issue. We determined the structure of RcsC_Ec_ at 2.72 Å resolution with Cryo-EM (Fig. 5C-D, S14, Supplementary Table S2). As with the RcsD dimer, we were only able to visualize the periplasmic and TM1/TM2 domains. Based on the 2D class averages (Fig. S13-14), we did not detect any density for the large cytoplasmic domain, likely due to intrinsic flexibility in that region. RcsC forms a symmetrical homodimer, stabilized by interactions within both the periplasmic domain and the transmembrane helices (Fig. 5). We observed notable structural similarity between the periplasmic domains of RcsC and RcsD (Fig. 4-5, S15A-C). However, unlike RcsD, the transmembrane helices of RcsC align along a vertical axis, adopting a sheet-like orientation (Fig. 5A-D). By comparison, the transmembrane helices of RcsD pack against each other to form a four-helix bundle. The functional implications of the distinct TM helix packing for RcsC and RcsD remain to be explored but suggest that RcsC and RcsD would not form a 1:1 heterodimer.

**Figure 5:**
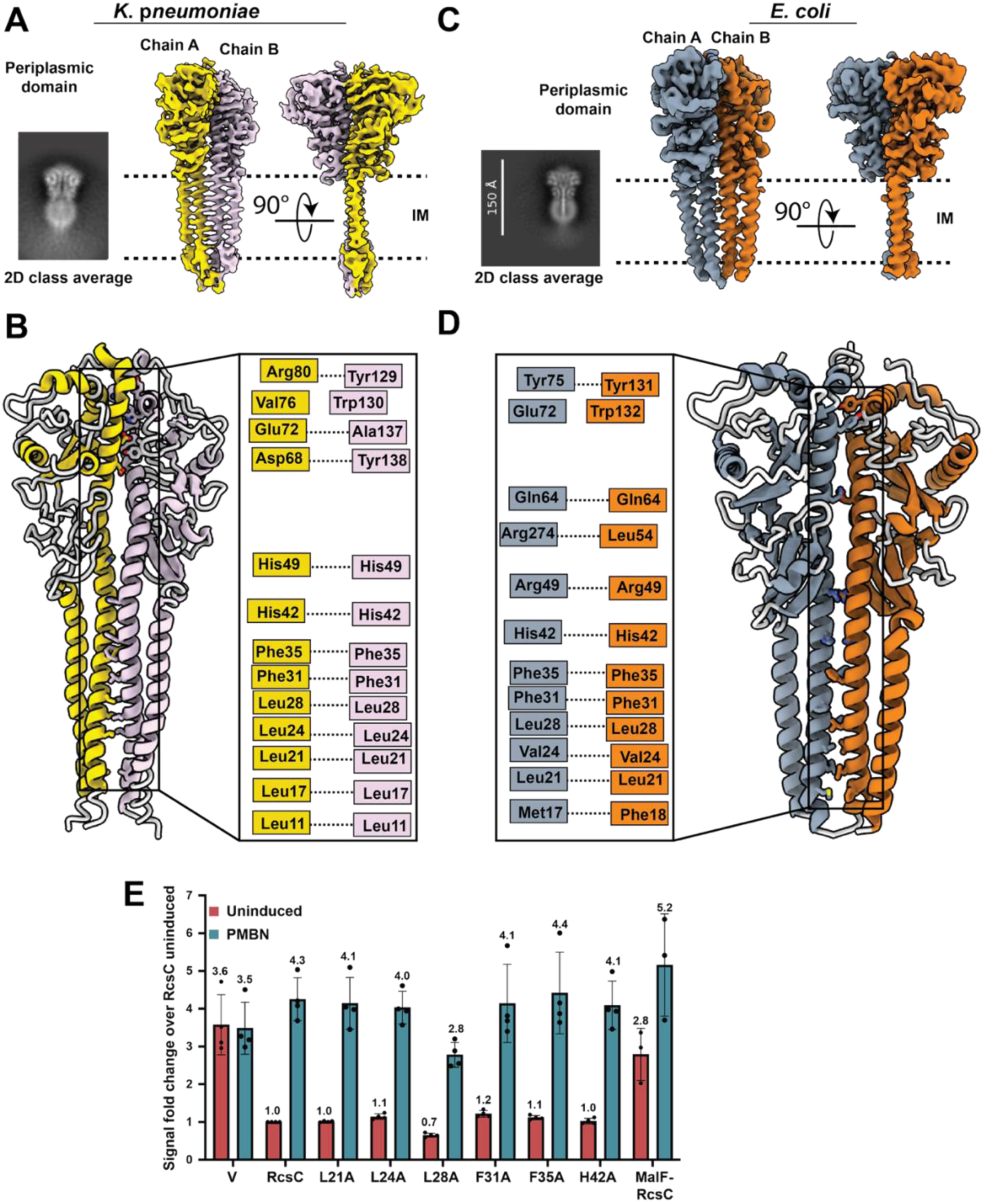
Overview of RcsC dimer structure and structure-guided mutational analysis. A) Multiple views of the RcsC_Kp_ Cryo-EM structure, including 2D projection image, and the Cryo-EM map covering periplasmic and TM1/TM2 domains. B) Refined structure of RcsC_Kp_ with detailed structural features of the dimer interface. C) Multiple views of the RcsC_Ec_ Cryo-EM structure, including 2D projection image, and the Cryo-EM map covering periplasmic and TM1/TM2 domains. D) Refined structure of RcsC_Ec_ with detailed structural features of the dimer interface. E) Signaling by RcsC dimer mutants. Strain EAW91 (Δ*rcsC*) was transformed with either pBAD24 vector derivatives encoding the RcsC mutants and their fluorescence assayed as described in 4E.

Overall, the structures of RcsC from both organisms reveal almost identical architecture and features. Notably, a long helix (**α1**, Fig. S15D, residues 2-82 in Kp, and 11-81 in Ec) encompassing part of the cytoplasmic, the TM, and the periplasmic domains establishes the dimer interface by bringing the monomers face-to-face in a ladder-like arrangement (Fig. 5A-D). This dimer interface is almost entirely formed by conserved key hydrophobic residues, covering an extensive binding interface area of 2607 A^2^ (2283 A^2^ for Ec). In the TM region, F7, L21, L24, L28, F31, F35, and H42 stabilize the dimer interface (Fig 5C, see Fig. 5D for *E.coli* residues). Y138, A137, and conserved W130, and Y129 (Fig 5C, see Fig. 5D for *E.coli* residues) play a role in stabilizing the periplasmic side. Furthermore, a salt bridge formed between R131/D85 (in Kp) and hydrogen bond interactions formed by Y75/D134 (in Ec) stabilize the periplasmic end of the dimeric structure. A disulfide bridge formed by C109/C152 on both monomers (C111/C154 in Ec) provides structural support. However, we have previously shown that neither the periplasmic region of RcsC nor these cysteines are needed for the normal RcsF-dependent induction of the Rcs phosphorelay^33^. The hydrophobic nature of conserved interactions in the dimer interface extends to the cytoplasmic domain as revealed by the AlphaFold3 structures shown in Fig. S15E-F.

We next probed how point mutations within the long helix involved in dimerization impact Rcs signaling *in vivo*. We expressed the corresponding RcsC mutants in an RcsC deletion strain to measure the fluorescence with our reporter assay (*E.coli*). We found no major change in signaling by any of the RcsC point mutants; only L28A had a modestly lower basal level (Fig. 5E). The RcsC structures resemble a ladder, with the elongated alpha helices representing the vertical poles and the interacting sidechains functioning as the rungs (Fig. 5B and 5D). The observation that the dimer interface mutants have minimal impact on signaling is not surprising. This is akin to removing individual rungs from a ladder; while it may compromise local stability, it does not entirely destabilize the overall structure and function of the ladder.

Our previous studies demonstrate that neither the periplasmic nor TM regions of RcsC are critical for RcsF-dependent Rcs induction. Deletion of the periplasmic region is functional for most but not all inducing signals^33^. A chimeric RcsC fused to the MalF TM helix shows a higher basal activity (akin to RcsC_Kp_ in Fig. S8B) but is capable of PMBN induction^4^. Overall, combined with the structures and mutants tested here, we speculate that the sequences of the TM helices are not critical for activity, but that changes in them may impact the regulation of kinase/phosphatase activity and/or interaction with RcsD.

### RcsF and RcsD have both shared and distinct binding sites on IgaA

Despite multiple attempts, we were unable to obtain an experimental structure of the RcsF/IgaA/RcsD complex. We therefore generated a model of the *E. coli* complex using Alphafold3^38^ combined with our Cryo-EM structures as templates. The RcsD 1-461 fragment, previously shown to be sufficient for IgaA interaction^4^, was modelled as a dimer and IgaA and RcsF as monomers. The AF3 model (Fig. 6A) suggests that RcsD and RcsF interact with IgaA in the same region of the periplasmic domain, raising the possibility that they compete for interaction with IgaA. The model correctly predicted the RcsF/IgaA binding interface resolved by Cryo-EM and validated by mutational analysis *in vivo* (Fig. 2-3). Specifically, as shown in Fig. 6B, IgaA is predicted to interact with RcsD via the alpha-helix containing the conserved IgaA polar residues R506, N509, and N513. This model also suggests that RcsF (residues G60, K61, and K134) and RcsD (residues Q221, H224, L225) interact with each other while bound to IgaA (Fig. 6C). However, mutations in K61 (Fig. 3D) or in the RcsD residues near to it (Fig. 6H) did not disrupt the ability to induce the phosphorelay, suggesting that this interaction is weak or not critical for Rcs activation. These RcsD residues do weaken the RcsD interaction with IgaA, leading to a higher basal level. The model also predicted a cytoplasmic interaction interface between the IgaA cyt1 domain and the RcsD PAS-like domain (Fig. 6D).

**Figure 6:**
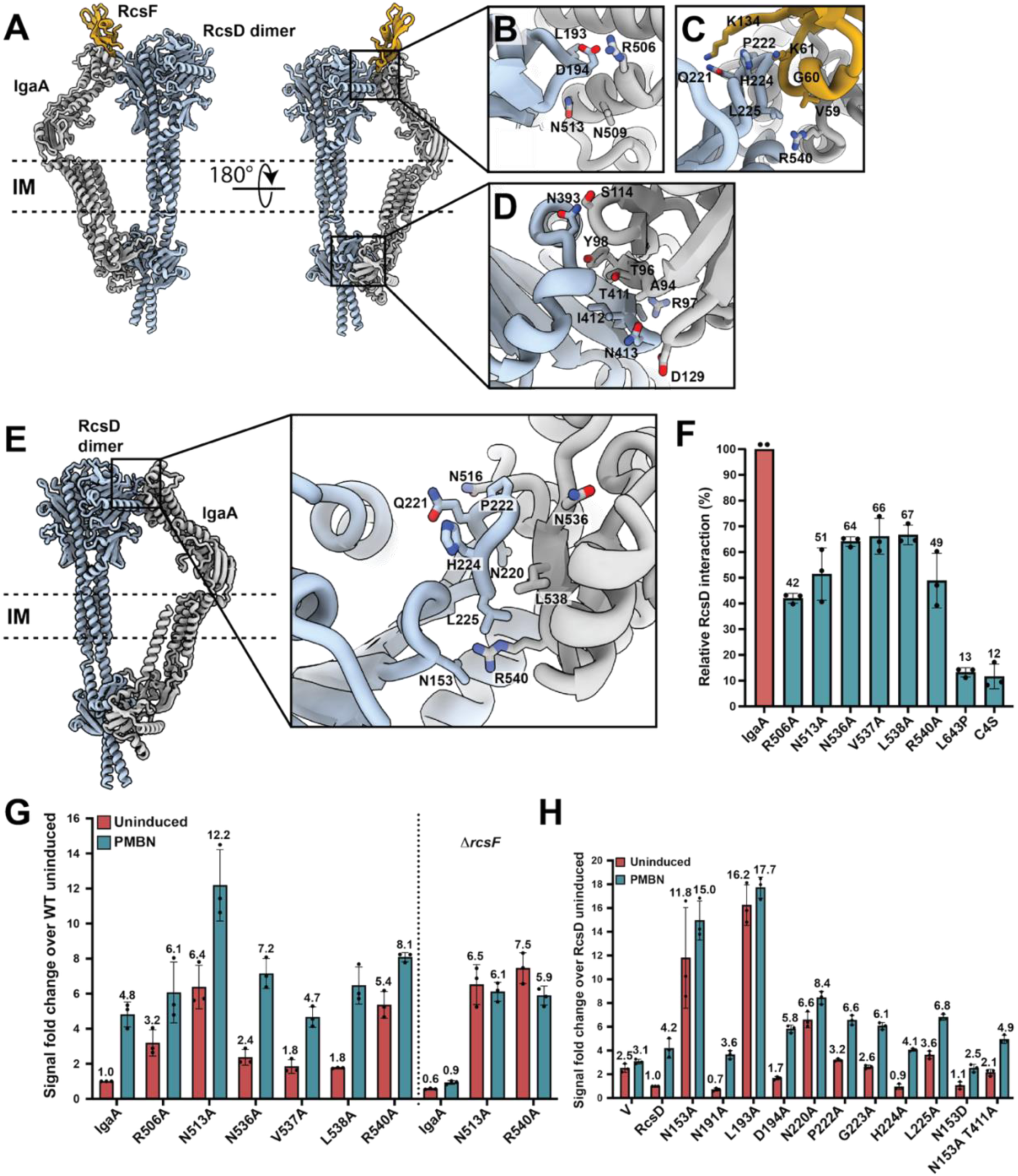
Analysis of RcsF/IgaA/RcsD interactions. A) AlphaFold3 model of *E.coli* RcsF/IgaA/RcsD_1-461_ complex. B) Detail of IgaA/RcsD binding interface at the periplasmic end. Interacting residues R506, N509, N513 are shown. C) Detail of contacts between RcsD, RcsF, and IgaA. D) Detail of cytoplasmic contact between IgaA/RcsD. E) AlphaFold3 model of *E.coli* IgaA/RcsD_1-461_ complex, including detailed view of IgaA/RcsD binding interface at the periplasmic end. F) Interaction of IgaA periplasmic mutants with RcsD. The interaction of RcsD was tested with IgaA and IgaA point mutants using BACTH in BTH101. All interactions are plotted relative to RcsD-IgaA interaction (set to 100). G) Signaling by RcsD-binding defective IgaA mutants. The P_rprA_:mCherry fluorescence of the IgaA mutant strains was measured to assay the Rcs signaling as described in Fig 3F. RFU at OD 0.4 is plotted and uninduced IgaA is considered as 1. The right panel shows IgaA alleles in the chromosome of a Δ*rcsD* Δ*rcsF* strain transformed with pEAW11. H) Signaling by RcsD periplasmic mutants. The Δ*rcsD* strain (EAW19) was transformed with either pBAD24 vector or RcsD mutants and their fluorescence assayed as described in Fig. 4E.

In parallel, we generated the AlphaFold3 model of the IgaA/RcsD complex and found that residues N536, L538, and R540 in the IgaA beta-sheet (β5), that are normally bound by RcsF in the absence of RcsD (Fig. 3A-B), are predicted to interact with RcsD in the absence of RcsF (Fig. 6E). These models thus suggest the presence of multiple modes of interaction of IgaA and RcsD, one found both in the presence and absence of RcsF (Fig. 6A-D), the other in the absence of RcsF where we observed additional contacts between RcsD/IgaA that suggest tighter interaction in the periplasmic crest (Fig 6E).

To address whether there is a shared IgaA interface with RcsF and RcsD and whether the predicted changes in IgaA/RcsD contacts in the absence of RcsF might regulate IgaA signaling to RcsD, we tested mutants in the periplasmic region of IgaA for interactions with RcsF (Fig. S8E) and with RcsD (Fig. 6F and S16A-B) using the BACTH assay and for effects on signaling (Fig. 6G). Note that the IgaA mutants L472A, I476A, M481A, and L483, shown to possess lower affinity for RcsF (Fig. 3E) and to no longer be inducible (Fig. 3F), interacted very well with RcsD (≥90% relative to WT IgaA, Fig. S16B), and thus appear to specifically disrupt the IgaA/RcsF interface. The subset of IgaA mutants that had the largest reduction in interaction with RcsD are shown in Fig. 6F, and include both regions predicted to interact with RcsD by the AlphaFold3 models (R506A and N513A in the alpha helix and N536A, V537A, L538A and R540A in the beta sheet). Mutants L643P and C4S have previously been shown to lead to high expression of the Rcs phosphorelay and are believed to have major effects on the stability and/or folding of IgaA^4,33^.

We first consider the behavior of the mutants in the IgaA periplasmic alpha-helix (residues 506-514) involved in binding to RcsD (small loop residues 191-195) (Fig. 6B). Consistent with disruption of the IgaA/RcsD interface, the R506A and N513A mutants led to a significant increase in the basal level of signaling (Fig. 6G). However, they showed no reduction in RcsF affinity (≥94% relative to IgaA, Fig. S8E), suggesting that these residues are specific to the IgaA/RcsD interface. For the N513A mutant, the RcsF contact is likely functional, since PMBN leads to a two-fold induction in expression; in the absence of RcsF the high basal level expression of N513A is similar to *rcsF*^+^ cells but is now uninducible by PMBN (Fig. 6G). Our AlphaFold3 model predicted that RcsD residues L193 and D194 interact with these IgaA residues. Mutations in either residue show reduced interaction with IgaA (Fig. S16C). L193A is constitutively on for Rcs signaling (Fig. 6H), while D194A has an increase in basal level but is still RcsF-inducible, similar to the behavior of the IgaA alleles.

The IgaA beta-sheet residues (residues 535-540, β5) needed for tight binding to RcsD were close to the RcsF binding interface both in our Cryo-EM structure (Fig. 3) and in the AlphaFold3 model, but were predicted to be closer to RcsD when RcsF was absent (Fig. 6E). Unlike the IgaA alpha-helix residues discussed above, N536A, V537A and R540A had a modestly lower RcsF affinity (<90% relative to WT IgaA, Fig. S8E), consistent with being close to and involved in the RcsF binding interface. Mutations in these residues increased the basal level of expression, and most showed further induction with PMBN (Fig. 6G). R540A had the highest basal level of expression of this set of residues, and very modest induction with PMBN (Fig. 6G). In the AlphaFold3 model for the RcsD/IgaA complex, R540 is close to RcsD N153 (Fig. 6E). RcsD N153A reduced the interaction with IgaA, while N153D increased it (Fig. S16C), consistent with a direct interaction with R540. Further supporting this interaction, N153A, like R540A, is constitutive (Fig. 6H). N153D, on the other hand, had a very low basal level and reduced induction in the presence of PMBN (Fig. 6H). This data strongly supports the AlphaFold3 model for IgaA/RcsD and identifies IgaA R540 and RcsD N153 as critical residues for signaling by PMBN.

These results demonstrate that there are RcsF-independent effects of IgaA interacting with RcsD and RcsD-independent effects of IgaA interacting with RcsF. IgaA L483A, which had a reduced interaction with RcsF but retained interaction with RcsD (Fig. 3E, Fig. S16B), shows reduced PMBN induction (Fig. 3F), consistent with it behaving like an RcsF contact. A double mutant was constructed with L483A and L538A, a mutant in the region identified as important for RcsD interaction and repression. The double mutant L483A/L538A had reduced interaction with both RcsD and RcsF (Fig. S8E and Fig. S16B). It also had a higher basal level, as seen in L538A, but a very muted induction upon PMBN treatment (Fig. S8F), consistent with a reduced recognition of RcsF, reinforcing the idea that these two regions of IgaA act somewhat independently. These results supports the need for IgaA to interact with RcsD in the periplasm to repress signaling, and is fully consistent with previous work by us and others showing that deletion of the periplasmic domain of IgaA leads to run-away (lethal) induction of Rcs^4^.

### RcsD has highly conserved residues for IgaA interaction in its periplasm and its cytoplasmic PAS-like domain

We have previously found that a mutation in the cytoplasmic PAS-like domain of RcsD-T411A (in *E.coli*), tightened interactions with the IgaA cytoplasmic region and dulled PMBN induction, suggesting that this cytoplasmic contact was a critical part of the switch regulating Rcs induction^4^. Interestingly, while RcsD-T411A also interfered with induction in a *rcsD* Δ*peri* strain (Fig. 7A), it was unable to block induction in cells carrying *igaA* deleted for the periplasmic region^4^, suggesting an asymmetry in these periplasmic interactions.

**Figure 7:**
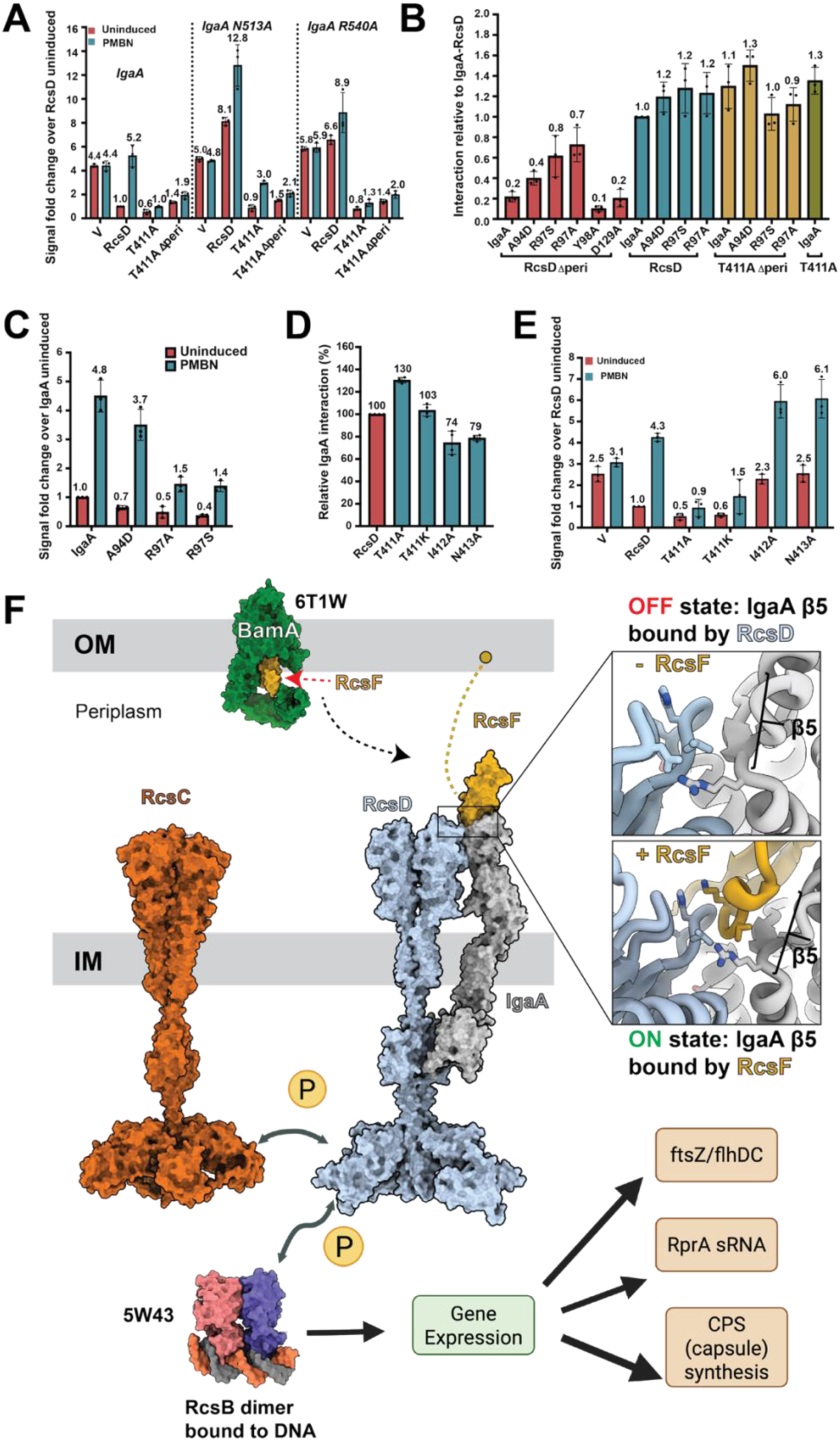
Analysis of IgaA/RcsD interactions. A) RcsD T411A can repress signaling by RcsD-binding defective IgaA mutants. The IgaA alleles are in the chromosome in a Δ*rcsD* strain as in 3F. These strains were transformed with either pBAD24 vector, *rcsD* (pEAW11), RcsD T411A (pEAW11T), or T411A del peri (pAP1102) and their fluorescence assayed as earlier. T411A lowers signaling by tightening cytoplasmic interactions. B) Interaction of IgaA cyt1 mutants with RcsD. Interaction of IgaA and its cyt1 mutants with RcsD, RcsD del peri, RcsD T411A del peri was tested using BACTH. C) Signaling by IgaA cytoplasmic mutant strains. The fluorescence of IgaA mutant strains was assayed as described in 3F. D) Interaction of RcsD cytoplasmic mutants with IgaA. The interaction of RcsD and its mutants was tested with IgaA using BACTH in BTH101. E) Signaling by RcsD cytoplasmic mutants with altered IgaA interaction. The Δ*rcsD* strain (EAW19) was transformed with either pBAD24 vector or RcsD mutants and their fluorescence assayed as in Fig. 4E. F) Updated model for Rcs phosphorelay signaling cascade. RcsF and OMPs such as BamA, sense disruptions in the outer membrane or peptidoglycan layer. The inner membrane protein IgaA receives the stress cues from RcsF, which weakens the repressive interaction between IgaA and RcsD. The signal is then transmitted to RcsC/RcsD, leading to the autophosphorylation of RcsC. RcsC then relays the phosphate to RcsD. The phosphate is then transferred to RcsB, resulting in activation of gene expression (see discussion for more details). The cytoplasmic domains of RcsD and RcsC were modeled using AlphaFold3.

We examined the effect of RcsD-T411A in two mutations, one in each of the two *igaA* regions that the AlphaFold3 modeling and BACTH suggest interact with RcsD (Fig. 6B-C, 6G) but retain interactions with RcsF: N513A in the alpha-helix and R540A in the beta-sheet (Fig. 6A-C, 6G). Mutations in either residue leads to high level expression of the phosphorelay (N513A and R540A, Fig. 7A). We expressed RcsD-T411A and RcsD-T411A Δperi in the *igaA* N513A and R540A *ΔrcsD* strains; both T411A and T411A Δperi lower the fluorescence in the context of N513A and R540A background, suggesting a tight cytoplasmic interaction is enough to block Rcs activation (Fig. 7A). For *igaA* N513A, PMBN is capable of inducing signaling 3-fold even in the T411A context, consistent with a continued interaction of IgaA with RcsF. This induction is significantly reduced by T411A Δperi, reinforcing the idea that PMBN induction acts initially through the distinct periplasmic contacts of these three proteins. R540A was less inducible, suggesting that this allele blocks the RcsF-dependent sensing, consistent with predictions of the AlphaFold3 model for this region of IgaA interacting with both RcsF and RcsD (Fig. 6A-C).

The behavior of RcsD T411A led us to previously propose an interaction critical for signaling of the RcsD PAS-like domain (containing T411) and the IgaA cyt1 domain^4^ (shown in Fig. 6D). Here we examined the PAS-like domain of RcsD and the cyt1 domain of IgaA to further understand the nature of this signaling downstream from the periplasmic contacts discussed above.

Previous BACTH assays have shown that most of the RcsD/IgaA interaction is due to the periplasmic contact^4^. Deletion of the periplasmic domain of RcsD reduced the IgaA interaction five-fold compared to WT RcsD, but the interaction of RcsD-T411A Δperi was very similar to WT RcsD (Fig. 7B), supporting the tighter interaction due to T411A. We used this property to screen for critical IgaA cytoplasmic regions that could also lead to tighter interactions with RcsD, suppressing the low BACTH interaction of IgaA with RcsD Δperi. We carried out a random mutagenesis of the IgaA cyt1 and cyt2 domains and screened this library to identify IgaA mutants with increased interactions with RcsD Δperi. IgaA cyt1 domain mutants A94D, R97S and R97A, show increased affinity for RcsD Δperi (Fig. 7B). The AlphaFold3 structure (Fig. 6D) suggested that IgaA R97 is in the vicinity of RcsD T411 and IgaA A94 is near RcsD N413/N414; thus, these mutants provide some confirmation for this structure, and are fully consistent with our previous conclusion that IgaA cyt1 is likely to be the site of interaction with the T411 region of RcsD^4^. All three alleles also showed somewhat increased interaction with WT RcsD (Fig. 7B) and A94D showed a modest increase even in the background of RcsD-T411A Δperi. These IgaA mutants were introduced into the chromosome of a Δ*rcsD* strain. When complemented with *rcsD*^+^ (Fig. 7C) they behaved similarly to RcsD-T411A, with a lower basal signal and significantly decreased PMBN induction, consistent with the idea that IgaA A94 and R97 are part of the switch that changes the IgaA/RcsD interaction upon induction.

Mutations in the PAS-like domain of RcsD near T411 had a modest decrease in IgaA interaction (Fig. 7D, S16D). These mutants were also tested for activity. T411K, like T411A, had a lower basal signal and a reduced PMBN response than RcsD (Fig. 7E). The remaining mutants tested, A410G, I412A, N413A, N414A, and E415A, all had a higher basal signals (1.7-2.7-fold higher than RcsD) and a slightly higher PMBN signal (6.2-6.7 vs 5.0 for RcsD) (Fig. S16E). This is consistent with the RcsD-PAS domain playing both positive and negative roles in regulation of Rcs induction, presumably at least in part by its interactions with the IgaA cytoplasmic domains.

## DISCUSSION

The Rcs phosphorelay is an evolved complex example of a classic two-component signal system (Fig. 1B), in which activation is dependent on the RcsF outer membrane lipoprotein and its interactions with IgaA. IgaA in turn transmits the signal by its interactions with the phosphotransferase RcsD, a protein with a similar domain architecture to its histidine kinase RcsC. Clear understanding of the mechanism by which stress signals are received and how the phosphorelay signal is initiated to change gene expression in this system have been lacking. In this study, we provide structural insights into several key components (IgaA, IgaA/RcsF, RcsD, and RcsC) using Cryo-EM, AlphaFold modeling, and structure-guided *in vivo* mutational analysis. Together our studies provide new understanding of the Rcs signal transduction mechanism in *K. pneumoniae* and *E. coli*.

### Structural basis for IgaA recognition of RcsF

Our Cryo-EM structures of the IgaA/RcsF complex (from *K. pneumoniae*) shed light on how outer membrane stress signals are transmitted across the bacterial envelope (Fig. 2-3). RcsF binds to the outer membrane-facing side of the periplasmic domain of IgaA, positioned to effectively intercept signals from the exterior (Fig. 2). Notably, IgaA residues L472, M481, I476, L483, and RcsF residues A55, L58, and F63 stabilize the binding between the two proteins. Mutations in these residues abolish binding and severely disrupt Rcs activation (Fig. 3). Our Cryo-EM studies also unveil the first full-length structures of apo IgaA and the IgaA/RcsF complex. Although the structures are limited in high-resolution details, we were able to build full-length models showing that no major conformational changes occur in IgaA upon RcsF binding. This raises an intriguing question: how is the stress signal transmitted from RcsF to IgaA, then to RcsD and RcsC?

### Insights into RcsF-dependent Rcs activation

RcsF is thought to reside in the lumen of OMPs under normal growth conditions, and recent studies have suggested that OMPs play an active role in the Rcs polymyxin response, with identical regions in RcsF contacting both OMPs and IgaA^32^. The structures here, determined in the absence of the OMPs, reveal a strong interaction between RcsF and IgaA. This interaction is essential for Rcs activation by the RcsF-dependent signal PMBN; mutations in either IgaA or RcsF that disrupt this interface lead to phenotypes akin to an RcsF-null strain (Fig. 3). Our Cryo-EM complex structures agree with a recently reported crystal structure of the *E. coli* IgaA periplasmic region in complex with RcsF^26^. Both studies found no observable changes in the structure of IgaA upon interaction with RcsF.

How does RcsF binding to IgaA affect its interaction with RcsD? Our results here and in previous works support interactions of RcsD with IgaA both in the periplasm and in the cytoplasm, with the periplasmic contacts critical for keeping tight control (repression) of signaling whereas the cytoplasmic contacts are particularly key for signaling^4^. These interactions can be detected in the absence of RcsF and while deletion of *igaA* is toxic, due to overactivation of the Rcs system, deletion of *rcsF* is not. Thus, RcsF is not needed for IgaA to repress the phosphorelay. We can therefore conclude that RcsF activation does not totally disrupt IgaA repression of the Rcs phosphorelay. Hyperinduction of Rcs by inner membrane anchored RcsF is not sufficient to cause toxic activation, suggesting that IgaA is still able to provide some repression of RcsD even under this condition.

The structural analysis of RcsF/IgaA, AlphaFold structures for IgaA interactions with RcsF and RcsD (Fig. 6), combined with our mutational analysis, allows us to propose a model in which IgaA has two sets of important but functionally different bipartite interactions in the periplasm, one specifically with RcsF and one specifically with RcsD. The second site of IgaA interaction is shared with RcsF and RcsD, but at different times – this contact acts to signal the state of induction and initiate signaling to the phosphorelay (Fig. 7F). Thus, RcsF partially disrupts the IgaA/RcsD periplasmic interaction and favorably binds to IgaA (Fig. 6-7). We propose that this process weakens the repressive contact between IgaA/RcsD, ultimately lifting the default negative regulation IgaA exerts on the phosphorelay. This model is supported by mutations with three distinct properties (Fig. 7F).

The first set of mutations are in residues at the IgaA periplasmic crest that interacts with RcsF, including L472, I476, M481 and L483, and define the RcsF-specific interaction with IgaA. Mutations in these residues are generally uninducible, but do not significantly perturb interactions with RcsD (Fig. 3E, F, and S16B). This interaction is clearly visible in the high resolution structure of IgaA and RcsF (Fig. 3B), and mutations in RcsF interacting residues have similar properties (Fig. 3C, D).

The second set of mutations retain full interaction with RcsF but disrupt the interaction of the IgaA periplasmic region with RcsD, defining a unique RcsD/IgaA anchoring interface. These reside in the periplasmic alpha-helix of IgaA and include residues R506 and N513. Mutations in these sites lead to high basal signal from the phosphorelay, consistent with repression requiring this periplasmic contact, but retain some inducibility by PMBN (Fig. 6G). This inducibility is RcsF-dependent (Fig. 6G). Therefore, these residues represent an interaction with RcsD that is independent of RcsF. We define this periplasmic contact as an important anchoring interaction, protecting the cell from runaway (and toxic) Rcs induction when RcsF perturbs the periplasmic switch interaction (Fig. 7F). Mutations in the AlphaFold-predicted RcsD region that interacts with this anchoring region have similar properties (Fig. 6B, 6H), also leading to high but inducible activity.

The third set of mutations in the IgaA periplasm reside in the beta-strand (β5) (Fig. 6E) and define the RcsF-sensitive switch interaction with RcsD (Fig. 7F). Mutations in these residues, including N536, R540 and the spanning region, reduce interaction with both RcsF and RcsD (Fig. 6F, Fig. S8E). In theory, such mutations could identify residues shared by both RcsF and RcsD in a heteromeric complex. However, in the structure determined for RcsF and IgaA, these residues (including N536 and R540) are close to RcsF, and this is also seen in the AlphaFold models of the three proteins (Fig. 6C). In contrast, in the absence of RcsF, the AlphaFold model suggests that this region is close to RcsD (Fig. 6E). Thus, we suggest that this region is the RcsF-sensitive switch, repressing RcsD under non-inducing conditions but engaged by RcsF in the presence of an inducing signal. The RcsF interaction then frees RcsD to switch on signaling. That transition to the ON state is transmitted through the periplasmic contacts to the cytoplasmic switch region; it is here that the interactions of IgaA, RcsD and RcsC lead to induction of the phosphorelay. Consistent with this switching role, mutations in this region (such as R540A) have high expression of the phosphorelay, but unlike N513, are blind to RcsF (Fig. 6). Mutations in the region in RcsD close to this IgaA switch region also strongly support the AlphaFold prediction and the role of this model; N153A leads to constitutive and poorly inducible expression of the phosphorelay while N153D, which may stabilize interactions with R540, has low and uninducible expression (Fig. 6H).

This model does not require the existence of the heterotrimeric complex of RcsF/IgaA/RcsD shown in Fig. 6, but does not rule it out, and certainly the existence of regions within IgaA that can independently interact with RcsD or RcsF support the possibility of such a complex. Two independent sets of contacts between IgaA and RcsD may explain why most of our mutations in the IgaA periplasmic domain (or the RcsD periplasmic domain) did not dramatically disrupt the interactions of these two proteins in BACTH assays (Fig. 6F, S16A), while deletion of the periplasmic region had a much stronger effect. This model predicts that mutations that disrupt both regions will be synergistic in interactions (as seen with the L483A L538A double mutant). Additionally, it is possible that mutations in another IgaA beta-sheet region (containing L643) may affect IgaA flexibility/conformation thereby impacting its RcsD interaction and signal transduction.

### Roles of the cytoplasmic switch regions

It is striking that mutations in either the anchor or periplasmic switch region lead to high level expression of the Rcs phosphorelay but can be overcome by tightening the cytoplasmic switch region with an RcsD-T411A mutation, demonstrating that the state of the cytoplasmic switch is epistatic to loosening either contact in the periplasm. However, disrupting both with a Δ*peri igaA* mutation cannot be rescued by RcsD-T411A^4^. Here, we identified IgaA residues in the first cytoplasmic region (cyt1) that could also tighten the interaction, blocking induction; this region is predicted by our AlphaFold3 model to interact with the T411 region of RcsD. Thus, it seems clear that a contact between IgaA cyt1 and the RcsD-PAS domain responds to changes in the periplasmic contacts, leading to induction. Consistent with a repressive effect of the interaction here, loosening this contact via other mutations in this region of RcsD (Fig. 7E) led to an increase in the basal level of induction. A study by the Collet lab^39^ overexpressing domains of IgaA found partial repression of IgaA lethality and high signaling when cyt1 alone was expressed, consistent with the epistasis we see for the T411A mutation. However, in their experiments, full repression required overexpression of both the cytoplasmic domains and the periplasmic domain (on separate plasmids) consistent with our finding of repressive contacts in both the cytoplasm and periplasm. The communication between the periplasmic and cytoplasmic contacts remains a subject for future analysis.

### Insights into the role of RcsC and RcsD dimerization in Rcs signaling

Most membrane histidine kinases are present as dimers and the state of dimerization dictates their behavior (reviewed in^40,41^). The Cryo-EM structures of RcsC and RcsD presented in this study represent the first transmembrane structures of any Rcs phosphorelay. We find that the TM helices dimerize differently in RcsC and RcsD. In RcsD, the TM helices form a four-helix bundle while in RcsC, the TM helices are arranged in a sheet-like architecture (are all in the same plane). Our current model for signal transduction suggests that RcsD, via communication with IgaA, regulates the kinase/phosphatase balance of RcsC. Because it is unusual for a phosphotransfer protein to carry the extra domains found in RcsD, we considered the possibility that RcsD and RcsC might heterodimerize. However, the transmembrane arrangements of our RcsD and RcsC Cryo-EM structures suggest that this is unlikely; rather, we would predict that heterotetramers (containing one dimer each of RcsD and RcsC) are assembled during phosphorelay signaling.

RcsC is also unusual among membrane-localized histidine kinases in that in many cases signals are sensed by the periplasmic domain. For RcsC, we have previously shown that the periplasmic domain is dispensable for RcsF-dependent signaling, although we have identified at least one pathway for Rcs activation via overproduction of the DrpB protein, that depends upon the RcsC periplasmic regions^33^. Previous tests on RcsC chimeras also implicated the TM regions and periplasmic loop in regulation of biofilm production^29^. Movement of TM helices have been shown to transduce conformational changes in periplasmic sensor domains to cytoplasmic PAS or receiver domains. It seems feasible that a “classic” histidine kinase has evolved to allow new signals and regulation via RcsF, IgaA and RcsD. In this evolved signaling system, the critical communication between RcsD and RcsC must take place in the cytoplasm. However, the cytoplasmic domains are not visible in our structures of these proteins, leaving important questions for future studies.

### Universal model for regulation of Rcs phosphorelay cascade

Our Cryo-EM structures of the Rcs proteins, together with structure-guided genetic manipulations, provide new insights into the functional roles of key components within the Rcs phosphorelay system. We and others have demonstrated that, due to their sequence and structural similarity, these proteins utilize similar signaling mechanisms across most enterobacteria^3,19^. Our study enables us to propose a universal model for how Rcs signaling is regulated in enterobacteria (Fig. 7F). This model not only integrates but also refines previously proposed partial models that describe the dynamics of envelope stress by RcsF and signal transduction to downstream components^4,23,27,32,42–44^. In the absence of stress, RcsF remains sequestered, unavailable to fully interact with IgaA. This allows IgaA and RcsD to maintain a tight bipartite interaction, and the Rcs phosphorelay remains in the OFF state. However, when the bacteria encounter stress, OMPs and RcsF sense the disruption, leading to the release of RcsF to interact with IgaA, disrupting the switch contacts between IgaA and RcsD. This leaves RcsD able to activate the RcsC phosphorelay, switching RcsC from primarily a phosphatase to a kinase, by interactions still to be defined. RcsC both autophosphorylates and transfers the phosphate to RcsD. Finally, the phosphate is received by the transcriptional regulator RcsB, which then binds to promoter DNA and effectively alters gene expression to respond to stress.

Our study uncovers the mechanistic and structural insights into how the Rcs phosphorelay is activated from a “repressed” state during envelope stress. This activation enables bacteria to rapidly and reversibly respond to various perturbations, thereby regulating essential behaviors that are critical for virulence. As the critical interactions we identified in our study are well conserved, our structures provide a valuable tool for structure-based design of novel small molecule ligands against Rcs proteins in multi-drug resistant bacteria like *Klebsiella*.

## DATA AVAILABILITY

Cryo-EM maps have been deposited at Electron Microscopy Data Bank (EMDB) under accession codes EMDB-75234 (IgaA_Kp_/RcsF_Kp_), EMDB-75235 (IgaA_Kp_/Fab57), EMDB-75236 (Full-length IgaA_Kp_/Fab57), EMDB-75238 (Full-length IgaA_Kp_-Fab57/RcsF_Kp_), EMDB-75237 (IgaA_Kp_-Fab57/RcsF_Kp_), EMDB-75239 (RcsD_Kp_), EMDB-75240 (RcsD_Ec_), EMDB-75241 (RcsC_Kp_), and EMDB-75242 (RcsC_Ec_). The corresponding atomic coordinates have been deposited to Protein Data Bank (PDB) using accession codes 10KG (IgaA_Kp_/Fab57), 10KH (IgaA_Kp_-Fab57/RcsF_Kp_), 10KI (RcsD_Kp_), 10KJ (RcsD_Ec_), 10KK (RcsC_Kp_), and 10KL (RcsC_Ec_).

## ACKNOWLEGEMENTS

We thank Yan Li, Anneliese Faustino, and Stephen Fried for their help in conducting mass spec experiments in the early stages of this study. We also thank Ulrich Baxa, Yanxiang Cui, Huabin Wang, Rick Huang, and Chih-Ta Chien for their assistance with Cryo-EM data collection and Di Wu and Grzegorz Piszczek for their help with biophysical experiments. M.N., I.B, R. G., S.S., AA., B.M.B. and S.K.B. are supported by the Intramural Research Program of the National Institute of Diabetes and Digestive and Kidney Diseases, National Institutes of Health. M.N. is also supported by Nancy Nossal fellowship and the NIGMS Postdoctoral Research Associate Training (PRAT) Program. A.P., N.M., and S.G. are supported by the Intramural Research Program of the National Cancer Institute, National Institutes of Health. We are grateful to members of both Dr. Susan Buchanan’s and Dr. Susan Gottesman’s labs for providing helpful feedback throughout the development of this work. This work was made possible by the state-of-the-art facilities, including the NIH Multi-Institute Cryo-EM Facility (MICEF), the NIH Intramural Cryo-EM Consortium (NICE), and the Biophysics Core Facility (NHLBI), as well as the National Institutes of Health high-performance computing cluster (NIH HPC), Biowulf. This research was supported by the Intramural Research Program of the National Institute of Diabetes and Digestive and Kidney Diseases (NIDDK) within the National Institutes of Health (NIH). The contributions of the NIH authors are considered Works of the United States Government. The findings and conclusions presented in this paper are those of the authors and do not necessarily reflect the views of the NIH or the U.S. Department of Health and Human Services.

## MATERIALS AND METHODS

### Bacterial growth conditions

*E. coli* strains were grown in LB medium with appropriate antibiotics (ampicillin 100 μg/ml, kanamycin 30-50 μg/ml, chloramphenicol 10 μg/ml for *cat-sacB* allele and 25 μg/ml for plasmids, tetracycline 25μg/ml). All cultures were grown in 37°C, unless stated otherwise. Glucose (1%) was added to the medium to reduce the basal expression from the plasmids containing pBAD and pLac promoters.

Strains, plasmids, primers, and synthetic DNA fragments (gBlocks) used in this study are listed in Supplementary Tables S3, S4, S5 and S6 respectively. Oligonucleotides and gBlocks were from IDT DNA, Coralville, IA.

### DNA mutagenesis and strain generation

Plasmids were generated by Gibson assembly method using the In-fusion HD Cloning kit (Takara Bio USA)^45^. PCR products were purified using column purification (Qiagen) and transformed into NEB DH5a F’*lacIQ* cells. Site-directed mutagenesis in the genes was carried out using the QuikChange Site-directed mutagenesis kit (Agilent). Sequencing was done to confirm all chromosomal and plasmid modifications. Strains were constructed by recombineering or P1 transduction with selectable markers (Table S3). Recombineering was done in strains carrying a chromosomal mini-λ Red system (*miniλ::tet*) or a plasmid-borne Red system (pSIM27/pSIM6). Some strains were generated by direct P1 transduction from the corresponding mutant strains in the Keio collection^46^ and their antibiotic resistance cassette removed using plasmid pCP20 (Cherepanov et al.,1995).

### Fluorescence reporter assays

The strains used for the growth and fluorescence assays carried a P_rprA_::mCherry transcriptional fusion at the *ara* locus as a reporter for Rcs signaling^4^. They were grown overnight at 37°C in MOPS minimal 0.2% glucose medium containing the necessary antibiotics. Subsequently, they were diluted to OD_600_ 0.04-0.05 in the MOPS minimal glucose medium and transferred to 96-well plates. Their optical density (OD_600_) and mCherry fluorescence were monitored every fifteen minutes for 12 hours at 37°C in a Tecan Spark microplate reader. Polymyxin B nonapeptide (PMBN; Sigma) was added to the medium in the beginning at a final concentration of 20 μg/ml to induce the Rcs system. For cells expressing RcsF-T25 constructs (Fig. 3D), overnight cultures were grown in MOPS minimal glucose, washed, and then diluted into fresh MOPS minimal glycerol media (0.05% glucose, 0.5% glycerol) with or without 0.5 mM IPTG. Each assay was performed in technical duplicates in the microtiter plate, with the biological replicates performed on different days. The relative fluorescence values (RFU) at equivalent OD_600_ values (0.4 ± 0.03) for each strain was used to plot the bar graphs using GraphPad Prism 10 software. Additionally, these values were depicted as signal fold change of fluorescence signal over an uninduced wild type control. Error bars represent the standard deviation of at least 3 assays. Values derived from at least three independent experiments were plotted to show the mean with error bars indicating standard deviation.

### Bacterial adenylate cyclase two hybrid assay (BACTH)

An adenylate cyclase mutant strain (BTH101) or BTH101 Δ*rcsD*, where indicated, was used to assay protein-protein interactions by BACTH^30,31^. The two proteins whose interactions were being measured were fused to either the T18 or the T25 portion of adenylate cyclase protein. The interaction of the two proteins leads to the reconstitution of adenylate cyclase allowing the cAMP synthesis necessary for activating the *lac* operon via its CRP interaction. This leads to beta-galactosidase activity which is assayed as a measure of protein-protein interaction. There is no beta-galactosidase activity if the two proteins do not interact. IgaA and RcsD were fused at their C-terminal to the Cya fragments^4^. RcsF was cloned with a MalF transmembrane helix and fused at its N-terminal to the T25 Cya fragment. Plasmids expressing IgaA, RcsD, or RcsF as T18/T25 fusion constructs were co-transformed into BTH101 strain, plated on LB-agar medium containing 100 μg/ml ampicillin and 50 μg/ml kanamycin and incubated at 30°C for 2 days. The resulting colonies were then inoculated and grown overnight in LB medium containing 100 μg/ml ampicillin, 50 μg/ml kanamycin, and 0.5 mM IPTG at 30°C. These cultures were used to carry out the beta-galactosidase assay in 96-well plates. For this assay, the degradation of ONPG was monitored at the OD 420 nm at 28°C for 30 min at intervals of 1 min in a Tecan Spark microplate reader. The beta-galactosidase activity was based on the kinetics of ONPG degradation and calculated using the slope of OD 420 divided by the OD 600 of the culture. The protein fusions paired with their cognate vector acted as negative controls and showed very low activity. The beta-galactosidase activity is plotted in arbitrary units or relative to a IgaA/RcsD/RcsF interaction, set to 1. Every graph is compiled from at least 3 separate sets of assays.

### Western Blotting

For Fig. S12G, the *E. coli* DH5-alpha F’IQ strains were transformed with the RcsD Δperi constructs. The colonies were inoculated in LB medium containing ampicillin (100 μg/ml) and the cultures grown to OD 0.4 at 37°C after which 0.5 mM IPTG was added for induction. The cultures were allowed to grow for 4 hours and then a 1 ml aliquot was collected and used for TCA precipitation. They were washed with acetone, resuspended in sample buffer normalized according to their OD. Equal volumes were run on parallel 4-12% NuPage gradient gels (Invitrogen, CA). The transfer for western blotting was done in an iBlot2 onto nitrocellulose membrane as per manufacturer’s protocol (Invitrogen). After blocking in 5% BSA, membranes were incubated with the primary antibody, washed, incubated with an anti-mouse DyLight800 fluorescent secondary antibody (1:10000 dilution; Biorad, CA). The primary antibodies used were mouse anti-CyaA (1:10000 dilution; SantaCruz Biotechnology) and mouse anti-Ef-Tu (1:10000 dilution; Hycult Biotech). Imaging was done on a ChemiDoc MP system (Biorad).

### Protein expression and purification

#### Purification of IgaA

The gene encoding IgaA_Kp_ was cloned into pBAD28 vector (Addgene) harboring 10x his-tag and Tobacco Etch Virus (TEV) cleavage site at the C-terminus codon optimized for protein expression in *E. coli* (ThermoFisher). *E. coli* OverExpress^TM^ C41(DE3) cells were transformed with the expression plasmid, plated on LB agar and incubated overnight at 37 °C. Fresh colonies were picked and used to inoculate 1 ml of LB broth, grown for 30-60 minutes, and subsequently transferred to 100 ml of LB broth starter culture. The 100 ml culture was grown at 37 °C until an OD600 of 0.4, after which 12 ml was transferred into 1 L of 2x YT (3-12 liters per prep). Protein expression was induced with 0.1% L-arabinose (Sigma-Aldrich) at an OD of 0.6. After 16 hours of induction at 25 °C, cells were harvested by centrifugation at 5,000 × g for 12 minutes. Cell pellets were resuspended in 50 ml of lysis buffer (20 mM Sodium Phosphate buffer pH 7.6, 1 M NaCl, 0.5 mM TCEP) and a combination of 2-4 protease inhibitor tablets, 1 mM AEBSF (ThermoFisher), and 1 µl/volume of 10 mg/ml DNase I (NEB). Cells were lysed using emulsiflex (Avestin), cell debris was clarified by centrifugation at 10,000 × g for 10 minutes, and membranes were pelleted by spinning at 45,000 rpm for 45 minutes using Ti45 rotor in ultracentrifuge. Pelleted membranes were resuspended in solubilization buffer (1x PBS at pH 7.4, 10% glycerol, 10 mM BME, 4 ULTRA protease inhibitor tables, 0.5 mM EDTA, 200 mM NaCl, and 50 mM imidazole) and 1% LMNG added dropwise manner slowly. After 3 hours of solubilization at 4 °C (or overnight), another high-speed spin was performed in ultracentrifuge to isolate the LMNG solubilized protein. The supernatant was then applied to a 5 ml HisTrap HP column (Cytiva) pre-equilibrated with binding buffer (1x PBS at pH 7.4, 2% glycerol, 10 mM BME, 0.5 mM EDTA, 200 mM NaCl, and 50 mM imidazole, 0.002% LMNG) then washed and eluted with a buffer containing 300 mM imidazole following gradient elution protocol. Eluted fractions were pooled, concentrated, and further purified by size exclusion chromatography using a Superose 6 Increase 10/300 GL column (Cytiva) equilibrated with 20 mM HEPES pH 7.5, 300 mM NaCl, 2% glycerol, 1 mM TCEP, 0.01% sodium azide, and 0.002% LMNG. When necessary, TEV site cleavage was performed at 1:100 IgaA/protease ratio in 50 mM Tris-HCl pH 7.5, 300 mM NaCl, 2% glycerol, 1 mM TCEP, 0.01% sodium azide, and 0.002% LMNG, 50 mM imidazole. IgaA from other species were purified in a similar manner.

IgaA was also stably purified in amphipols, specifically NAPol (Anatrace). Initially, IgaA was purified in n-dodecyl-β-D-maltoside (DDM) using a procedure similar to that was used for LMNG. NAPol was resuspended in reconstitution buffer (20 mM Tris pH 7.4, 1 mM TCEP, 0.01% sodium azide, 200 mM NaCl, and 1% glycerol) and added to IgaA at a concentration of 1 mg IgaA per 4 mg NAPol, in the presence of 0.2 mM *E.coli* polar lipids. The mixture was incubated at 4 °C with rocking for 90 minutes. Pre-washed Bio-beads SM-2 adsorbent media (Bio-Rad) were added at 0.1 g per 2 ml reaction to remove excess detergent, and incubated overnight at 4 °C with rocking. The beads were then separated from the protein using a Steri-flip, and NAPol-IgaA was centrifuged at 50,000 rpm for 20 minutes using a TLA 55 rotor in ultracentrifuge to clarify any aggregates. Finally, the sample was purified on Superose 6 Increase 10/300 GL column (Cytiva), equilibrated with 20 mM Tris-HCl pH 7.5, 200 mM NaCl, 1 mM TCEP, and 0.01% sodium azide.

#### Purification of RcsD

The expression of full-length RcsD_Kp_ containing a C-terminal 10xHis and TEV site in pBAD28 vector (ATCC) was generated codon optimized for protein expression in *E. coli* (ThermoFisher Scientific). *E. coli* OverExpress^TM^ C41(DE3) or OverExpress^TM^ C43(DE3) cells (Sigma-Aldrich) were transformed with the expression plasmid, plated on LB agar and incubated overnight at 37°C. Fresh colonies were picked and used to inoculate 1 ml of LB broth, grown for 30-60 minutes, and subsequently transferred to 100 ml of LB broth starter culture. The 100 ml culture was grown at 37 °C until an OD600 of 0.4, after which 12 ml was transferred into 1 L of 2x YT media (2-4 liters per prep). Protein expression was induced with 0.1% L-arabinose at an OD of 0.6. After 16 hours of induction at 25 °C, cells were harvested by centrifugation at 5,000 × g for 12 minutes. In some cases, short induction times such as 2-4 hours yielded similar results. Cell pellets were resuspended in 50 ml of lysis buffer (20 mM Sodium Phosphate buffer pH 7.6, 1 M NaCl, 0.5 mM TCEP, 2 mM MgCl_2_) and a combination of 2 cOmplete^TM^ EDTA-free protease inhibitor tablets (Sigma), 1 mM AEBSF, and 1 µl/volume of 10 mg/ml DNase I (NEB). Cells were lysed using emulsiflex (Avestin), cell debris was clarified by centrifugation at 10,000 × g for 10 minutes, and membranes were pelleted by spinning at 45,000 rpm for 45 minutes using Ti45 rotor in ultracentrifuge. Pelleted membranes were resuspended in solubilization buffer (1x PBS at pH 7.4, 10% glycerol, 10 mM BME, 4 ULTRA protease inhibitor tables, 0.5 mM EDTA, and 50 mM imidazole) and 1% LMNG added slowly in dropwise manner. After 3 hours of solubilization at 4 °C (or overnight), another high-speed spin was performed in ultracentrifuge to isolate the LMNG solubilized protein. The supernatant was then applied to a 5 ml HisTrap HP column (Cytiva) pre-equilibrated with binding buffer (1x PBS at pH 7.4, 2% glycerol, 10 mM BME, 0.5 mM EDTA, 200 mM NaCl, and 50 mM imidazole, 0.002% LMNG) then washed and eluted with a buffer containing 300 mM imidazole following gradient elution protocol. The addition of NaCl and 50 mM imidazole are critical for preventing contaminants that co-purify with RcsD. Eluted fractions were pooled, concentrated, and further purified by size exclusion chromatography using a Superose 6 Increase 10/300 GL column (Cytiva) equilibrated with 20 mM Tris pH 7.5, 300 mM NaCl, 2% glycerol, 1 mM TCEP, 0.01% sodium azide, and 0.002% LMNG. This step is important to ensure the aggregates are separated from dimer species. When necessary, TEV site cleavage was performed at 1:100 RcsD/protease ratio in 50 mM Tris-HCl pH 7.5, 300 mM NaCl, 2% glycerol, 1 mM TCEP, 0.01% sodium azide, and 0.002% LMNG, 50 mM imidazole. RcsD_Ec_ was purified in a similar manner.

#### Purification of RcsC

The expression and purification of RcsC follows the protocol described for RcsD with minor modifications. RcsC_Kp_ generally has poor expression and its prone to mutation, degradation, and aggregation during an overnight induction. Therefore, cells harboring the RcsC expression plasmid were grown to 1.5-2 OD, then induced with 0.1%. L-arabinose for 1-2 hours at 25 °C. For protein purification, it’s important to use Superose 6 column to separate the many aggregate species from stable dimer peaks that elute during the purification. The full-length RcsC_Ec_ was purified in a similar manner.

#### Purification of RcsF

The gene encoding RcsF_Kp_ aa48-134 (or aa1-134) was cloned into pET28 vector (EMD Millipore) harboring 10x his-tag and TEV cleavage site at the N-terminus optimized for protein expression in *E. coli* (ThermoFisher Scientific). *Rosetta-gami2* C41(DE3) cells (NEB) were transformed with the expression plasmid, plated on LB agar and incubated overnight at 37 °C. Fresh colonies were picked and used to inoculate 1 ml of LB broth, grown for 60 minutes, and subsequently transferred to 100 ml of LB broth starter culture. Cells were then diluted to 1L media, grown at 37 °C to an OD of 0.6, and induced overnight at 20 °C with 0.5 mM IPTG. Harvested cells were lysed using Emulsiflex in 100 mM Sodium Phosphate buffer pH 7.4, 300 mM NaCl, 5% glycerol, 1 mM EDTA, and 20 mM imidazole, then clarified by centrifugation at 23000g for 45 minutes, and loaded onto 5 ml HisTrap column. Bound protein was eluted with the lysis buffer supplemented with 300 mM imidazole. When necessary, TEV site cleavage was performed at 1:100 RcsF/protease ratio in 50 mM Tris-HCl pH 7.5, 300 mM NaCl, 2% glycerol, 20 mM imidazole and repassed over HisTrap column. Before use for structural or binding studies, RcsF was purified by Superose 6 Increase 10/300 GL column equilibrated with 20 mM Tris pH 7.4, 200 mM NaCl, 5% glycerol, 0.5 mM EDTA.

### Anti-IgaA monoclonal antibody discovery and purification

The mouse monoclonal IgaA antibody was generated against the full-length *Klebsiella* IgaA purified in LMNG (by Covance/LabCorp, Denver, PA). The 9 month protocol was divided in 3 phases. In phase I, the native protein combined with Freund complete adjuvant was used to inoculate and boost 5 Balb/C mice for 87 days. In Phase II, spleen cells from hyper-immunized mice were prepared and fused with P3X63Ag8.653 or SP2/OAG14 myelomas using polyethylene glycol. Viable hybridomas were selected and screened for antigen-specific antibodies by ELISA to the native protein. Media from positive hybridomas was also tested by Western blot against the denatured protein. Since the goal was to generate a monoclonal against the native form of IgaA, we selected 3 clones that were positive in ELISAs and negative on a Western blot to use in Phase III. The 3 selected primary clones were subcloned for further expansion. Wells with growing cells following cloning were screened for antibody secretion by ELISA. The one clone that met our criteria, HL7828.813.57 (Fab57), was ultimately used in this work. A sample of HL7828.813.57 was sent to GenScript (Piscataway, NJ) for SC2188, standard antibody sequencing for variable domain and signal sequence.

Using the full-length monoclonal antibody for structural studies was not feasible in our initial attempts. We therefore used Fab purification kit supplied by ThermoFisher to isolate the Fab domain using the provided protocol (Pierce^TM^ Fab preparation kit, Cat #44985).

### Sedimentation Velocity Analytical ultracentrifugation (AUC)

Sedimentation velocity experiments were conducted at 20°C and 50,000 rpm (corresponding to 201,600 x *g* at 7.20 cm) on a Beckman Coulter ProteomeLab XL-I or Beckman Optima XL-A analytical ultracentrifuge following standard protocols^47^. Samples of purified IgaA and RcsF were obtained in 20 mM Tris-HCl pH 7.5, 200 mM NaCl, 0.5 mM TCEP, 5% (v/v) glycerol, and 0.002% LMNG. The individual proteins and their mixtures were studied in two-channel, 12 mm pathlength cells. Size exclusion chromatography fractions of purified *E. coli* and *K. pneumoniae* RcsD were obtained in 20 mM Tris-HCl pH 7.5, 200 mM NaCl, 0.5 mM TCEP, 5% (v/v) glycerol, and 0.002% LMNG. Size exclusion chromatography fractions of purified EcRcsC were obtained in 50 mM HEPES-NaOH pH 8.0, 300 mM NaCl, 1 mM TCEP, 0.01% NaN_3_, and 0.001% LMNG. Fractions of KpRcsC were obtained in phosphate buffer saline with 1 mM TCEP, 0.2 mM MgCl_2_, 0.1 mM ATP, 0.01% NaN_3_, 0.1 mM AEBSF, 5% (v/v) glycerol, and 0.02% GDN. All RcsD and RcsC fractions were analyzed in two-channel, 12 mm pathlength cells. Sedimentation velocity scans were collected using the absorbance optical system at 280 nm, and when available, the Rayleigh interference system at 655 nm. Data were analyzed in SEDFIT^48^ in terms of a continuous c(*s*) distribution of Lamm equations. In the case of polydisperse sample (*e.g.*, KpRcsC), a ls*-g(s) distribution was used to model the data. Solution densities ρ, solution viscosities η, and protein partial specific volumes of the peptides were calculated in SEDNTERP^49^. Discrete species were analyzed using the membrane protein calculator in GUSSI^50,51^ using the best-fit frictional ratios from SEDFIT.

### Isothermal Titration Calorimetry (ITC)

Isothermal titration calorimetry (ITC) experiments were performed at room temperature with a stirring speed of 750-1000 rpm using a MicroCal iTC200 microcalorimeter (Malvern). Prior to the experiments, IgaA and RcsF were dialyzed four times in a buffer containing 20 mM HEPES pH 7.5, 0.2 mM TCEP (Tris(2-carboxymethyl)phosphine hydrochloride), 200 mM NaCl, 0.01% sodium azide, and 5% glycerol. At least one of the four dialysis steps was conducted overnight. The calorimeter cell was loaded with 5 µM full-length IgaA, and titrations were performed by injecting 2 µl of 40-50 µM RcsF over 18-20 injections at 300-second intervals. The raw isotherms were integrated using origin 8 and NITPIC, then fitted and further analyzed using SEDPHAT and GUSSI to generate publication-quality figures^52–54^.

### Cryo-EM sample preparation and data collection

All Cryo-EM samples were vitrified by plunge freezing in liquid ethane using a vitrobot Mark IV (Thermo Fisher Scientific) with blot force of 5-7, blot time 4-8 seconds, 100% humidity, temperature set at 4 C, and blotted using a Whatman filter paper (Ted Pella). Grids were prepared by applying 3-4 µl of samples on glow-discharged holey UltraAUFoil grids (R1.2/1.3, 300 mesh, Electron Microscopy Services) using PELCO Easiglow (Ted Pella) for 45-60 seconds at 15 mA. In the case of RcsD, orientational bias was resolved by freezing another set of grids using a Chameleon (SPT Labtech LTD).

To prepare IgaA/RcsF complex, IgaA-NAPol and RcsF were mixed at 1:5 molar ratio and incubated for 2 hours. The sample was then purified by Superose 6 column in buffer containing 50 mM Tris-HCl (pH 7.5), 2% glycerol, 1 mM TCEP, 0.01% sodium Azide, and 300 mM NaCl. Grids were prepared at ∼ 2 mg/ml complex concentration. Cryo-EM data were collected on a 200 keV Glacios equipped with Gatan K3 direct electron detector at the NIH National Institute of Diabetes and Digestive and Kidney Diseases Cryo-EM core (NIDDK). Data was collected using SerialEM at a magnification of 45,000x in super-resolution mode. A total of 8,398 movies were acquired from two grids at a physical pixel size of 0.445 Å/pixel in counting mode with a nominal dose of 65-70 e^-^/ Å^2^ divided over 32 frames. The defocus range was set to –0.8 – −2.6 µm.

To prepare IgaA/Fab57 complex, IgaA-NAPol and Fab57 were mixed at 1:1 molar ratio and incubated for 2 hours. The sample was then purified by Superose 6 column in buffer containing 20 mM HEPES pH 7.5, 1 mM TCEP, 0.01% sodium Azide, and 200 mM NaCl. Grids were prepared at ∼ 2 mg/ml complex concentration. Cryo-EM data were collected on a 300 keV Titan Krios equipped with a Gatan K3 direct electron detector and BioQuantum Imaging filter with slit-width of 20 eV, at the NIH Multi-Institute Cryo-Electron Microscopy Facility (MICEF). Data collection was performed using SerialEM, resulting in 10,281 movies acquired from a single grid at a physical pixel size of 0.429 Å/pixel. The data were collected in counting mode with a nominal dose of 60.24 e^-^/ Å^2^ distributed over 28 frames. The defocus range was set from – 0.8 to –2.4 µm.

To prepare IgaA-Fab57/RcsF complex, we first reconstituted IgaA-LMNG/RcsF by initially mixing IgaA and RcsF at 1:5 molar ratio and incubating for 2 hours. Isolated Fab57 is then added to the IgaA/RcsF complex at 1:1 molar ratio and incubated for additional 2 hours. The sample was then purified by Superose 6 column in buffer containing 50 mM HEPES pH 7.5, 1 mM TCEP, 0.01% sodium Azide, 0.002% LMNG, and 300 mM NaCl. Grids were prepared at ∼ 1 mg/ml complex concentration. Cryo-EM data were acquired on a 300 keV Titan Krios equipped with a Gatan K3 direct electron detector and a BioQuantum imaging filter with a slit width of 20 eV at MICEF. Data collection was performed using EPU, resulting in 16,153 movies acquired from a three grids at a physical pixel size of 0.412 Å/pixel. The data were collected in counting mode with a nominal dose of 46-58 e^-^/ Å^2^ distributed over 40 frames. The defocus range was set from –0.8 to –2.8 µm.

The grids for all RcsD and RcsC samples were prepared using their respective purification buffers (SEC). For RcsD_Kp,_ grids were prepared by applying 2 mg/ml protein concentration for vitrobot and 9 mg/ml for Chameleon grids. RcsD_Ec_ grids were prepared at 1-2 mg/ml protein concentration. RcsC_Kp_ grids were prepared at 0.4 mg/ml, while RcsC_Ec_ at 1.5 – 2 mg/ml.

RcsD_Kp_ data were acquired using 300 keV Titan Krios equipped with Gatan K3 direct electron detector and BioQuantum Imaging filter with slit-width of 20 eV at the National Cancer Institute of NIH Intramural Cryo-EM Consortium (NICE). Data acquisition was performed using SerialEM at magnification of 105,000 in super-resolution mode, resulting in 10,502 movies acquired from two grids, with 6,646 movies from Vitrobot grid and 3,856 from Chameleon grid. The pixel size was 0.429 Å/pixel. Data were collected in counting mode with a nominal dose of 60 e^-^/ Å^2^ distributed over 40-50 frames. The defocus range was set from –0.8 to –2.2 µm.

RcsD_Ec_ data were acquired using a 300 keV Titan Krios equipped with Gatan K3 direct electron detector and BioQuantum Imaging filter with slit-width of 20 eV, and a 100 µm objective aperture at MICEF facility. Data acquisition was performed using SerialEM at magnification of 105,000 in super-resolution mode, resulting in 9,742 movies acquired from two grids. The pixel size was 0.412 Å/pixel. Data were collected in counting mode with a nominal dose of 59.12 e^-^/ Å^2^ distributed over 40 frames. The defocus range was set from –0.8 to –2.2 µm.

RcsC_Kp_ data were acquired using a 300 keV Titan Krios equipped with Gatan K3 direct electron detector and BioQuantum Imaging filter with slit-width of 20 eV, and a 100 µm objective aperture at NICE facility. Data acquisition was performed using SerialEM at magnification of 105,000 in super-resolution mode, resulting in 12,005 movies acquired from a single grid. The pixel size was 0.415 Å/pixel. Data were collected in counting mode with a nominal dose of 61 e^-^/ Å^2^ distributed over 40 frames. The defocus range was set from –0.8 to –2.6 µm.

RcsC_Ec_ data were acquired using a 300 keV Titan Krios equipped with Gatan K3 direct electron detector and BioQuantum Imaging filter with slit-width of 20 eV, and a 100 µm objective aperture at MICEF facility. Data acquisition was performed using EPU at magnification of 105,000 in super-resolution mode, resulting in 11,007 movies acquired from a single grid. The pixel size was 0.412 Å/pixel. Data were collected in counting mode with a nominal dose of 55.26 e^-^/ Å^2^ distributed over 40 frames. The defocus range was set from –0.8 to –2.6 µm.

### Cryo-EM image processing

For image processing, the detailed data processing pipeline for all structures is provided in supplementary figures. All Cryo-EM datasets were imported into CryoSPARC V4.6, where beam-induced motion of the exposures were corrected using patch motion correction, followed by contrast transfer function (CTF) estimation. Corrected micrographs were further curated, and those micrographs with resolutions worse than 4-5 Å, along with other outliers, were discarded.

For the IgaA/RcsF complex structure, approximately 11.4 million particles were initially picked using a blob picker from 7,619 curated exposures. These particles underwent 2D classification and particle curation. *Ab initio* models were generated from both selected and rejected particles. Iterative heterogeneous refinement was then performed with both the “good” and “bad” maps against an entire set of particles to isolate the “good” particles for initial reconstruction. Subsequently, 2D templates were generated using the good models for another round of particle picking, yielding 9 million particles. Additionally, 6.9 million particles were picked using elliptical blobs since IgaA/RcsF complex appears to be elongated and biased towards one direction. Particles from all three picking methods were combined and duplicates were removed, resulting in 14 million particles. These particles were subjected to multiple rounds of heterogeneous refinement, ab initio reconstruction, homogeneous refinement, NU refinement, and local refinement to curate and identify a set of 209K particles, which were used to generate a 5.44 Å map (as reported by CryoSPARC). Additionally, local masks around the periplasmic domains of IgaA and RcsF, as well as the TM and cytoplasmic domains of IgaA, were applied during refinement to improve local map resolution. This improved the clarity of the backbone, facilitating more accurate model fitting and building (see Fig. S2 for additional details).

For the IgaA/Fab57 structure, an initial blob picking on 7457 curated exposures resulted in selection of 13 million particles. These particles underwent several iterative rounds of reference-free 2D classification, resulting in a curated set of 4.23 million particles. This curated particle set was then used to create 2D templates and used for template picking, yielding 7.5 million particles. The particles from blob and template picking were combined, and the duplicates were removed. Several rounds of multi-class heterogeneous refinements, *ab initio* reconstruction, homogeneous refinement, NU refinement, and local refinement were performed to further refine and isolate a set of 489K high-quality particles. These particles were then subjected to a final round of reference-based motion correction and local refinement, which produced a ∼2.87 Å structure that is presented in this study. Both the sharpened and DeepEM maps were used for model building. For the full-length IgaA/Fab57 complex, a separate set of 377K particles was used to build a 5.5 Å map (see Fig. S4 for additional details).

The high-resolution structure of IgaA-Fab57/RcsF complex was achieved by using a total of 11,930 curated exposures. An initial blob picking identified 19.2 million particles, which underwent several rounds of 2D classification, multi-class *ab initio* reconstructions to generate good and bad volumes, and heterogeneous refinements to isolate a set of high-quality particles. The selected particles were then used to generate 2D templates. Template picking resulted in 13 million particles. The particles from blob picking and template picking were combined, and duplicates were removed to yield a total of 22 million particles. Several rounds of multi-class heterogeneous refinements, *ab initio* reconstruction, homogeneous refinement, NU refinement, and local refinement were performed to further refine and isolate a set of 360K high-quality particles. To further improve the resolution and overall quality of the reconstruction, multiple rounds of reference-based motion correction, along with non-uniform and local refinements using masks were applied. The final set of curated particles produced a 2.79 Å map that is presented in this study. For the full-length IgaA-Fab57/RcsF complex, a separate set of 460K particles was used to build a 2.88 Å map. Although the TM and cytoplasmic domains showed poor resolution, both CryoSPARC and Phenix are in agreement regarding the resolution estimation (see Fig. S6 for additional details).

To solve the high-resolution structure of RcsD_Kp_, a total of 7330 curated exposures were used to pick 8 million particles using blob picker. These particles underwent rigorous 2D classifications and multi-class *ab initio* reconstructions to isolate a set of high-quality particles. This subset was then used for template picking, resulting in 8.2 million particles. Several rounds of 2D classifications and multi-class heterogeneous refinements were employed to curate further and refine the particles, ultimately yielding 274K high-quality particles. These particles were then subjected to a reference-based motion correction, non-uniform refinement, and local refinement to produce the 3.16 Å resolution map shown in this study. Both the sharpened and DeepEM maps were used for model building (see Fig. S10 for additional details).

To determine the high-resolution structure of RcsD_Ec_, a total of 8,276 curated exposures were used to pick 13.6 million particles using blob picker. These particles underwent multiple rounds of 2D classifications and multi-class *ab initio* reconstructions to isolate a set of high-quality particles and generate preliminary maps. This refined subset was then used for template-based picking, resulting in 7.3 million particles. The particles from both blob and template-based picking were combined, and duplicates removed. Following this, the dataset was further processed through several rounds of 2D classifications and multi-class heterogeneous refinements to ensure the highest quality of particle selection. Additionally, TOPAZ was employed for another round of picking, resulting in 3.7 million particles. Particles from all three picking methods were combined, and duplicates removed. Several rounds of multi-class heterogeneous refinements were employed to curate further and refine the particles, ultimately yielding 511K high-quality particles. These particles were then subjected to a reference-based motion correction, non-uniform refinement, and local refinement to produce the 2.6 Å resolution map shown in this study. Since CryoSPARC appeared to have overestimated the resolution (2.29 Å), the resolution from Phenix is reported, as it agrees with the manual inspection of the map. Both the sharpened and DeepEM maps were used for model building (see Fig. S11 for additional details).

To determine the high-resolution structure of RcsC_Kp_, a total of 11,552 curated exposures were used to pick 15 million particles using blob picker. These particles underwent multiple rounds of 2D classifications and multi-class *ab initio* reconstructions to isolate a set of high-quality particles and generate preliminary maps. This refined subset was then used for template-based picking, resulting in 17.7 million particles. The particles from both blob and template-based picking were combined, and duplicates removed, yielding 20.9 million particles. Several rounds of multi-class heterogeneous refinements were employed to curate further and refine the particles, ultimately yielding 189K high-quality particles. These particles were then subjected to reference-based motion correction, non-uniform refinement, and local refinement to produce the 3.74 Å resolution map shown in this study. Both the sharpened and DeepEM maps were used for model building (see Fig. S13 for additional details).

To determine the high-resolution structure of RcsC_Ec_, a total of 10,607 curated exposures were used to pick 19.2 million particles using blob picker. These particles underwent multiple rounds of 2D classifications and multi-class *ab initio* reconstructions to isolate a set of high-quality particles and generate preliminary maps. This refined subset was then used for template-based picking, resulting in 13.6 million particles. The particles from both blob and template-based picking were combined, and duplicates were removed, yielding 18.8 million particles. Several rounds of multi-class heterogeneous refinements were employed to curate further and refine the particles, ultimately yielding 288K high-quality particles. These particles were then subjected to reference-based motion correction, non-uniform refinement, and local refinement to produce the 2.72 Å resolution map shown in this study. Both the sharpened and DeepEM maps were used for model building (see Figure S14 for additional details).

#### Model building and refinement

AlphaFold3^55^ models of IgaA, RcsD, RcsC, and RcsF were initially generated and fitted to the corresponding Cryo-EM maps in ChimeraX^56^, serving as the starting point for model building. The Fab57 fragment was manually built *de novo*. Subsequently, Phenix^57^ and Coot^58^ were used iteratively to build and refine the final structures shown in this work (see Supplementary Table S1-2 for more details). Further analysis and validation of the structures was done using ChimeraX plugins, Q-scores^59^, and ISOLDE^60^. UCSF ChimeraX and UCSF Chimera^61^ were used for figure generation and the visualization of Cryo-EM maps and models (see supplementary Table S7 for all software used).

### AlphaFold3 modeling of IgaA/RcsD and RcsF/IgaA/RcsD complex structures

AlphaFold3 was used to model the complex containing IgaA, RcsF, and RcsD(1-461) dimer^38,55,56^. We specifically chose models that align well (lowest rmsd) to our individual Cryo-EM structures RcsD and IgaA/RcsF complex.

