## Supplementary Material for "Structural and functional insights into the Rcs phosphorelay"

This document includes:
Supplementary tables, figures and figure legends

**A**

| Protein | Size (a.a) | % Identity | % Similarity |
|---|---|---|---|
| $RcsB_{Kp}/RcsB_{Ec}$ | 216/216 | 97 | 98 |
| $RcsF_{Kp}/RcsF_{Ec}$ | 135/134 | 83 | 93.3 |
| $IgaA_{Kp}/IgaA_{Ec}$ | 709/711 | 72.2 | 84.2 |
| $RcsD_{Kp}/RcsD_{Ec}$ | 885/890 | 71.6 | 85 |
| $RcsC_{Kp}/RcsC_{Ec}$ | 946/949 | 79.7 | 88.1 |

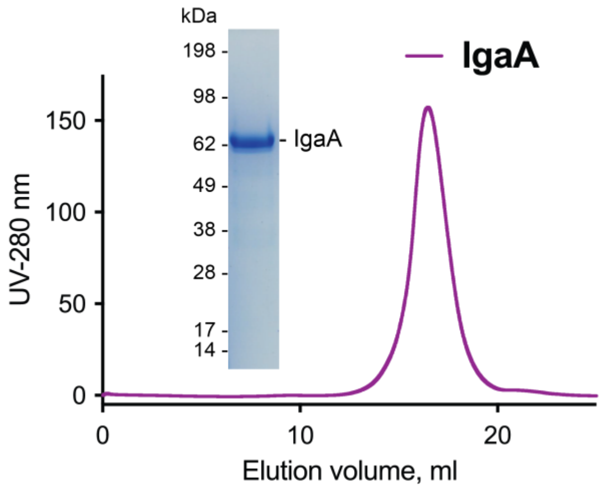

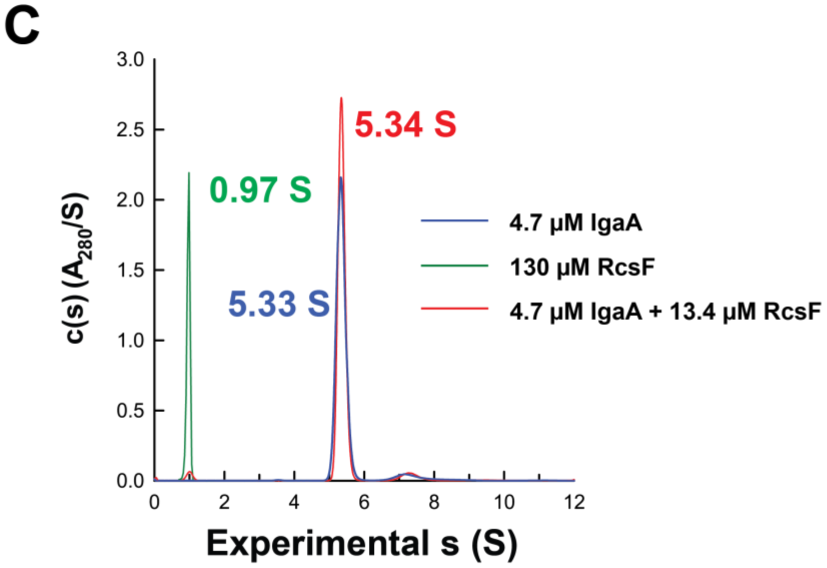

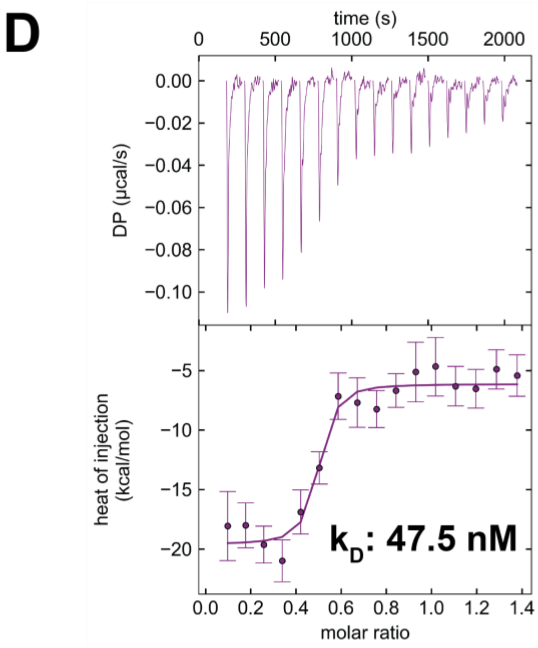

**Figure S1: A)** Sequence conservation of Rcs proteins. The proteins from *E. coli* and *K. pneumoniae* were analyzed using EMBOSS NEEDLE pairwise sequence alignment tool[1]. **B)** SEC and SDS-PAGE gel analysis of purified full-length IgaA. **C**) AUC results corresponding to panel A, including results for Apo RcsF and IgaA/RcsF complex. **D**) ITC results showing the binding interaction between full-length $IgaA_{Kp}$ and $RcsF_{Kp}$.

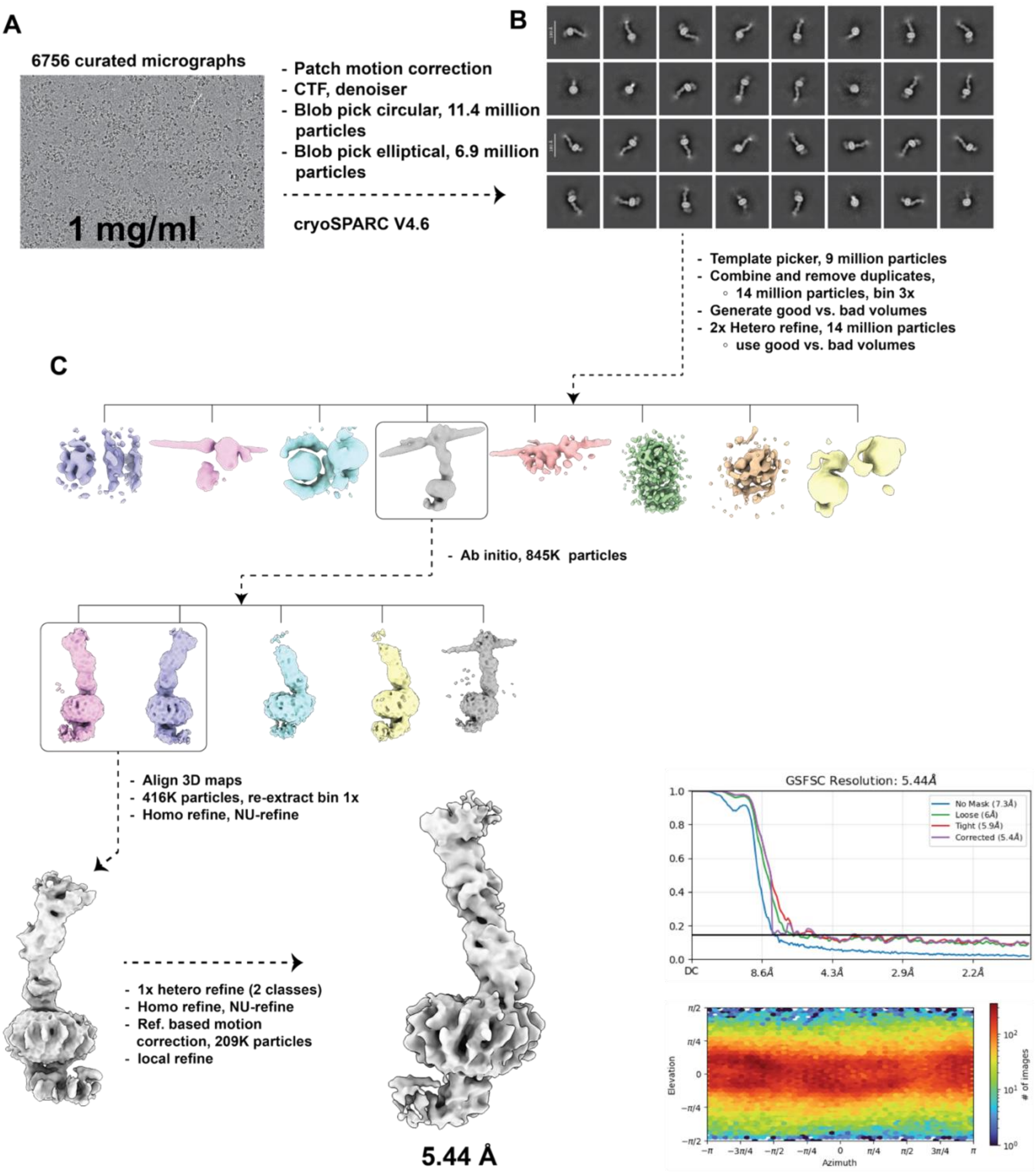

**Figure S2: Overview of the IgaA/RcsF data-processing workflow**. A) Representative motion-corrected micrograph. B) Selected 2D class averages. C) Schematic representation of the Cryo-EM data-processing pipeline for IgaA/RcsF complex. This includes the Fourier correlation plot (GFSC) showing the map quality and resolution and viewing direction plot (Azimuth) illustrating the orientation diversity of the data used to construct the structure.

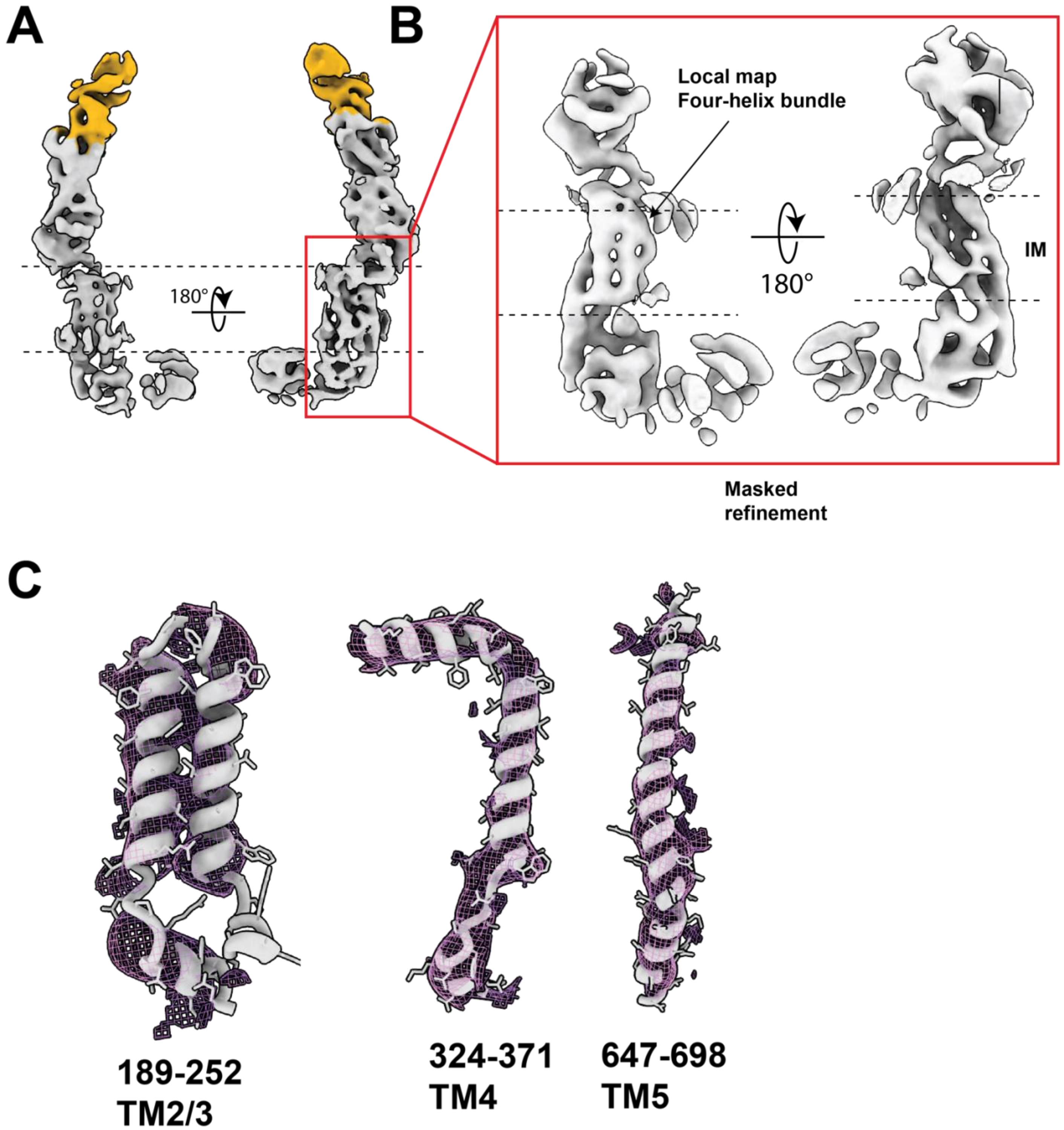

**Figure S3: Detailed views of IgaA/RcsF structure**. A) Same as in Fig. 2A but with the detergent micelle removed for improved clarity. B) Detailed views of the transmembrane domains obtained through masked refinement. C) Density maps and model fits for IgaA transmembrane domains; four of the five transmembrane domains have been successfully identified.

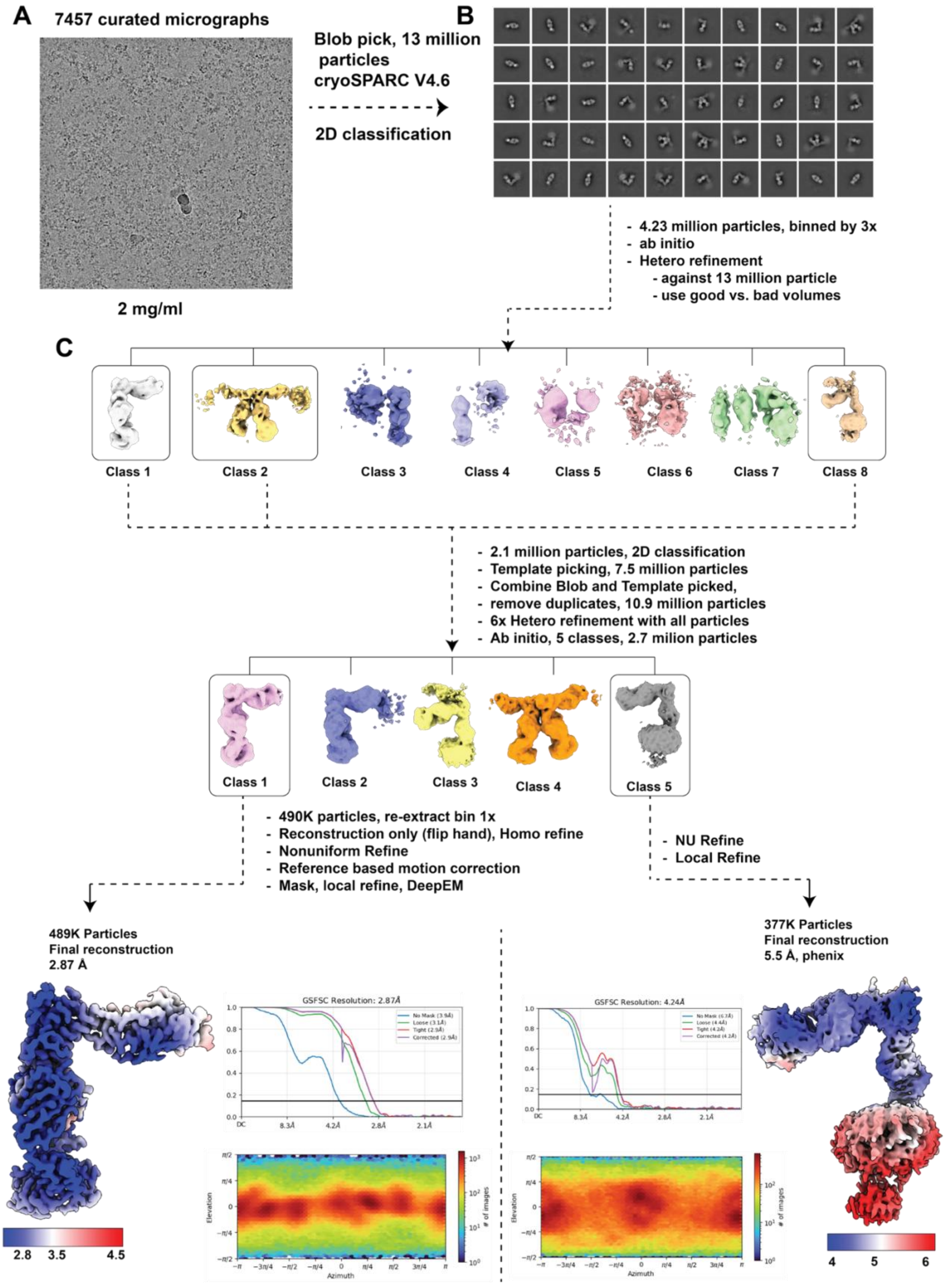

**Figure S4: overview of IgaA/Fab57 data processing workflow.** A) Representative motion-corrected micrograph. B) Selected 2D class averages used for further analysis. C) Schematic representation of the Cryo-EM data processing pipeline, including FSC curves and azimuth plots. It is important to note that the resolution estimation for the full-length map obtained from cryosparc appears to be overestimated, therefore, we report the unmasked resolution.

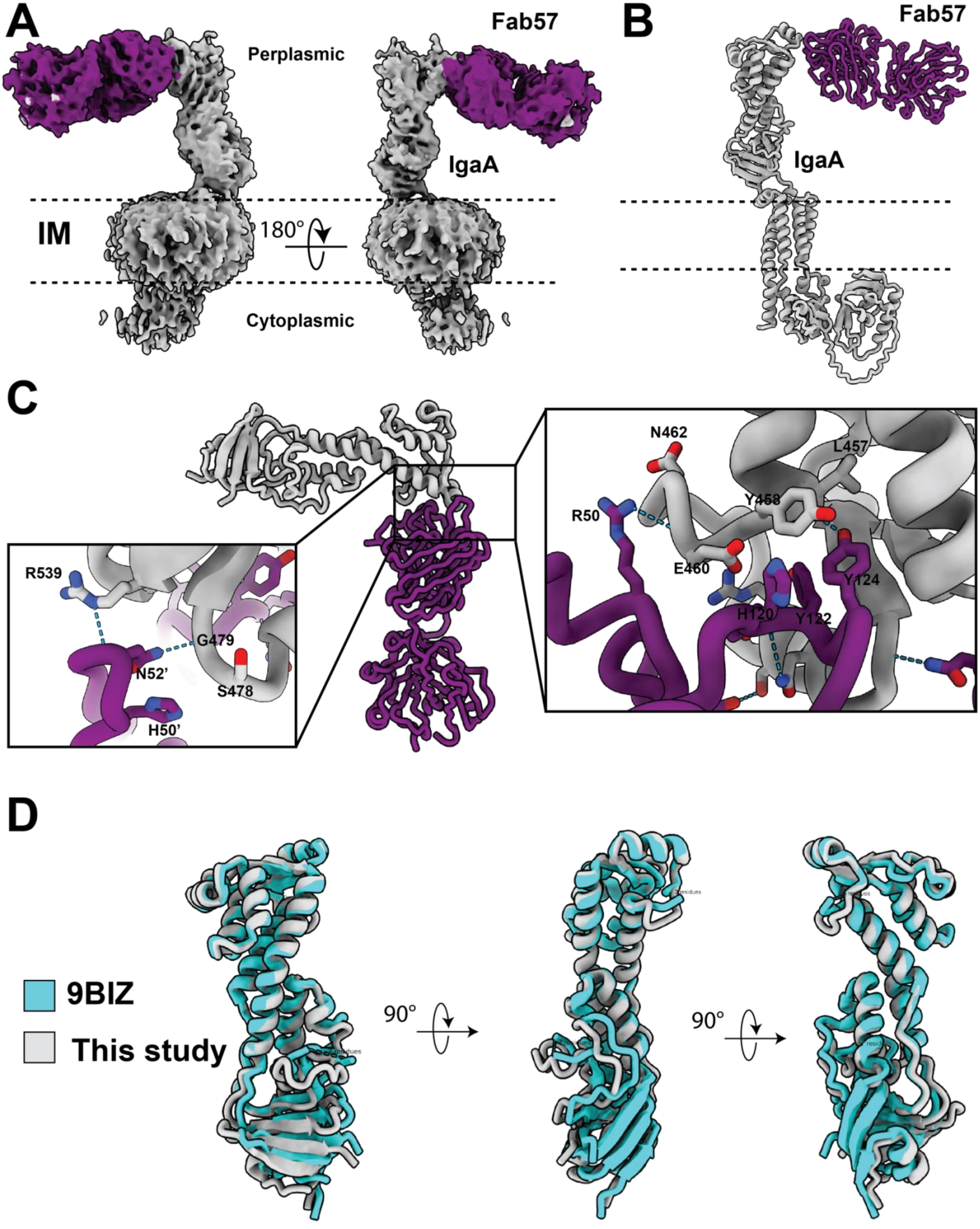
A
Fab57
Perplasmic
IgaA
IM
180°
Cytoplasmic
B
Fab57
IgaA
C
N462
L457
R50
Y458
E460
Y124
H120
Y122
R539
G479
N52'
S478
H50'
D
9BIZ
This study
90°
90°

**Figure S5: Detailed views of IgaA/Fab57 binding interface.** A) Multiple views of maps of the full-length IgaA in complex with Fab57. B) Alphafold3 model of apo IgaA and Fab57 fitted into the cryoEM map in panel A. C) Atomic-level views of the IgaA/Fab57 binding interface. D) Superposition of the crystal structure of IgaA (9BIZ) with IgaA structure from Fab57 complex; the overall rmsd is 0.89 Å.

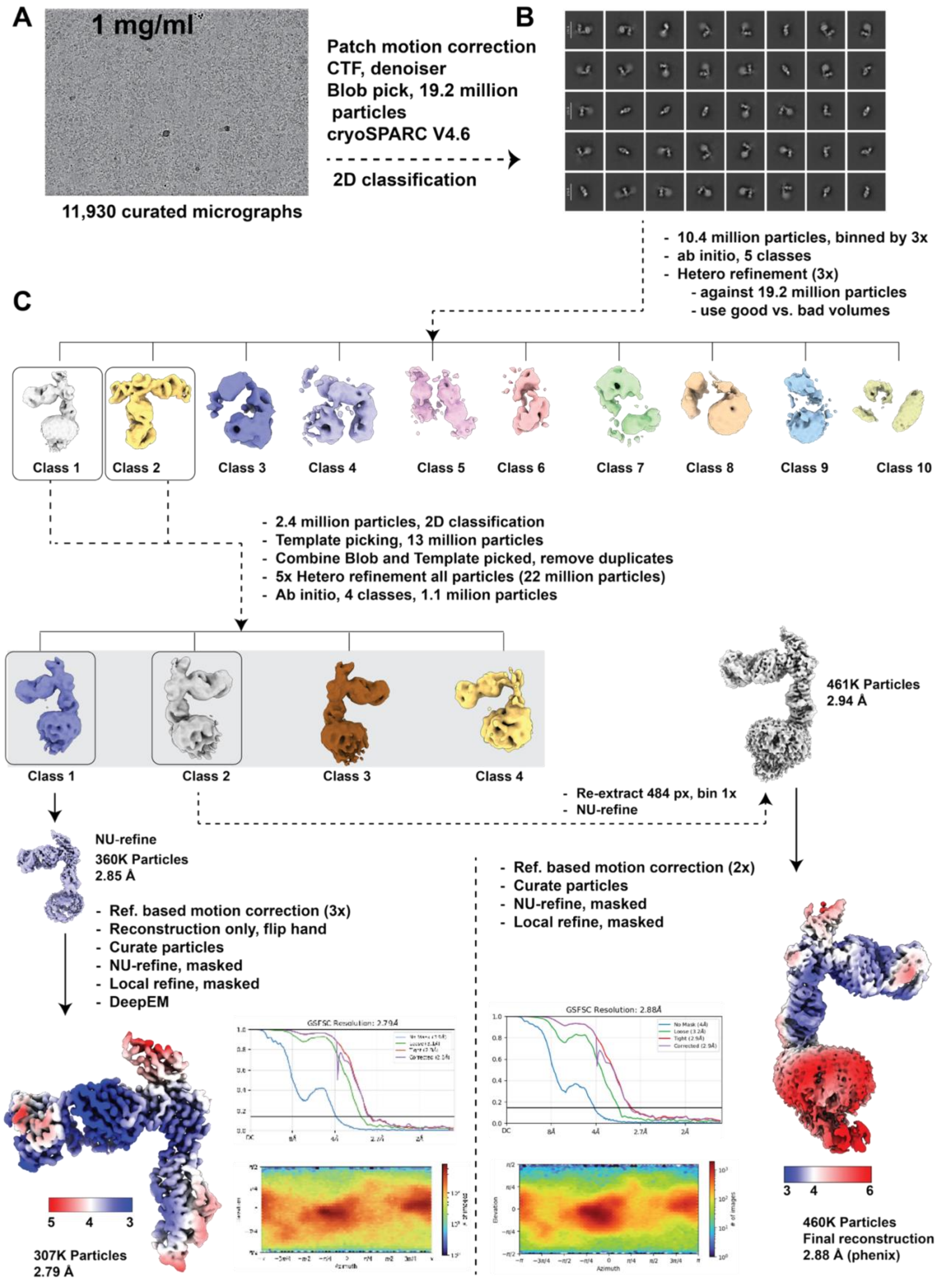

**Figure S6: Overview of the IgaA/Fab57/RcsF data-processing workflow.** A) Representative motion-corrected micrograph of IgaA/Fab57/RcsF particles. B) Selected 2D class averages that highlight distinct particle features used for further analysis. C) A schematic representation of the Cryo-EM data-processing pipeline, outlining key steps from particle picking to reconstruction, along with local-resolution maps, viewing direction plots, and gold-standard FSC plots, illustrating the resolution of the final maps generated from this data.

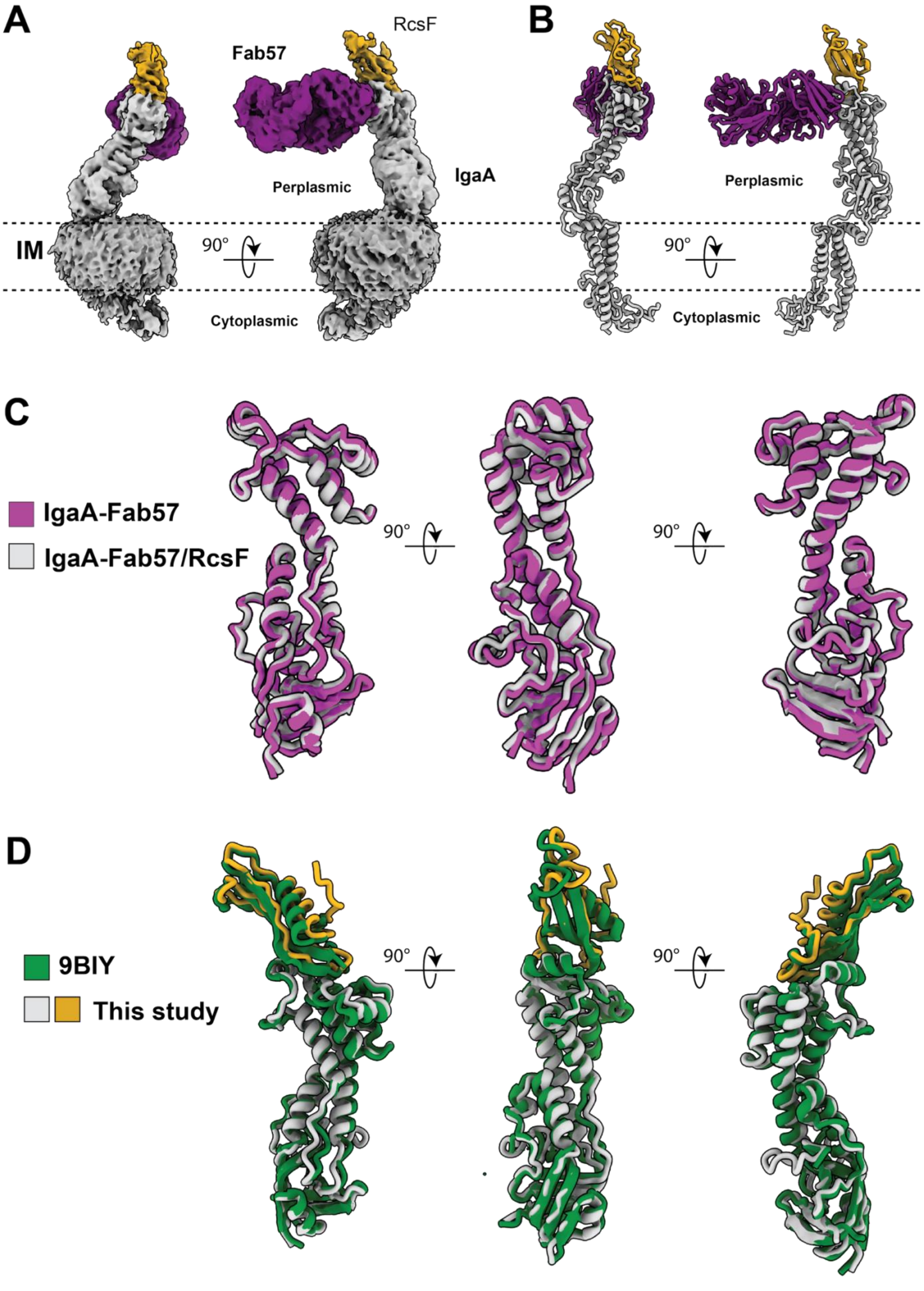
A
RcsF
Fab57
B
Perplasmic
IgaA
IM
90°
Cytoplasmic
Perplasmic
90°
Cytoplasmic
C
IgaA-Fab57
IgaA-Fab57/RcsF
90°
90°
D
9BIY
This study
90°
90°

**Figure S7: Detailed views of the IgaA/Fab57/RcsF complex consisting of the full-length IgaA and structure comparison.** A) Multiple views of maps of the full-length IgaA in complex with RcsF and Fab57. B) De novo built accurate structure of the IgaA/Fab57/RcsF complex into the map from panel A. C) Superposition of the Cryo-EM structure of IgaA/Fab57 with the IgaA/Fab57/RcsF complex; the overall rmsd is 0.9 Å. For clarity, Fab57 and RcsF have been removed. D) Superposition of the crystal structure of *E.coli* IgaA/RcsF (9BIY) with the IgaA/RcsF complex from this study; the overall rmsd is 1.3 Å.

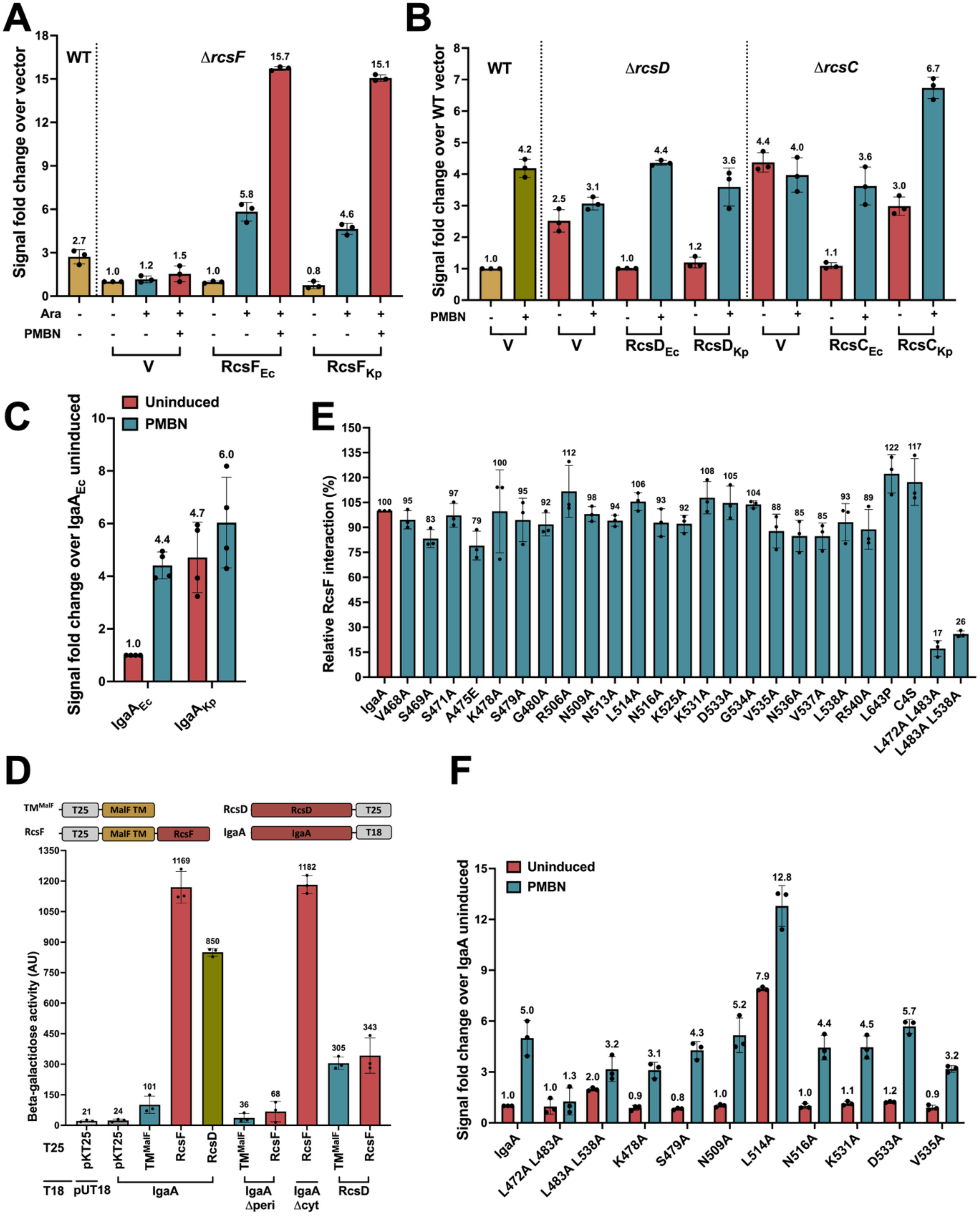
A
WT
ΔrcsF
Signal fold change over vector
Ara
PMBN
V
RcsF_Ec
RcsF_Kp
B
WT
ΔrcsD
ΔrcsC
Signal fold change over WT vector
PMBN
V
RcsD_Ec
RcsD_Kp
RcsC_Ec
RcsC_Kp
C
Uninduced
PMBN
Signal fold change over IgaA_Ec uninduced
IgaA_Ec
IgaA_Kp
D
TM^MalF
T25
MalF TM
RcsF
RcsD
IgaA
T18
Beta-galactidose activity (AU)
pKT25
pUT18
IgaA Δperi
IgaA Δcyt
E
Relative RcsF interaction (%)
IgaA
V468A
S469A
S471A
A475E
K478A
S479A
G480A
R506A
N509A
N513A
L514A
N516A
K525A
K531A
D533A
G534A
V535A
N536A
V537A
L538A
R540A
L643P
C4S
L472A L483A
L483A L538A
F
Uninduced
PMBN
Signal fold change over IgaA uninduced
IgaA
L472A L483A
L483A L538A
K478A
S479A
N509A
L514A
N516A
K531A
D533A
V535A

**Figure S8: Measuring Rcs phosphorelay signaling and binding. A) Signaling by Klebsiella RcsF protein in *E. coli.*** The strains WT (EAW8) and D*rcsF* (AP52) carry a *rprA* promoter fusion to mCherry ($P_{rprA}$::mCherry); mCherry fluorescence acts as a measure of Rcs activation. For the $P_{rprA}$::mCherry assay, the strains overexpressing pBAD33 (vector) or pBAD plasmids expressing $RcsF_{Ec}$ or $RcsF_{Kp}$ were grown in MOPS minimal glycerol medium containing chloramphenicol (25 mg/ml) and either 0.2% glucose or 0.01% arabinose at 37°C. The RFU at OD 0.4 compared to the uninduced (in the presence of glucose) D*rcsF* vector control (set to 1) is plotted. **B) Signaling by Klebsiella RcsD and RcsC proteins in *E. coli.*** The strain D*rcsD* (EAW19) overexpressing pBAD24 (vector), pBAD-$RcsD_{Ec}$ or $RcsD_{Kp}$ was grown in MOPS minimal glucose medium containing ampicillin (100 mg/ml) at 37°C. PMBN (20 mg/ml) was used for Rcs induction. The RFU at OD 0.4 compared to the uninduced WT vector control (set to 1) is plotted. The strain D*rcsC* (EAW91) overexpressing pBAD24 (vector), pBAD-$RcsC_{Ec}$ or $RcsC_{Kp}$ was assayed similarly. **C) Signaling by Klebsiella IgaA protein in *E. coli.*** The IgaA alleles are in the chromosome in a D*rcsD* strain carrying a $P_{rpra}$:mCherry reporter fusion. These strains with $IgaA_{Ec}$ (AP150) or $IgaA_{Kp}$ (AP149) were transformed with either pBAD24 vector (devoid of RcsD) or pEAW11 (*rcsD*$^{+}$) and their fluorescence assayed during growth in MOPS minimal glucose medium (with ampicillin). PMBN (20 µg/ml) was added in the medium at the beginning for Rcs activation. The RFU at OD 0.4 for each strain relative to the uninduced *igaA*+ *rcsD*$^{+}$ strain (set to 1) is plotted. **D.** BACTH for **interaction of *E. coli* Rcs proteins.** Beta-galactosidase activity was measured in a Δ*cyaA* strain (BTH101) expressing two plasmids encoding the T18 and T25 domains of adenylate cyclase fused to the proteins of interest, as shown at the top, and the expression measured compared to background. Note that the RcsF constructs all contain the MalF-TM domain in place of the usual RcsF lipoprotein domain. The IgaA/RcsD/RcsF protein fusion plasmids paired with their cognate vector had very low activity and were used as controls. **E. Interaction of IgaA mutants with RcsF**. The interaction of IgaA point mutants was tested with RcsF by BACTH in BTH101 as in S8E. **F. Signaling by IgaA mutant strains** Signaling was assayed with chromosomal *igaA* mutants, using the same method as Fig. S8C.

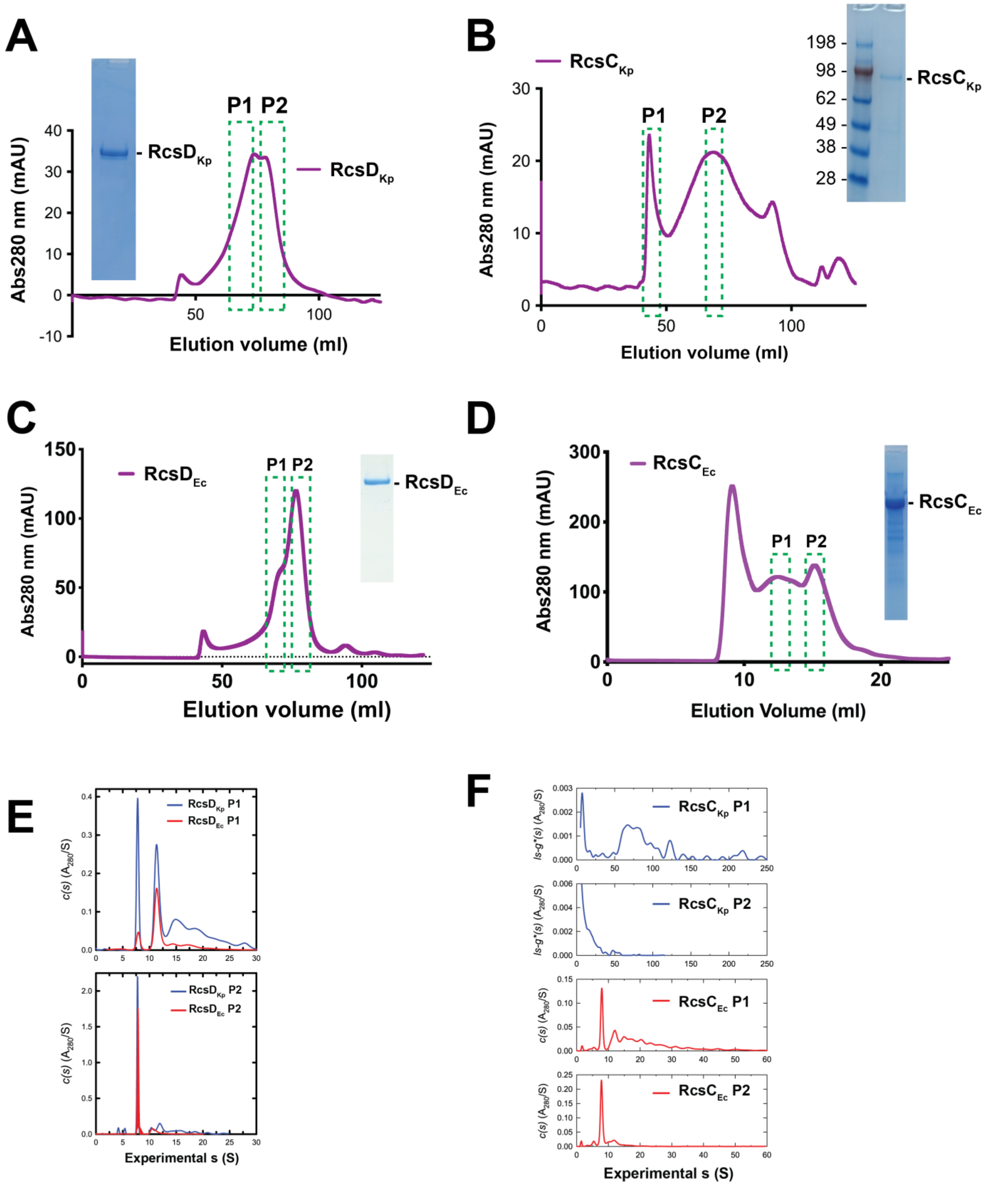

**Figure S9:** Biophysical studies of RcsD and RcsC. **A-D)** Size-exclusion chromatography (SEC) and SDS-PAGE gel analysis of purified RcsD and RcsC (from both *E. coli* and *K. pneumoniae*). Green boxes indicate the peaks/fractions used for AUC analysis. **E-F)** AUC results corresponding to panel A-D.

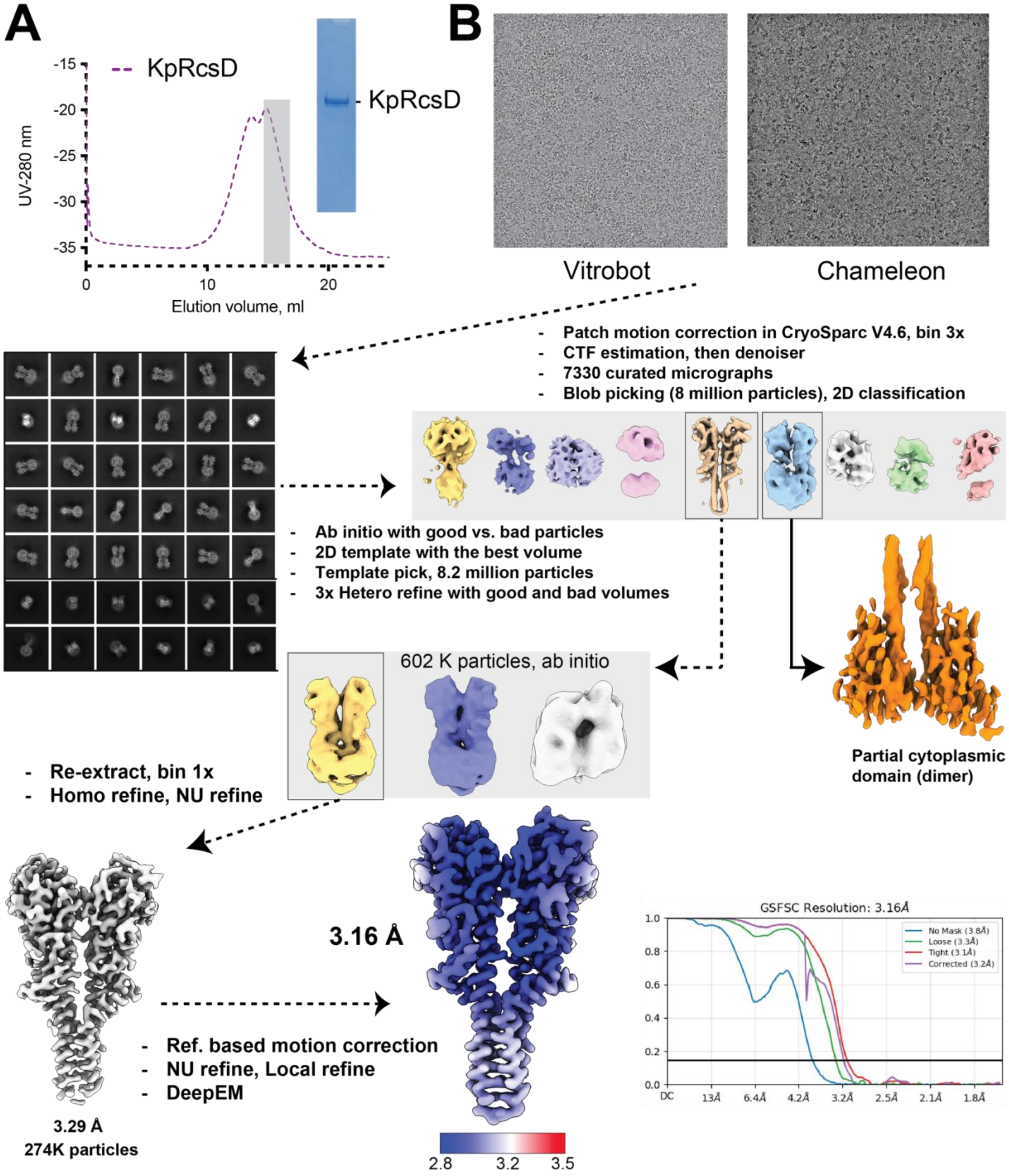

**Figure S10: RcsD$_{Kp}$ data processing workflow. A)** Chromatogram showing full-length RcsD purified by SEC column. The fractions used for structural studies are shaded gray. **B)** Schematic showing Cryo-EM data processing pipeline for RcsD. Note that only the TM/Periplasmic domains are visualized.

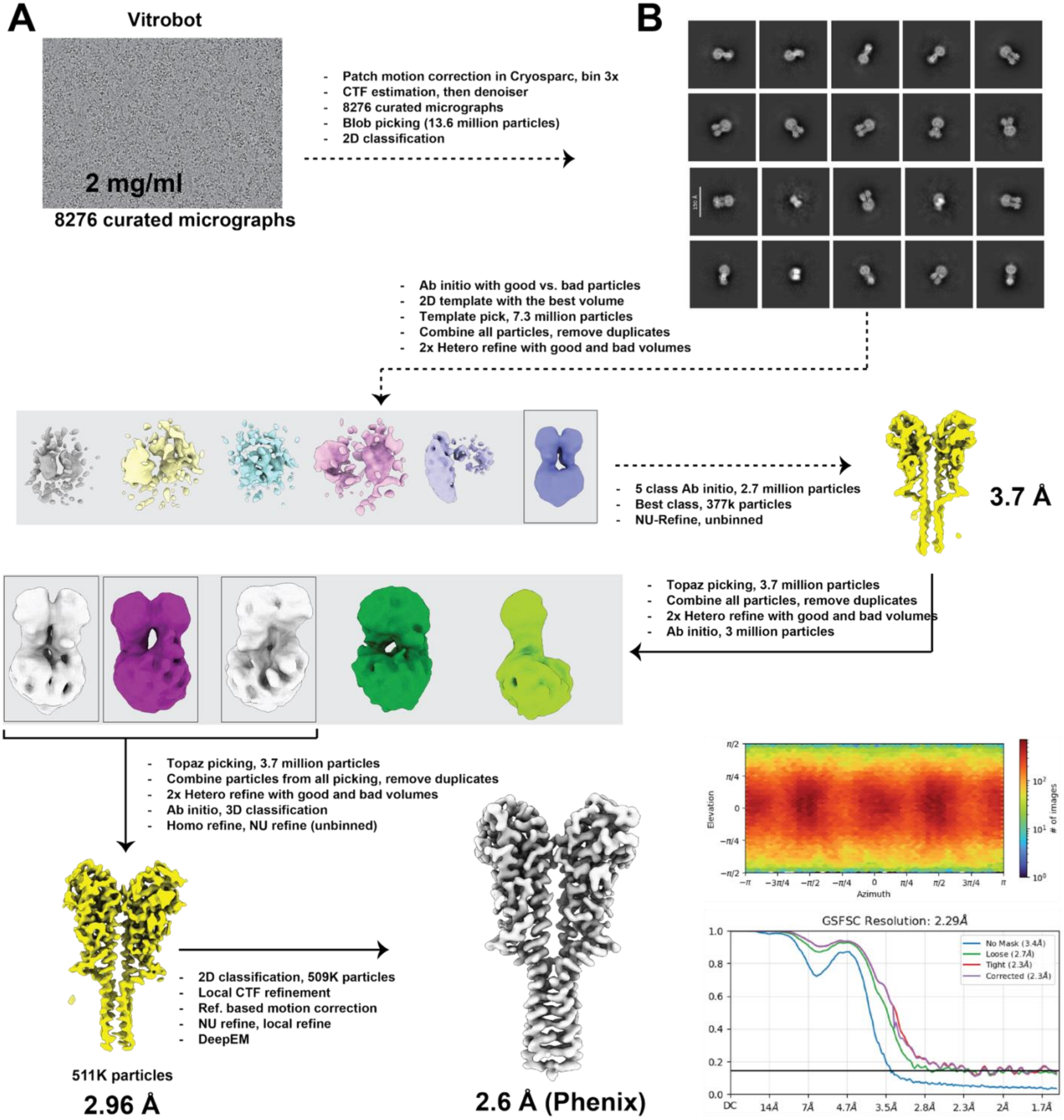

**Figure S11: EcRcsD data processing workflow. A)** Representative motion-corrected micrograph of EcRcsD particles. **B)** Examples of selected 2D class averages and schematic diagram of the Cryo-EM data-processing pipeline, viewing direction plot, and gold-standard FSC plot. Due to overfitting and resolution overestimation error in cryosparc, we are reporting the value calculated by Phenix, which is 2.6 Å.

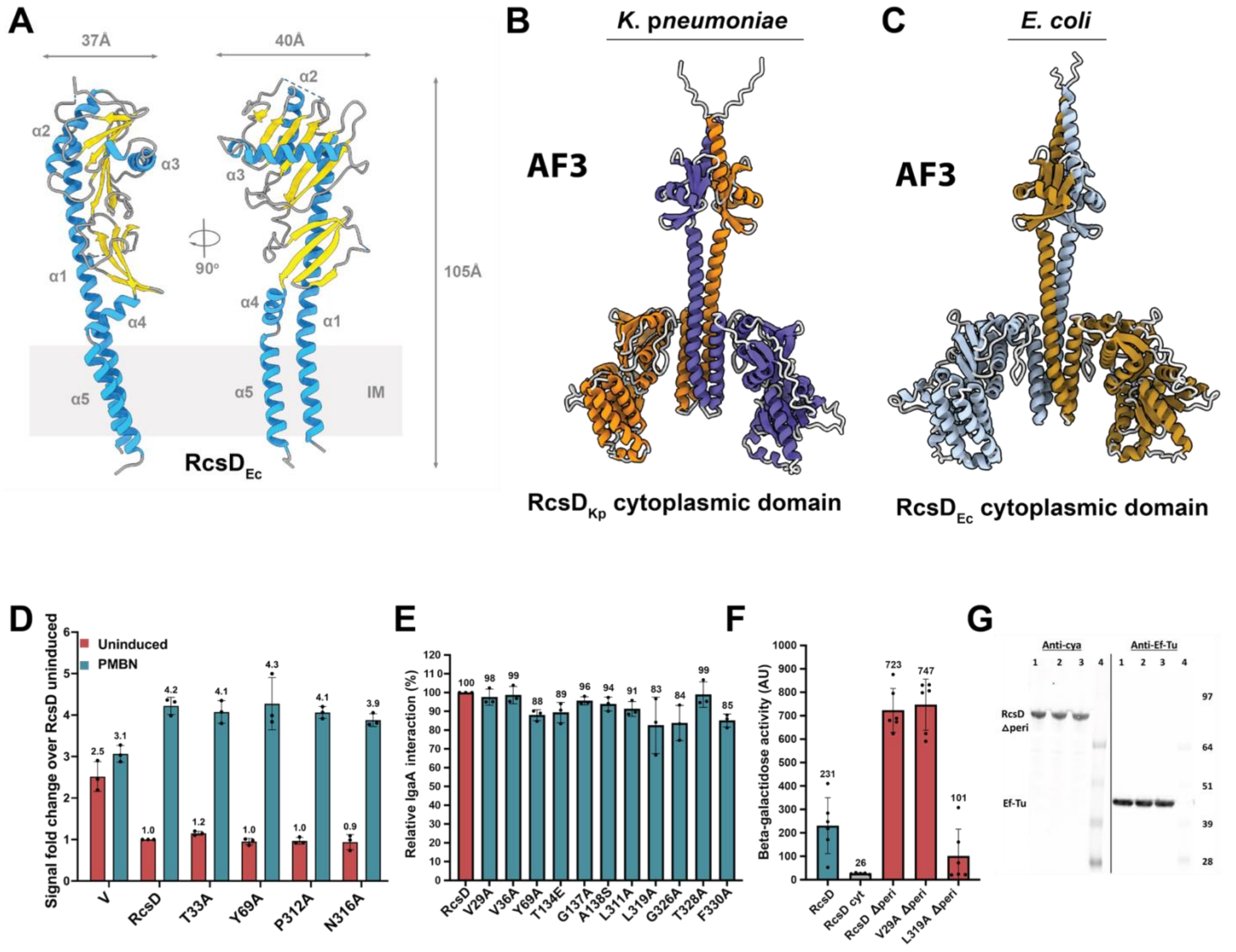

**Figure S12: A)** Domain organization of $RcsD_{Ec}$ (TM1/2 + Periplasmic, based on Cryo-EM structures)**. B-C)** AlphaFold3 predicted structures of cytoplasmic domains of RcsD homodimers from *K. pneumoniae* and *E. coli***. D)** Signaling by RcsD dimer interface mutants (assayed as in S8B). **E)** Interaction of RcsD mutants with IgaA. BACTH in BTH101 was used to measure the interaction of RcsD mutants with IgaA as in S8E. IgaA/RcsD interaction is set to 100 and other interactions are plotted relative to this. **F)** BACTH for RcsD homodimer formation. Homodimerization of various RcsD constructs was determined using BACTH in BTH101. **G)** Expression of RcsD mutants. Western blot of the T18 fusion constructs of RcsD Dperi, RcsD V29A Dperi, and RcsD L319A Dperi, expressed in a *cya*$^{+}$ strain (*DH5 alpha lacIQ*), is depicted here. Samples were probed on parallel gels with either the anti-CyaA antibody or the anti Ef-Tu antibody (used as a control).

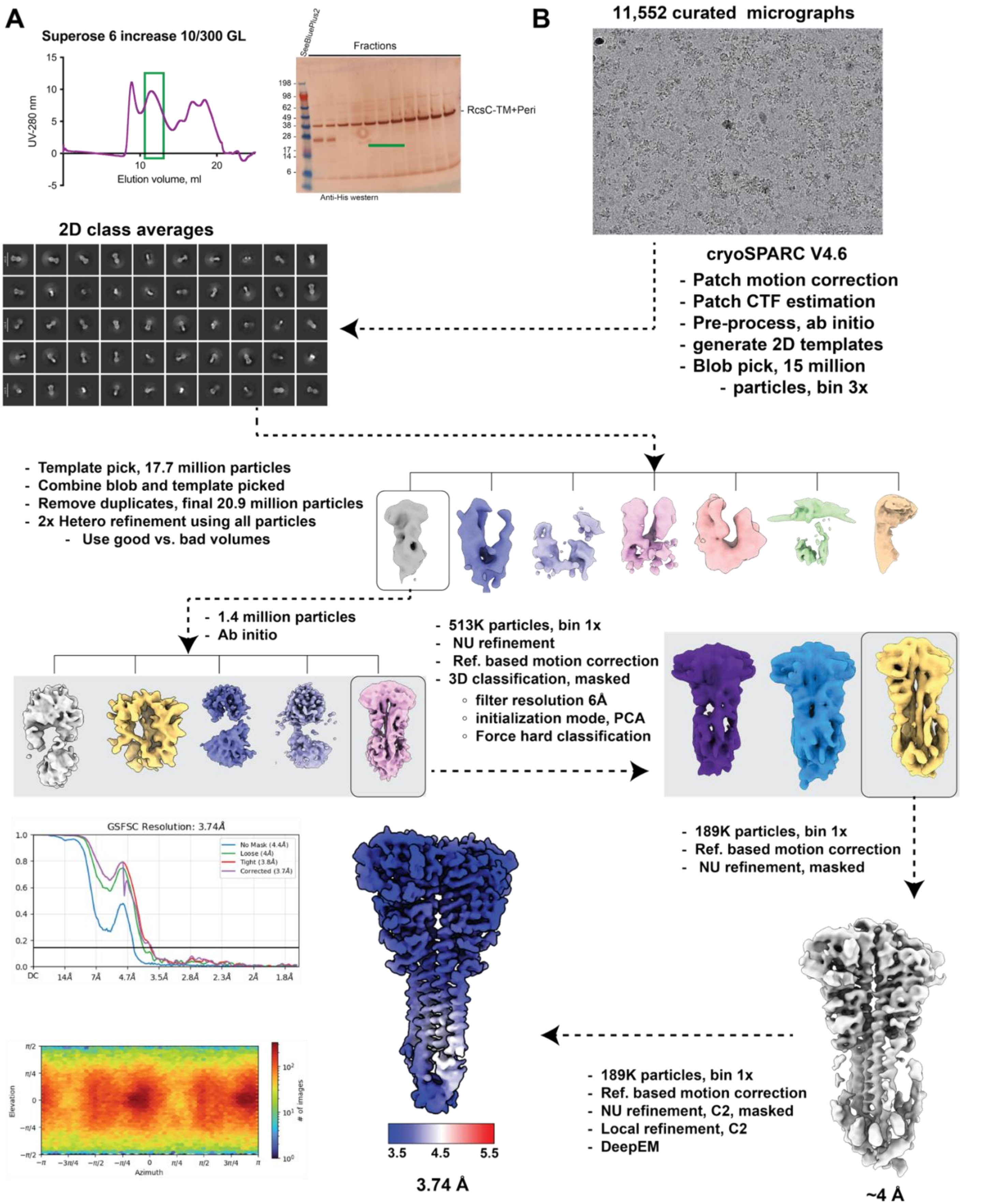

**Figure S13:** ***K. pneumoniae*** **RcsC data processing workflow.** A) Representative motion-corrected micrograph of RcsC$_{Kp}$ particles. B) Examples of selected 2D class averages and schematic diagram of the Cryo-EM data-processing pipeline, including local-resolution map, viewing direction plot, and gold-standard FSC plot.

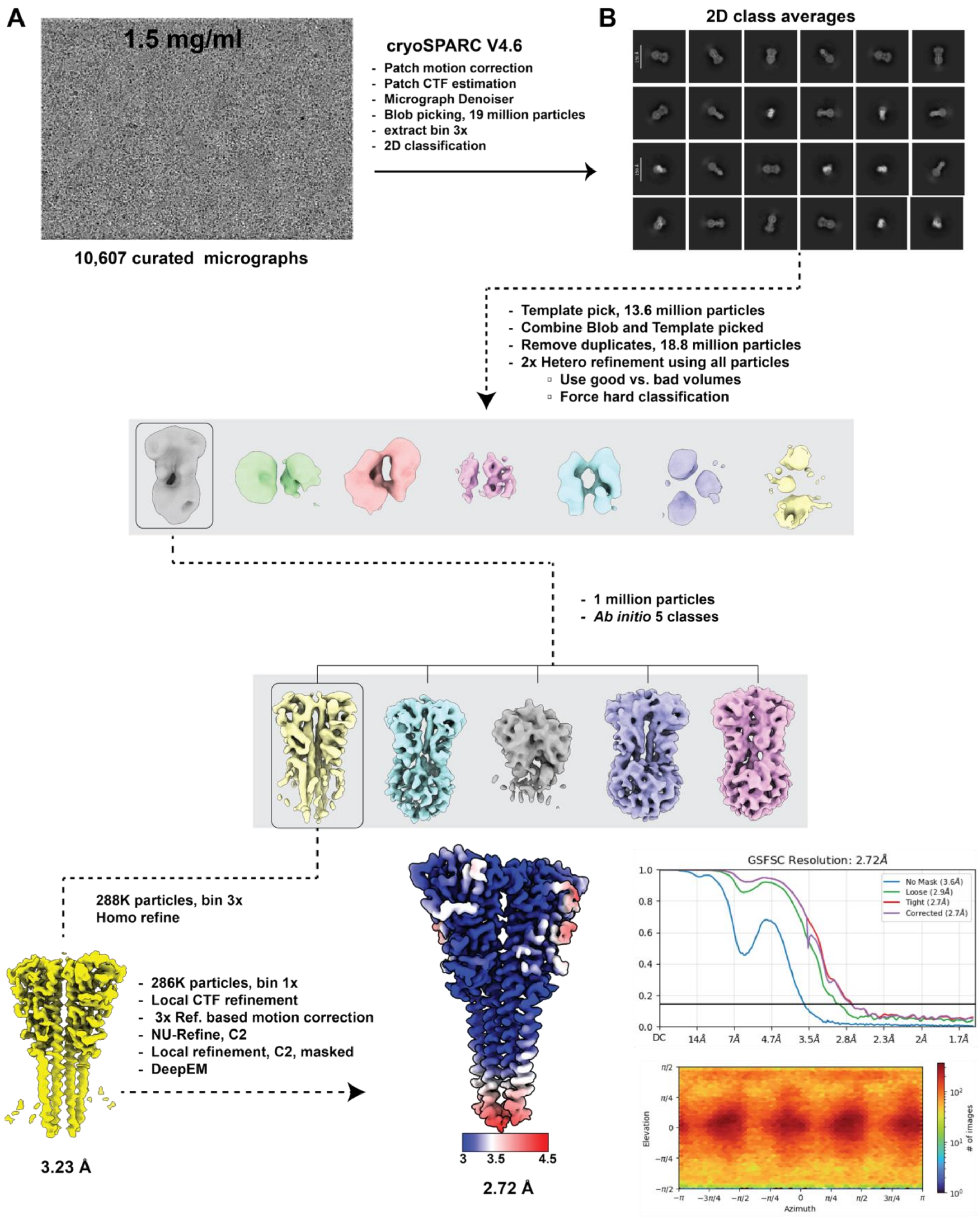

**Figure S14:** ***E. coli*** **RcsC data processing workflow.** A) Representative motion-corrected micrograph of $RcsC_{Ec}$ particles. B) Examples of selected 2D class averages and schematic diagram of the Cryo-EM data-processing pipeline, including local-resolution map, viewing direction plot, and gold-standard FSC plot.

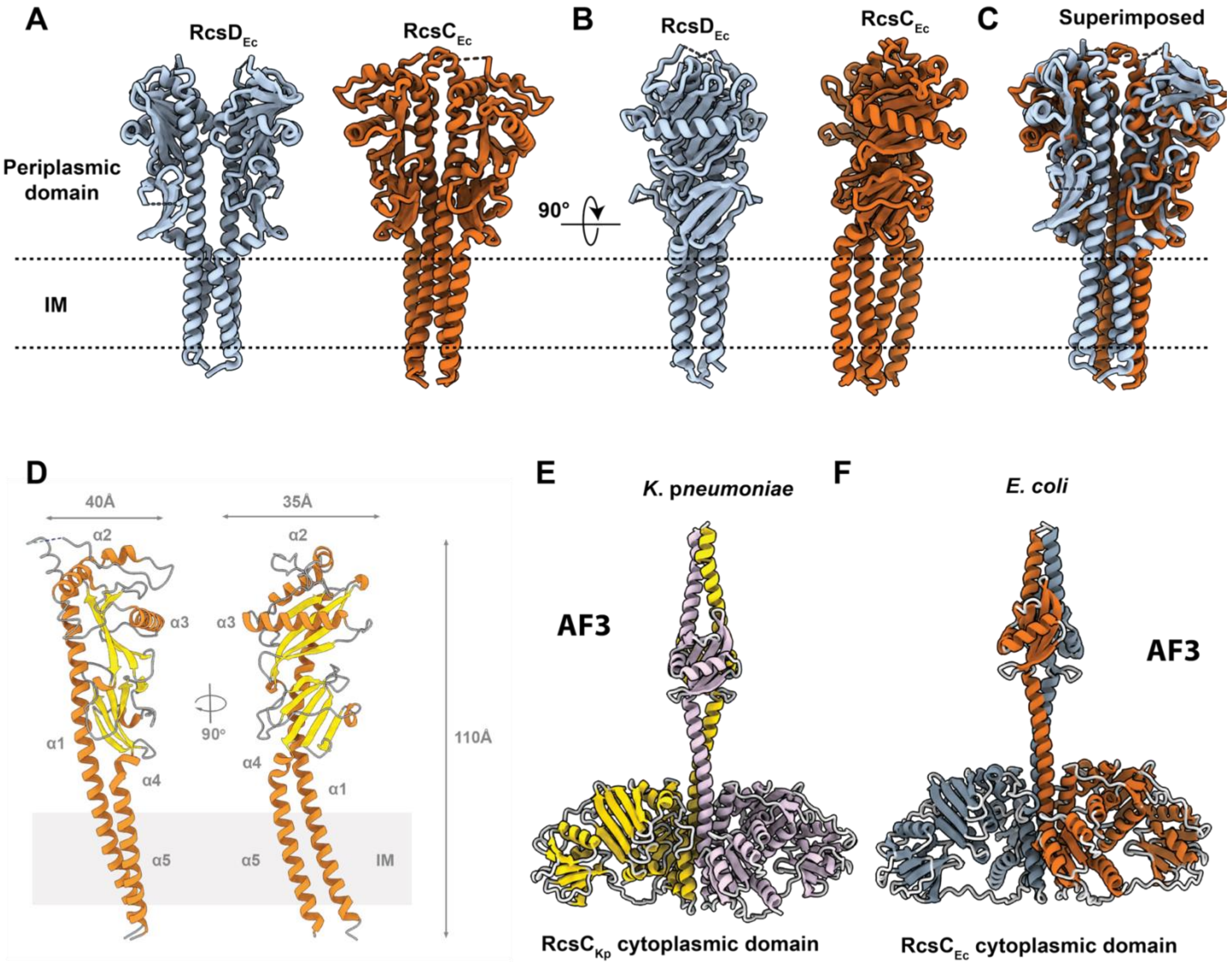

**Figure S15: Comparison of RcsD and RcsC Structures. A-B)** Side-by-side views of RcsD$_{Ec}$ and RcsC$_{Ec}$ Cryo-EM structures. **C)** Superposition of RcsD$_{Ec}$ and RcsC$_{Ec}$. **D)** Domain organization of RcsC$_{Ec}$ (TM1/2 + Periplasmic, based on Cryo-EM structures). **E-F)** AlphaFold3 predicted structures of cytoplasmic domains of RcsC homodimers from *K. pneumoniae* and *E. coli*.

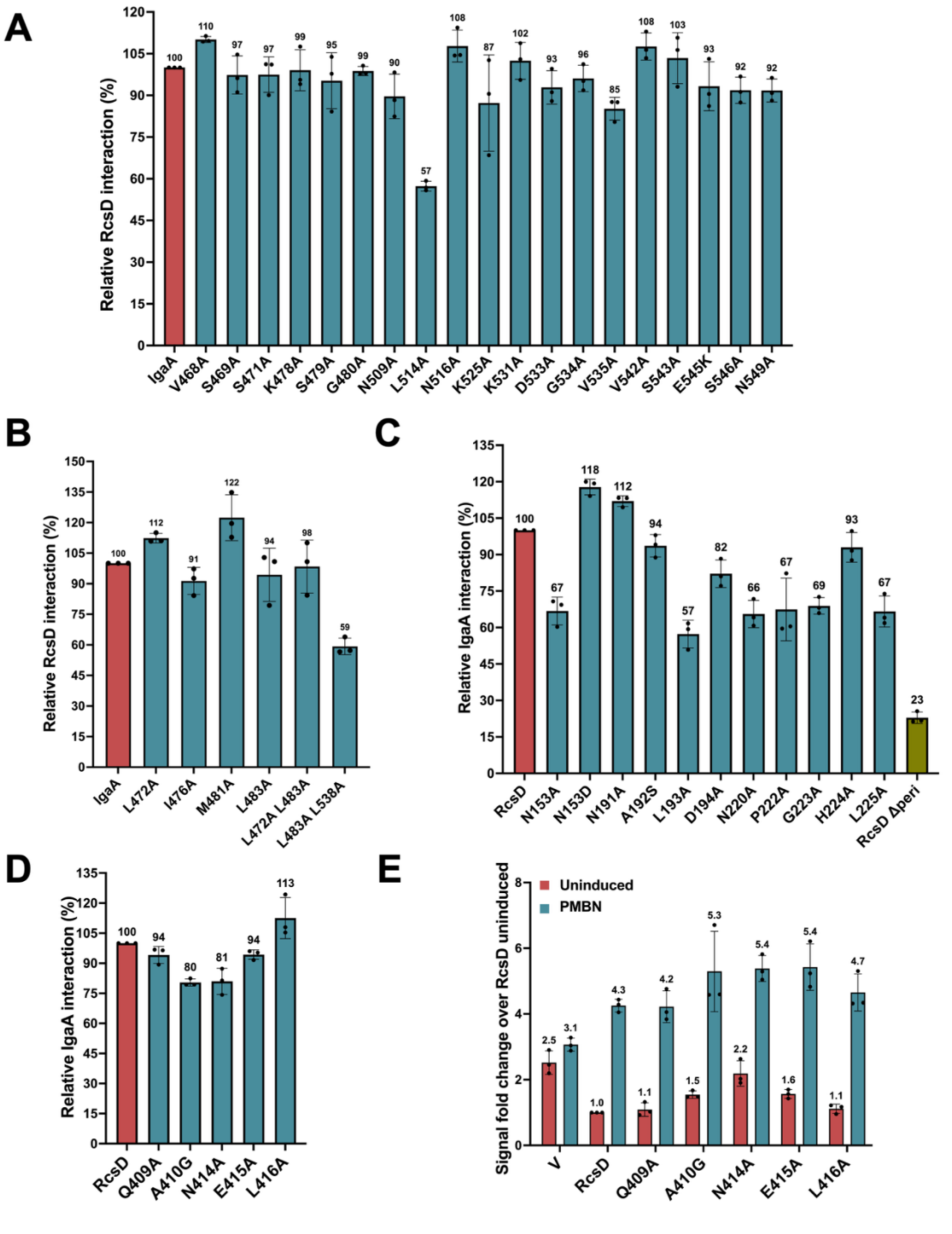

**Figure S16: A) Interaction of IgaA mutants with RcsD.** The interaction of RcsD was tested with IgaA and IgaA point mutants using BACTH as described in S8F. These IgaA mutants did not show a significant decrease in RcsD interaction. **B) Interaction of RcsF-defective IgaA mutants with RcsD.** The interaction of RcsD was tested with IgaA and IgaA point mutants in BTH101. These IgaA mutants have low interaction in RcsF but interact well with RcsD. **C) Interaction of RcsD periplasmic mutants with IgaA**. The binding of RcsD and its periplasmic point mutants to IgaA was tested using BACTH in BTH101. **D) Interaction of RcsD periplasmic mutants with IgaA**. The binding of RcsD and its periplasmic point mutants to IgaA was tested using BACTH in BTH101 as above. **E**) Signaling by RcsD cytoplasmic mutants. The D*rcsD* strain (EAW19) was transformed with either pBAD24 vector or RcsD mutants and their fluorescence assayed as described in S8B.

### Supplementary Tables

**Table S1: Cryo-EM data collection parameters, image processing and structure model building statistics**

| | $IgaA_{Kp}/RcsF_{Kp}$ | Full-length $IgaA_{Kp}$/Fab57 | Full-length $IgaA_{Kp}$-Fab57/$RcsF_{Kp}$ |
|---|---|---|---|
| **EMDB-ID** | 75234 | 75236 | 75238 |
| **Data collection** | | | |
| Microscope | Glacios | Titan Krios | Titan Krios |
| Voltage (kV) | 200 | 300 | 300 |
| Magnification | 45000x | 45000x | 105,000x |
| Pixel size (Å) | 0.89 | 0.86 | 0.824 |
| Electron dose ($e^-/Å^2$) | 65.45-70.43 | 60.24 | 46-58 |
| Number of frames | 32 | 28 | 40 |
| Defocus range | -0.8 - 2.4 | -0.8 - 2.4 | -0.8 - 2.8 |
| **Image processing** | | | |
| Micrographs selected (#) | 7,619 | 7,457 | 11,930 |
| Total extracted particles (#) | 11.4 | 13 mil | 22 mil |
| Final particles used (#) | 209K | 377K | 460K |
| Symmetry imposed | C1 | C1 | C1 |
| Map resolution, FSC 0.143 (Å) | 5.44 | 5.5 | 2.88 |
| Map sharpening *B*-factor ($Å^2$) | -341.4 | -156.8 | -85.1 |

**Table S2: Cryo-EM data collection parameters, image processing and structure model building statistics**

| | $IgaA_{Kp}$/Fab57 | $IgaA_{Kp}$-Fab57/$RcsF_{Kp}$ | $RcsD_{Kp}$ | $RcsD_{Ec}$ | $RcsC_{Kp}$ | $RcsC_{Ec}$ |
|---|---|---|---|---|---|---|
| **EMDB-ID** | 75235 | 75237 | 75239 | 75240 | 75241 | 75242 |
| **PDB-ID** | 10KG | 10KH | 10KI | 10KJ | 10KK | 10KL |
| **Data collection** | | | | | | |
| Microscope | Titan Krios | Titan Krios | Titan Krios | Titan Krios | Titan Krios | Titan Krios |
| Voltage (kV) | 300 | 300 | 300 | 300 | 300 | 300 |
| Magnification | 45,000x | 105,000x | 105,000x | 105,000x | 105,000x | 105,000x |
| Pixel size (Å) | 0.86 | 0.824 | 0.858 | 0.824 | 0.858 | 0.824 |
| Electron dose ($e^-/Å^2$) | 60.24 | 46-58 | 60 | 59.12 | 61 | 55.26 |
| Number of frames | 28 | 40 | 40 and 50 | 40 | 40 | 40 |
| Defocus range | -0.8 - 2.4 | -0.8 - 2.8 | -0.8 - 2.2 | -0.8 - 2.2 | -0.8 - 2.6 | -0.8 - 2.6 |
| **Image processing** | | | | | | |
| Micrographs selected (#) | 7,457 | 11,930 | 7,330 | 8,276 | 11,552 | 10,607 |
| Total extracted particles (#) | 13 mil | 22 mil | 8.2 mil | 13.6 mil | 15 mil | 19 mil |

| | | | | | | |
|---|---|---|---|---|---|---|
| Final particles used (#) | 489K | 307K | 274K | 511K | 189K | 286K |
| Symmetry imposed | C1 | C1 | C1 | C2 | C2 | C2 |
| Map resolution, FSC 0.143 (Å) | 2.87 | 2.79 | 3.16 | 2.6 | 3.74 | 2.72 |
| Map sharpening *B*-factor (Å$^2$) | -102.7 | -87.5 | -143.2 | -70.5 | -117.6 | -91.8 |
| **Model Refinement & validation** | | | | | | |
| r.m.s.d. Bond lengths (Å) | 0.006 | 0.004 | 0.004 | 0.009 | 0.007 | 0.005 |
| r.m.s.d. Bond angles (°) | 0.835 | 0.588 | 0.695 | 0.981 | 0.912 | 0.838 |
| Ramachandran Favored (%) | 95.2 | 97.74 | 97.81 | 95.27 | 93.29 | 97.97 |
| Ramachandran Allowed (%) | 4.8 | 2.26 | 2.91 | 4.73 | 6.71 | 2.03 |
| Ramachandran Disallowed (%) | 0.0 | 0.0 | 0.0 | 0.0 | 0.0 | 0.0 |
| Clash score | 2.42 | 8.02 | 7.17 | 5.35 | 6.33 | 3.93 |
| Rotamer outliers (%) | 0.0 | 0.0 | 0.0 | 0.0 | 0.0 | 0.0 |
| Map CC (volume) | 0.9 | 0.84 | 0.85 | 0.8 | 0.73 | 0.81 |
| **Model Composition** | | | | | | |
| Protein residues | 715 | 805 | 642 | 604 | 690 | 648 |

**Table S3: List of strains used in this study**

| Name | Genotype | Method of construction or reference |
|---|---|---|
| MG1655 | Wild-type *E. coli* K-12 | Lab collection |
| BTH101 | *F-, cya-99, araD139, galE15, galK16, rpsL1 (Str$^{r}$), hsdR2, mcrA1, mcrB1* | Karimova et al., 1998[2] |
| NEB DH5-alpha F'IQ | F′ *proA$^{+}$B$^{+}$ lacI$^{q}$* Δ*(lacZ)M15 zzf::Tn10* (Tet$^{R}$) / *fhuA2*Δ*(argF-lacZ)U169 phoA glnV44 Φ80Δ(lacZ)M15 gyrA96 recA1 relA1 endA1 thi-1 hsdR17* | New England Biolabs |
| EAW8 | Δ*araBAD*::$P_{rprA142}$-mCherry, Δ*araE*p $P_{CP6}$::*gent::*$P_{cp18}$-*araE* | Wall et al., 2020[3] |
| EAW12 | *rcsD*541(::FRT), *cya* | Wall et al., 2020[3] |
| EAW19 | Δ*araBAD*::$P_{rprA142}$-mCherry, *rcsD*541(::FRT), Δ*araEp* $P_{cp6}$*gent::*$P_{cp18}$-*araE* | Wall et al., 2020[3] |
| EAW90 | Δ*araBAD*::$P_{rprA142}$-mCherry, Δ*araE*p $P_{cp6}$*gent*::$P_{cp18}$-*araE*, *rcsD541(::FRT)*, Δ*igaA*::*kan-araC-kid* | Wall et al., 2020[3] |
| EAW91 | Δ*araBAD*::$P_{rprA142}$-mCherry, Δ*araEp* $P_{cp6}$*gent::*$P_{cp18}$-*araE,* Δ*rcsC91* | Wall et al., 2020[3] |
| AP24 | Δ*araBAD*::$P_{rprA142}$-mCherry, Δ*araEp* $P_{cp6}$*gent::*$P_{cp18}$-*araE, rcsD*541(::FRT), Δ*rcsF* | EAW19 + P1 (Keio JW0192) |
| AP27 | Δ*araBAD*::$P_{rprA142}$-mCherry, Δ*araE*p $P_{cp6}$*gent*::$P_{cp18}$-*araE*, *rcsD541(::FRT)*, Δ*rcsF, igaA N513A* | AP184 + P1 (Keio JW0192) |
| AP29 | Δ*araBAD*::$P_{rprA142}$-mCherry, Δ*araE*p $P_{cp6}$*gent*::$P_{cp18}$-*araE*, *rcsD541(::FRT)*, Δ*rcsF, igaA R540A* | AP216 + P1 (Keio JW0192) |
| AP51 | Δ*araBAD*::$P_{rprA142}$-mCherry, Δ*araE*p $P_{CP6}$::*gent::*$P_{cp18}$-*araE, rcsF::kan* | Petchiappan et al., 2024[4] |
| AP52 | Δ*araBAD*::$P_{rprA142}$-mCherry, Δ*araE*p $P_{CP6}$::*gent::*$P_{cp18}$-*araE,* Δ*rcsF* | AP51 + pCP20 |
| AP140 | Δ*araBAD*::$P_{rprA142}$-mCherry, Δ*araE*p $P_{cp6}$*gent*::$P_{cp18}$-*araE*, *rcsD541(::FRT)*, *igaA L438A L538A* | AP179 recombination with PCR product of oligos EAW213 and EAW214 on pAP105382 template |
| AP141 | Δ*araBAD*::$P_{rprA142}$-mCherry, Δ*araE*p $P_{cp6}$*gent*::$P_{cp18}$-*araE*, *rcsD541(::FRT)*, *igaA A94D* | AP179 recombination with PCR product of oligos EAW213 and EAW214 on pAP1094 template |

| | | |
|---|---|---|
| AP142 | Δ*araBAD*::P$_{rprA142}$-mCherry, Δ*araE*p P$_{cp6}$*gent*::P$_{cp18}$-*araE*, *rcsD541(::FRT)*, *igaA I476A* | AP179 recombination with PCR product of oligos EAW213 and EAW214 on pAP10476 template |
| AP144 | Δ*araBAD*::P$_{rprA142}$-mCherry, Δ*araE*p P$_{cp6}$*gent*::P$_{cp18}$-*araE*, *rcsD541(::FRT)*, *igaA L483A* | AP179 recombination with PCR product of oligos EAW213 and EAW214 on pAP10483 template |
| AP145 | Δ*araBAD*::P$_{rprA142}$-mCherry, Δ*araE*p P$_{cp6}$*gent*::P$_{cp18}$-*araE*, *rcsD541(::FRT)*, *igaA V535A* | AP179 recombination with PCR product of oligos EAW213 and EAW214 on pAP10535 template |
| AP146 | Δ*araBAD*::P$_{rprA142}$-mCherry, Δ*araE*p P$_{cp6}$*gent*::P$_{cp18}$-*araE*, *rcsD541(::FRT)*, *igaA N536A* | AP179 recombination with PCR product of oligos EAW213 and EAW214 on pAP10536 template |
| AP147 | Δ*araBAD*::P$_{rprA142}$-mCherry, Δ*araE*p P$_{cp6}$*gent*::P$_{cp18}$-*araE*, *rcsD541(::FRT)*, *igaA V537A* | AP179 recombination with PCR product of oligos EAW213 and EAW214 on pAP10537 template |
| AP148 | Δ*araBAD*::P$_{rprA142}$-mCherry, Δ*araE*p P$_{cp6}$*gent*::P$_{cp18}$-*araE*, *rcsD541(::FRT)*, *igaA L538A* | AP179 recombination with PCR product of oligos EAW213 and EAW214 on pAP10538 template |
| AP149 | Δ*araBAD*::P$_{rprA142}$-mCherry, Δ*araE*p P$_{cp6}$*gent*::P$_{cp18}$-*araE*, *rcsD541(::FRT)*, *igaA*$_{Kp}$ | AP179 recombination with PCR product of oligos EAW213 and EAW214 on pAP104 template |
| AP150 | Δ*araBAD*::P$_{rprA142}$-mCherry, Δ*araE*p P$_{cp6}$*gent*::P$_{cp18}$-*araE*, *rcsD541(::FRT)*, *igaA* | AP179 recombination with PCR product of oligos EAW213 and EAW214 on pEAW1 template |
| AP151 | Δ*araBAD*::P$_{rprA142}$-mCherry, Δ*araE*p P$_{cp6}$*gent*::P$_{cp18}$-*araE*, *rcsD541(::FRT)*, *igaA R97A* | AP179 recombination with PCR product of oligos EAW213 and EAW214 on pAP1097 template |
| AP152 | Δ*araBAD*::P$_{rprA142}$-mCherry, Δ*araE*p P$_{cp6}$*gent*::P$_{cp18}$-*araE*, *rcsD541(::FRT)*, *igaA R97S* | AP179 recombination with PCR product of oligos EAW213 and EAW214 on pAP10972 template |
| AP153 | Δ*araBAD*::P$_{rprA142}$-mCherry, Δ*araE*p P$_{cp6}$*gent*::P$_{cp18}$-*araE*, *rcsD541(::FRT)*, *igaA R506A* | AP179 recombination with PCR product of oligos EAW213 and EAW214 on pAP10506 template |
| AP179 | Δ*araBAD*::P$_{rprA142}$-mCherry, Δ*araE*p P$_{cp6}$*gent*::P$_{cp18}$-*araE*, *rcsD541(::FRT)*, Δ*igaA*::*kan-araC-kid,* pSIM6 | EAW90 + pSIM6 |
| AP183 | Δ*araBAD*::P$_{rprA142}$-mCherry, Δ*araE*p P$_{cp6}$*gent*::P$_{cp18}$-*araE*, *rcsD541(::FRT)*, *igaA M481A* | AP179 recombination with PCR product of oligos EAW213 and EAW214 on pAP10481 template |
| AP184 | Δ*araBAD*::P$_{rprA142}$-mCherry, Δ*araE*p P$_{cp6}$*gent*::P$_{cp18}$-*araE*, *rcsD541(::FRT)*, *igaA N513A* | AP179 recombination with PCR product of oligos EAW213 and EAW214 on pAP10513 template |

| | | |
|---|---|---|
| AP185 | Δ*araBAD*::$P_{rprA142}$-mCherry, Δ*araE*p $P_{cp6}$*gent*::$P_{cp18}$-*araE*, *rcsD541(::FRT)*, *igaA L472A* | AP179 recombination with PCR product of oligos EAW213 and EAW214 on pAP10472 template |
| AP186 | Δ*araBAD*::$P_{rprA142}$-mCherry, Δ*araE*p $P_{cp6}$*gent*::$P_{cp18}$-*araE*, *rcsD541(::FRT)*, *igaA K478A* | AP179 recombination with PCR product of oligos EAW213 and EAW214 on pAP10478 template |
| AP188 | Δ*araBAD*::$P_{rprA142}$-mCherry, Δ*araE*p $P_{cp6}$*gent*::$P_{cp18}$-*araE*, *rcsD541(::FRT)*, *igaA S479A* | AP179 recombination with PCR product of oligos EAW213 and EAW214 on pAP10479 template |
| AP215 | Δ*araBAD*::$P_{rprA142}$-mCherry, Δ*araE*p $P_{cp6}$*gent*::$P_{cp18}$-*araE*, *rcsD541(::FRT)*, *igaA L472A L483A* | AP179 recombination with PCR product of oligos EAW213 and EAW214 on pAP104722 template |
| AP216 | Δ*araBAD*::$P_{rprA142}$-mCherry, Δ*araE*p $P_{cp6}$*gent*::$P_{cp18}$-*araE*, *rcsD541(::FRT)*, *igaA R540A* | AP179 recombination with PCR product of oligos EAW213 and EAW214 on pAP10540 template |
| AP233 | Δ*araBAD*::$P_{rprA142}$-mCherry, Δ*araE*p $P_{cp6}$*gent*::$P_{cp18}$-*araE*, *rcsD541(::FRT)*, *igaA N509A* | AP179 recombination with PCR product of oligos EAW213 and EAW214 on pAP10509 template |
| AP234 | Δ*araBAD*::$P_{rprA142}$-mCherry, Δ*araE*p $P_{cp6}$*gent*::$P_{cp18}$-*araE*, *rcsD541(::FRT)*, *igaA L514A* | AP179 recombination with PCR product of oligos EAW213 and EAW214 on pAP10514 template |
| AP235 | Δ*araBAD*::$P_{rprA142}$-mCherry, Δ*araE*p $P_{cp6}$*gent*::$P_{cp18}$-*araE*, *rcsD541(::FRT)*, *igaA N516A* | AP179 recombination with PCR product of oligos EAW213 and EAW214 on pAP10516 template |
| AP236 | Δ*araBAD*::$P_{rprA142}$-mCherry, Δ*araE*p $P_{cp6}$*gent*::$P_{cp18}$-*araE*, *rcsD541(::FRT)*, *igaA K531A* | AP179 recombination with PCR product of oligos EAW213 and EAW214 on pAP10531 template |
| AP237 | Δ*araBAD*::$P_{rprA142}$-mCherry, Δ*araE*p $P_{cp6}$*gent*::$P_{cp18}$-*araE*, *rcsD541(::FRT)*, *igaA D533A* | AP179 recombination with PCR product of oligos EAW213 and EAW214 on pAP10533 template |

**Table S4: List of plasmids used in this study**

| Name | Description | Method of construction/ Reference |
|---|---|---|
| pBAD24 | pBR322-based vector for protein expression driven by the *araBAD* operon promoter (arabinose inducible) ($Amp^r$) | Guzman et al., 1995[5] |
| pBAD33 | pACYC184-based vector for protein expression driven by the *araBAD* operon promoter (arabinose inducible) ($Chl^r$) | Guzman et al., 1995[5] |
| pCP20 | Plasmid with temperature-sensitive origin of replication, encoding the FLP recombinase | Cherepanov et al., 1995[6] |
| pUT18 | Vector encoding the Cya T18 fragment under lac promoter control ($Amp^r$) | Karimova et al., 2001[7] |
| pKT25 | Vector encoding the Cya T25 fragment under lac promoter control ($Kan^r$) | Karimova et al., 2001[7] |
| pKNT25 | Vector encoding the Cya T25 fragment under lac promoter control ($Kan^r$) | Karimova et al., 2005[8] |
| pSIM6 | Plasmid with temperature-sensitive origin of replication, encoding l-Red cI857, gam-beta-exo ($Amp^r$) | https://ncifrederick.cancer.gov/ recombineering/strains-plasmids-and-primers |
| pSIM27 | Plasmid with temperature-sensitive origin of replication, encoding l-Red cI857, gam-beta-exo ($Tet^r$) | https://ncifrederick.cancer.gov/ recombineering/strains-plasmids-and-primers |
| pBAD28 | pBAD18, pACYC184 vector for protein expression driven by the *araBAD* operon promoter (arabinose inducible), ($Amp^r$) | Guzman et al., 1995[5] |
| pET28 | T7 promoter based for high-level protein expression, Kanamycin ($Kan^r$) | |
| **RcsF plasmids:** | | |
| pAP3330 | $RcsF_{Kp}$ cloned in pBAD33 vector | pBAD33 linearized with primers AP559 and AP560; Insert is gBlock AP_GF33kp |
| pAP3340 | RcsF cloned in pBAD33 vector | Petchiappan et al., 2024 |
| pAP405 | MalF TM-RcsF ($MalF_{1-38}$-$RcsF_{17-C}$) with T25 tag at N-terminal cloned in pKT25 | pKT25 linearized with EW3 and EW4; Insert is gBlock AP_FMF25 |
| pAP406 | RcsF A55K (with MalF TM at N-terminal) with T25 tag in pKT25 | pAP405 template with primers AP561 and AP562 (SDM) |
| pAP411 | RcsF A55V (with MalF TM at N-terminal) with T25 tag in pKT25 | pAP405 template with primers AP773 and AP774 (SDM) |

| | | |
|---|---|---|
| pAP412 | RcsF L58Y (with MalF TM at N-terminal) with T25 tag in pKT25 | pAP405 template with primers AP775 and AP776 (SDM) |
| pAP413 | MalF TM helix with T25 tag at N-terminal cloned in pKT25 | pAP405 template with primers AP397and EW4 |
| pAP415 | RcsF P62L (with MalF TM at N-terminal) with T25 tag in pKT25 | pAP405 template with primers AP819and AP 820 (SDM) |
| pAP417 | RcsF K61A (with MalF TM at N-terminal) with T25 tag in pKT25 | pAP405 template with primers AP903 and AP904 (SDM) |
| pAP418 | RcsF F63A (with MalF TM at N-terminal) with T25 tag in pKT25 | pAP405 template with primers AP905 and AP906 (SDM) |
| pAP419 | RcsF R64A (with MalF TM at N-terminal) with T25 tag in pKT25 | pAP405 template with primers AP907and AP908 (SDM) |
| **IgaA plasmids:** | | |
| pEAW1 | IgaA with T18 tag at C-terminal cloned in pUT18 | Wall et al., 2020[3] |
| pEAW1C4S | IgaA C404S C425S C498S C504S (C4S)-T18 tag in pUT18 | Petchiappan et al., 2024[4] |
| pEAW1L | IgaA L643P- T18 tag in pUT18 | Wall et al., 2020[3] |
| pEAW1peri | IgaA Δperi$_{384\text{-}649}$ with T18 tag at C-terminal cloned in pUT18 | Wall et al., 2020[3] |
| pEAW2 | IgaA with T25 tag at C-terminal cloned in pKNT25 | Wall et al., 2020[3] |
| pAP101 | IgaA with cyt1 and cyt2 domains deleted (Δ36-181 Δ263-329) -T18 tag at C-terminal cloned in pUT18 | Petchiappan et al., 2024[4] |
| pAP1094 | IgaA A94D – T18 tag in pUT18 | pEAW1 template with primers AP553 and AP554 (SDM) |
| pAP1097 | IgaA R97A– T18 tag in pUT18 | pEAW1 template with primers AP143 and AP144 (SDM) |
| pAP10972 | IgaA R97S– T18 tag in pUT18 | pEAW1 template with primers AP681 and AP682 (SDM) |
| pAP10468 | IgaA V468A– T18 tag in pUT18 | pEAW1 template with primers AP827 and AP828 (SDM) |
| pAP10469 | IgaA S469A– T18 tag in pUT18 | pEAW1 template with primers AP637 and AP638 (SDM) |
| pAP10471 | IgaA S471A– T18 tag in pUT18 | pEAW1 template with primers AP469 and AP470 (SDM) |
| pAP10472 | IgaA L472A– T18 tag in pUT18 | pEAW1 template with primers AP707 and AP708 (SDM) |
| pAP10475 | IgaA A475E– T18 tag in pUT18 | pEAW1 template with primers AP423 and AP424 (SDM) |
| pAP10476 | IgaA I476A– T18 tag in pUT18 | pEAW1 template with primers AP779 and AP780 (SDM) |

| | | |
|---|---|---|
| pAP10478 | IgaA K478A– T18 tag in pUT18 | pEAW1 template with primers AP735 and AP736 (SDM) |
| pAP10479 | IgaA S479A– T18 tag in pUT18 | pEAW1 template with primers AP639 and AP640 (SDM) |
| pAP10480 | IgaA G480A– T18 tag in pUT18 | pEAW1 template with primers AP709 and AP710 (SDM) |
| pAP10481 | IgaA M481A– T18 tag in pUT18 | pEAW1 template with primers AP711 and AP712 (SDM) |
| pAP10483 | IgaA L483A– T18 tag in pUT18 | pEAW1 template with primers AP809 and AP810 (SDM) |
| pAP10506 | IgaA R506A– T18 tag in pUT18 | pEAW1 template with primers AP589 and AP590 (SDM) |
| pAP10509 | IgaA N509A– T18 tag in pUT18 | pEAW1 template with primers AP299 and AP300 (SDM) |
| pAP10513 | IgaA N513A– T18 tag in pUT18 | pEAW1 template with primers AP591 and AP592 (SDM) |
| pAP10514 | IgaA L514A– T18 tag in pUT18 | pEAW1 template with primers AP939 and AP940 (SDM) |
| pAP10516 | IgaA N516A– T18 tag in pUT18 | pEAW1 template with primers AP813 and AP814 (SDM) |
| pAP10525 | IgaA K525A– T18 tag in pUT18 | pEAW1 template with primers AP713 and AP714 (SDM) |
| pAP10531 | IgaA K531A– T18 tag in pUT18 | pEAW1 template with primers AP783 and AP784 (SDM) |
| pAP10533 | IgaA D533A– T18 tag in pUT18 | pEAW1 template with primers AP785 and AP786 (SDM) |
| pAP10534 | IgaA G534A– T18 tag in pUT18 | pEAW1 template with primers AP891 and AP892 (SDM) |
| pAP10535 | IgaA V535A– T18 tag in pUT18 | pEAW1 template with primers AP787 and AP788 (SDM) |
| pAP10536 | IgaA N536A– T18 tag in pUT18 | pEAW1 template with primers AP415 and AP416 (SDM) |
| pAP10537 | IgaA V537A– T18 tag in pUT18 | pEAW1 template with primers AP789 and AP790 (SDM) |
| pAP10538 | IgaA L538A– T18 tag in pUT18 | pEAW1 template with primers AP791 and AP792 (SDM) |
| pAP10540 | IgaA R540A– T18 tag in pUT18 | pEAW1 template with primers AP815 and AP816 (SDM) |
| pAP10542 | IgaA V542A– T18 tag in pUT18 | pEAW1 template with primers AP301 and AP302 (SDM) |
| pAP10543 | IgaA S543A– T18 tag in pUT18 | pEAW1 template with primers AP303 and AP304 (SDM) |
| pAP10545 | IgaA E545K– T18 tag in pUT18 | pEAW1 template with primers AP185 and AP186 (SDM) |

| | | |
|---|---|---|
| pAP10546 | IgaA S546A– T18 tag in pUT18 | pEAW1 template with primers AP305 and AP306 (SDM) |
| pAP10549 | IgaA N549A– T18 tag in pUT18 | pEAW1 template with primers AP307 and AP308 (SDM) |
| pAP104722 | IgaA L472A L483A– T18 tag in pUT18 | pAP10472 template with primers AP809 and AP810 (SDM) |
| pAP105382 | IgaA L538A L483A– T18 tag in pUT18 | pAP10538 template with primers AP809 and AP810 (SDM) |
| **RcsD plasmids:** | | |
| pEAW7 | RcsD with T18 tag at C-terminal cloned in pUT18 | Wall et al., 2020[3] |
| pEAW7peri | RcsD Δperi$_{45\text{-}304}$ with T18 tag at C-terminal cloned in pUT18 | Wall et al., 2020[3] |
| pEAW7s | RcsD cytoplasmic domain (326-C) with T18 tag at C-terminal cloned in pUT18 | Wall et al., 2020[3] |
| pEAW8 | RcsD with T25 tag at C-terminal cloned in pKNT25 | Wall et al., 2020[3] |
| pEAW8peri | RcsD Δperi$_{45\text{-}304}$ with T25 tag at C-terminal cloned in pKNT25 | Wall et al., 2020[3] |
| pEAW8s | RcsD cytoplasmic domain (326-C) with T25 tag at C-terminal cloned in pKNT25 | Wall et al., 2020[3] |
| pEAW8T | RcsD T411A with T25 tag at C-terminal cloned in pKNT25 | Wall et al., 2020[3] |
| pEAW11 | RcsD cloned in pBAD24 | Wall et al., 2020[3] |
| pEAW11T | RcsD T411A cloned in pBAD24 | Wall et al., 2020[3] |
| pEAW11peri | RcsD Δperi$_{45\text{-}304}$ in pBAD24 | Wall et al., 2020[3] |
| pAP1102 | RcsD T411A Δperi (Δ45-304) cloned in pBAD24 | pEAW11peri template with primers AP169 and AP170 (SDM) |
| pAP1107 | RcsD$_{Kp}$ cloned in pBAD24 vector | pBAD24 with primers EW55fix and EW56fix; gene PCR amplified with primers AP857 and AP858 |
| pAP7029 | RcsD V29A – T18 tag in pUT18 | pEAW7 template with primers AP607 and AP608 (SDM) |
| pAP7036 | RcsD V36A – T18 tag in pUT18 | pEAW7 template with primers AP867 and AP868 (SDM) |
| pAP7069 | RcsD Y69A – T18 tag in pUT18 | pEAW7 template with primers AP887 and AP888 (SDM) |
| pAP7134e | RcsD T134E – T18 tag in pUT18 | pEAW7 template with primers AP505 and AP506 (SDM) |
| pAP7137 | RcsD G137A – T18 tag in pUT18 | pEAW7 template with primers AP873 and AP874 (SDM) |
| pAP7138 | RcsD A138S – T18 tag in pUT18 | pEAW7 template with primers AP715 and AP716 (SDM) |

| | | |
|---|---|---|
| pAP7153 | RcsD N153A – T18 tag in pUT18 | pEAW7 template with primers AP831 and AP832 (SDM) |
| pAP7153D | RcsD N153D – T18 tag in pUT18 | pEAW7 template with primers AP437 and AP438 (SDM) |
| pAP7191 | RcsD N191A – T18 tag in pUT18 | pEAW7 template with primers AP833 and AP834 (SDM) |
| pAP7192 | RcsD A192S – T18 tag in pUT18 | pEAW7 template with primers AP649 and AP650 (SDM) |
| pAP7193 | RcsD L193A – T18 tag in pUT18 | pEAW7 template with primers AP835and AP836 (SDM) |
| pAP7194 | RcsD D194A – T18 tag in pUT18 | pEAW7 template with primers AP623 and AP624 (SDM) |
| pAP7220 | RcsD N220A – T18 tag in pUT18 | pEAW7 template with primers AP219 and AP220 (SDM) |
| pAP7222 | RcsD P222A – T18 tag in pUT18 | pEAW7 template with primers AP913 and AP914 (SDM) |
| pAP7223 | RcsD G223A – T18 tag in pUT18 | pEAW7 template with primers AP915 and AP916 (SDM) |
| pAP7224 | RcsD H224A – T18 tag in pUT18 | pEAW7 template with primers AP917 and AP918 (SDM) |
| pAP7225 | RcsD L225A – T18 tag in pUT18 | pEAW7 template with primers AP309 and AP310 (SDM) |
| pAP7311 | RcsD L311A – T18 tag in pUT18 | pEAW7 template with primers AP877 and AP878 (SDM) |
| pAP7319 | RcsD L319A – T18 tag in pUT18 | pEAW7 template with primers AP613 and AP614 (SDM) |
| pAP7326 | RcsD G326A – T18 tag in pUT18 | pEAW7 template with primers AP617 and AP618 (SDM) |
| pAP7330 | RcsD F330A – T18 tag in pUT18 | pEAW7 template with primers AP885 and AP886 (SDM) |
| pAP7409 | RcsD Q409A – T18 tag in pUT18 | pEAW7 template with primers AP837 and AP838 (SDM) |
| pAP7410 | RcsD A410G – T18 tag in pUT18 | pEAW7 template with primers AP629 and AP630 (SDM) |
| pAP7411 | RcsD T411K – T18 tag in pUT18 | pEAW7 template with primers AP117 and AP118 (SDM) |
| pAP7412 | RcsD I412A – T18 tag in pUT18 | pEAW7 template with primers AP839 and AP840 (SDM) |
| pAP7413 | RcsD N413A – T18 tag in pUT18 | pEAW7 template with primers AP841and AP842 (SDM) |
| pAP7414 | RcsD N414A – T18 tag in pUT18 | pEAW7 template with primers AP843 and AP844(SDM) |
| pAP7415 | RcsD E415A – T18 tag in pUT18 | pEAW7 template with primers AP845 and AP846 (SDM) |

| | | |
|---|---|---|
| pAP7416 | RcsD L416A – T18 tag in pUT18 | pEAW7 template with primers AP847 and AP848 (SDM) |
| pAP7919 | RcsD L319A Δperi$_{45\text{-}304}$ with T18 tag in pUT18 | pEAW7peri template - primers AP613 and AP614 (SDM) |
| pAP7929 | RcsD V29A Δperi$_{45\text{-}304}$ with T18 tag in pUT18 | pEAW7peri template - primers AP607 and AP608 (SDM) |
| pAP804 | RcsD T411A Δperi$_{45\text{-}304}$ with T25 tag at C-terminal cloned in pKNT25 | Petchiappan et al., 2024[4] |
| pAP8919 | RcsD L319A Δperi$_{45\text{-}304}$ with T25 tag in pKNT25 | pEAW8peri template - primers AP607 and AP608 (SDM) |
| pAP8929 | RcsD V29A Δperi$_{45\text{-}304}$ with T25 tag in pKNT25 | pEAW8peri template - primers AP613 and AP614 (SDM) |
| pAP11029 | RcsD V29A cloned in pBAD24 | pEAW11 template with primers AP607 and AP608 (SDM) |
| pAP11033 | RcsD T33A cloned in pBAD24 | pEAW11 template with primers AP865 and AP866 (SDM) |
| pAP11036 | RcsD V36A cloned in pBAD24 | pEAW11 template with primers AP867 and AP868 (SDM) |
| pAP11069 | RcsD Y69A cloned in pBAD24 | pEAW11 template with primers AP887 and AP888 (SDM) |
| pAP11134 | RcsD T134A cloned in pBAD24 | pEAW11 template with primers AP871 and AP872 (SDM) |
| pAP11134e | RcsD T134E cloned in pBAD24 | pEAW11 template with primers AP505 and AP506 (SDM) |
| pAP11137 | RcsD G137A cloned in pBAD24 | pEAW11 template with primers AP873 and AP874 (SDM) |
| pAP11138 | RcsD A138S cloned in pBAD24 | pEAW11 template with primers AP715 and AP716 (SDM) |
| pAP11153 | RcsD N153A cloned in pBAD24 | pEAW11 template with primers AP831 and AP832 (SDM) |
| pAP11153D | RcsD N153D cloned in pBAD24 | pEAW11 template with primers AP437 and AP438 (SDM) |
| pAP11191 | RcsD N191A cloned in pBAD24 | pEAW11 template with primers AP833 and AP834 (SDM) |
| pAP11192 | RcsD A192S cloned in pBAD24 | pEAW11 template with primers AP649 and AP650 (SDM) |
| pAP11193 | RcsD L193A cloned in pBAD24 | pEAW11 template with primers AP835and AP836 (SDM) |
| pAP11194 | RcsD D194A cloned in pBAD24 | pEAW11 template with primers AP623 and AP624 (SDM) |
| pAP11220 | RcsD N220A cloned in pBAD24 | pEAW11 template with primers AP219 and AP220 (SDM) |
| pAP11222 | RcsD P222A cloned in pBAD24 | pEAW11 template with primers AP913 and AP914 (SDM) |
| pAP11223 | RcsD G223A cloned in pBAD24 | pEAW11 template with primers AP915 and AP916 (SDM) |
| pAP11224 | RcsD H224A cloned in pBAD24 | pEAW11 template with primers AP917 and AP918 (SDM) |

| | | |
|---|---|---|
| pAP11225 | RcsD L225A cloned in pBAD24 | pEAW11 template with primers AP309 and AP310 (SDM) |
| pAP11311 | RcsD L311A cloned in pBAD24 | pEAW11 template with primers AP877 and AP878 (SDM) |
| pAP11312 | RcsD P312A cloned in pBAD24 | pEAW11 template with primers AP879 and AP880 (SDM) |
| pAP11315 | RcsD L315A cloned in pBAD24 | pEAW11 template with primers AP881 and AP882 (SDM) |
| pAP11316 | RcsD N316A cloned in pBAD24 | pEAW11 template with primers AP883 and AP884(SDM) |
| pAP11319 | RcsD L319A cloned in pBAD24 | pEAW11 template with primers AP613 and AP614 (SDM) |
| pAP11326 | RcsD G326A cloned in pBAD24 | pEAW11 template with primers AP617 and AP618 (SDM) |
| pAP11328 | RcsD T328A cloned in pBAD24 | pEAW11 template with primers AP541 and AP542 (SDM) |
| pAP11330 | RcsD F330A cloned in pBAD24 | pEAW11 template with primers AP885 and AP886 (SDM) |
| pAP11409 | RcsD Q409A cloned in pBAD24 | pEAW11 template with primers AP837 and AP838 (SDM) |
| pAP11410 | RcsD A410G cloned in pBAD24 | pEAW11 template with primers AP629 and AP630 (SDM) |
| pAP11411 | RcsD T411K cloned in pBAD24 | pEAW11 template with primers AP117 and AP118 (SDM) |
| pAP11412 | RcsD I412A cloned in pBAD24 | pEAW11 template with primers AP839 and AP840 (SDM) |
| pAP11413 | RcsD N413A cloned in pBAD24 | pEAW11 template with primers AP841and AP842 (SDM) |
| pAP11414 | RcsD N414A cloned in pBAD24 | pEAW11 template with primers AP843 and AP844(SDM) |
| pAP11415 | RcsD E415A cloned in pBAD24 | pEAW11 template with primers AP845 and AP846 (SDM) |
| pAP11416 | RcsD L416A cloned in pBAD24 | pEAW11 template with primers AP847 and AP848 (SDM) |
| pAP113112 | RcsD L311A L319A cloned in pBAD24 | pAP113112 template with primers AP613 and AP614 (SDM) |
| pAP113192 | RcsD L319A T134E cloned in pBAD24 | pAP113192 template with primers AP613 and AP614 (SDM) |
| pAP114112 | RcsD N153A T411A cloned in pBAD24 | pAP114112 template with primers AP831 and AP832 (SDM) |
| **RcsC plasmids:** | | |
| pEAW13 | RcsC cloned in pBAD24 | Wall et al., 2020[3] |
| pAP1312 | $RcsC_{Kp}$ cloned in pBAD24 vector | pBAD24 with primers EW55fix and EW56fix; gene PCR amplified with primers AP909 and AP910 |
| pAP13021 | RcsC L21A cloned in pBAD24 | pEAW13 template with primers AP893 and AP894 (SDM) |
| pAP13024 | RcsC V24A cloned in pBAD24 | pEAW13 template with primers AP921 and AP922 (SDM) |

| | | |
|---|---|---|
| pAP13028 | RcsC L28A cloned in pBAD24 | pEAW13 template with primers AP895 and AP896 (SDM) |
| pAP13031 | RcsC F31A cloned in pBAD24 | pEAW13 template with primers AP897 and AP898 (SDM) |
| pAP13035 | RcsC F35A cloned in pBAD24 | pEAW13 template with primers AP899 and AP900 (SDM) |
| pAP13042 | RcsC H42A cloned in pBAD24 | pEAW13 template with primers AP901 and AP902(SDM) |
| pEAW26 | RcsC with MalF-TM ($RcsC_{1-19}$-$MalF_{2-59}$-$RcsC_{334-C}$) in pBAD24 | Wall et al., 2020[3] |

**Table S5: List of primers used in this study**

| Name | Sequence (5'-3') |
|---|---|
| AP117 | CTCATACAGCTCGTTATTGATTTTCGCCTGAATAATCCCCTGAT |
| AP118 | ATCAGGGGATTATTCAGGCGAAAATCAATAACGAGCTGTATGAG |
| AP137 | GCGTAAACTCACACCTGAAGAACGTAGCGCC |
| AP138 | GCCTTTCATCCATGAGAGAGTGAATTTCAGCGGCAT |
| AP143 | GTCGGTAGAGATGCCGTACGCCGTGATAGCGTGTGTCAG |
| AP144 | CTGACACACGCTATCACGGCGTACGGCATCTCTACCGAC |
| AP169 | CAGCTCGTTATTGATCGCCGCCTGAATAATCCCCT |
| AP170 | AGGGGATTATTCAGGCGGCGATCAATAACGAGCTG |
| AP185 | GTTATCCAGCGATTTCGCACTCACCGGGC |
| AP186 | GCCCGGTGAGTGCGAAATCGCTGGATAAC |
| AP213 | TTCAGGCAGCTTTCAAAATAGGCCTGAATCTCCATC |
| AP214 | GATGGAGATTCAGGCCTATTTTGAAAGCTGCCTGAA |
| AP219 | GATGTCCTGGCTGGGCAAAGGTAGTACGCAATGTAAAGTAATG |
| AP220 | CATTACTTTACATTGCGTACTACCTTTGCCCAGCCAGGACATC |
| AP299 | ATTGACCAGCGCAGCTTTCAGTCGCACACAGTCATCTTTG |
| AP300 | CAAAGATGACTGTGTGCGACTGAAAGCTGCGCTGGTCAAT |
| AP301 | GCGATTCCGCACTCGCCGGGCGTAATAAC |
| AP302 | GTTATTACGCCCGGCGAGTGCGGAATCGC |
| AP303 | ATCCAGCGATTCCGCAGCCACCGGGCGTAATAAC |
| AP304 | GTTATTACGCCCGGTGGCTGCGGAATCGCTGGAT |
| AP305 | GGTTATCCAGCGCTTCCGCACTCACCGGG |
| AP306 | CCCGGTGAGTGCGGAAGCGCTGGATAACC |
| AP307 | GTTGCCACCAGGGCATCCAGCGATTCCGCACTC |
| AP308 | GAGTGCGGAATCGCTGGATGCCCTGGTGGCAAC |
| AP309 | CCACGACCGTTGCCGCATGTCCTGGCTGGTTAAA |
| AP310 | TTTAACCAGCCAGGACATGCGGCAACGGTCGTGG |
| AP397 | CCG AAT TCT TAG TTA CCC TTG TGC GTA CAT TAA AAC AAC AAG GTA ACC |
| AP415 | CACCGGGCGTAATAACACCGCCACGCCATCGAGCTTCCC |
| AP416 | GGGAAGCTCGATGGCGTGGCGGTGTTATTACGCCCGGTG |
| AP423 | CATGCCGGATTTTTGAATTTCTGAACGTAATGAGGCACT |

| | |
|---|---|
| AP424 | AGTGCCTCATTACGTTCAGAAATTCAAAAATCCGGCATG |
| AP437 | ATCAGCACCAGACTGTCATCCTGACCATTCAGGTA |
| AP438 | TACCTGAATGGTCAGGATGACAGTCTGGTGCTGAT |
| AP469 | AATTGCTGAACGTAATGCGGCACTGACGCGAGAT |
| AP470 | ATCTCGCGTCAGTGCCGCATTACGTTCAGCAATT |
| AP505 | CATTTTCTGCGCCCCACAATTCATCCAGATAAGTGGACATCC |
| AP506 | GGATGTCCACTTATCTGGATGAATTGTGGGGCGCAGAAAATG |
| AP541 | GCGGAATGTGGCATAGCCAAATAACGCCAGCGC |
| AP542 | GCGCTGGCGTTATTTGGCTATGCCACATTCCGC |
| AP553 | GCCGTAACGCGTGATATCGTGTGTCAGCATCAT |
| AP554 | ATGATGCTGACACACGATATCACGCGTTACGGC |
| AP559 | GAG CTC GAA TTC GCT AGC CCA AAA AAA CG |
| AP560 | AAG CTT GGC TGT TTT GGC GGA TGA G |
| AP561 | GCCGACTAATTCTTCTTTATTGGTATAAATTCGGACCGGCGTG |
| AP562 | CACGCCGGTCCGAATTTATACCAATAAAGAAGAATTAGTCGGC |
| AP589 | ACCAGCGCATTTTTCAGTGCCACACAGTCATCTTTGGC |
| AP590 | GCCAAAGATGACTGTGTGGCACTGAAAAATGCGCTGGT |
| AP591 | GTCTTTACTGTTGCCGAGAGCGACCAGCGCATTTTTCAGT |
| AP592 | ACTGAAAAATGCGCTGGTCGCTCTCGGCAACAGTAAAGAC |
| AP607 | CACCCATCGTCACCAGTAACGCAATGATCAACAGTAAAAAG |
| AP608 | CTTTTTACTGTTGATCATTGCGTTACTGGTGACGATGGGTG |
| AP613 | CCAGCGCCAGCGCACCGATGTTCAGCAGCAGTG |
| AP614 | CACTGCTGCTGAACATCGGTGCGCTGGCGCTGG |
| AP617 | GCGGAATGTGGTATAGGCAAATAACGCCAGCGC |
| AP618 | GCGCTGGCGTTATTTGCCTATACCACATTCCGC |
| AP623 | ATTCCTGATCGTGAGCCAGCAGGCCGAGC |
| AP624 | GCTCGGCCTGCTGGCTCACGATCAGGAAT |
| AP629 | ACAGCTCGTTATTGATCGTACCCTGAATAATCCCCTGATG |
| AP630 | CATCAGGGGATTATTCAGGGTACGATCAATAACGAGCTGT |
| AP637 | GCTGAACGTAATGAGGCAGCGACGCGAGATTCATCTTC |
| AP638 | GAAGATGAATCTCGCGTCGCTGCCTCATTACGTTCAGC |
| AP639 | CAATACCATGCCGGCTTTTTGAATTGCTGAACGTAATGAGGC |
| AP640 | GCCTCATTACGTTCAGCAATTCAAAAAGCCGGCATGGTATTG |
| AP649 | GCGTTCATCGAGGGAGTTGGCCTGTTGCA |
| AP650 | TGCAACAGGCCAACTCCCTCGATGAACGC |
| AP681 | GTAGAGATGCCGTAACTCGTGATAGCGTGTGTC |
| AP682 | GACACACGCTATCACGAGTTACGGCATCTCTAC |
| AP707 | TTTGAATTGCTGAACGCGCTGAGGCACTGACGCGAGATTCATCTTC |
| AP708 | GAAGATGAATCTCGCGTCAGTGCCTCAGCGCGTTCAGCAATTCAAA |
| AP709 | AAATCATCAAGCAATACCATGGCGGATTTTTGAATTGCTGAAC |
| AP710 | GTTCAGCAATTCAAAAATCCGCCATGGTATTGCTTGATGATTT |
| AP711 | GCCAAAATCATCAAGCAATACCGCGCCGGATTTTTGAATTGCTGAA |
| AP712 | TTCAGCAATTCAAAAATCCGGCGCGGTATTGCTTGATGATTTTGGC |

| | |
|---|---|
| AP713 | GCGTTGGCGCGCTACCAGCGTCCCAGTCTTT |
| AP714 | AAAGACTGGGACGCGCTGGTAGCGCGCCAACGC |
| AP715 | CCACGGTACATTTCTGAGCCCCACAATGTATCCA |
| AP716 | TGGATACATTGTGGGGCTCAGAAAATGTACCGTGG |
| AP735 | CAATACCATGCCGGACGCTTGAATTGCTGAACGTAATGAGGCACTGA |
| AP736 | TCAGTGCCTCATTACGTTCAGCAATTCAAGCGTCCGGCATGGTATTG |
| AP773 | GTTTGCCGACTAATTCTTCTACATTGGTATAAATTCGGACC |
| AP774 | GGTCCGAATTTATACCAATGTAGAAGAATTAGTCGGCAAAC |
| AP775 | GAACGGTTTGCCGACATATTCTTCTGCATTGGTATAAATTCGGAC |
| AP776 | GTCCGAATTTATACCAATGCAGAAGAATATGTCGGCAAACCGTTC |
| AP779 | CATGCCGGATTTTTGAGCTGCTGAACGTAATGAGGCACTGA |
| AP780 | TCAGTGCCTCATTACGTTCAGCAGCTCAAAAATCCGGCATG |
| AP783 | CACGCCATCGAGCGCCCCGGCGTTGGCG |
| AP784 | CGCCAACGCCGGGGCGCTCGATGGCGTG |
| AP785 | CACATTCACGCCAGCGAGCTTCCCGGC |
| AP786 | GCCGGGAAGCTCGCTGGCGTGAATGTG |
| AP787 | GCGTAATAACACATTCGCGCCATCGAGCTTCCC |
| AP788 | GGGAAGCTCGATGGCGCGAATGTGTTATTACGC |
| AP789 | CACCGGGCGTAATAACGCATTCACGCCATCGAG |
| AP790 | CTCGATGGCGTGAATGCGTTATTACGCCCGGTG |
| AP791 | CACTCACCGGGCGTAATGCCACATTCACGCCATCGA |
| AP792 | TCGATGGCGTGAATGTGGCATTACGCCCGGTGAGTG |
| AP809 | TGTCGCCAAAATCATCAAGCGCTACCATGCCGGATTTTTGAATTG |
| AP810 | CAATTCAAAAATCCGGCATGGTAGCGCTTGATGATTTTGGCGACA |
| AP813 | CGTCCCAGTCTTTACTGGCGCCGAGATTGACCAGCG |
| AP814 | CGCTGGTCAATCTCGGCGCCAGTAAAGACTGGGACG |
| AP815 | GCACTCACCGGGGCTAATAACACATTCACGCCATCGA |
| AP816 | TCGATGGCGTGAATGTGTTATTAGCCCCGGTGAGTGC |
| AP819 | ACCGAGATCGCGGAACAGTTTGCCGACTAATTC |
| AP820 | GAATTAGTCGGCAAACTGTTCCGCGATCTCGGT |
| AP827 | AACGTAATGAGGCACTGGCGCGAGATTCATCTTCC |
| AP828 | GGAAGATGAATCTCGCGCCAGTGCCTCATTACGTT |
| AP831 | TGAGATCAGCACCAGACTGGCATCCTGACCATTCAGGTAAT |
| AP832 | ATTACCTGAATGGTCAGGATGCCAGTCTGGTGCTGATCTCA |
| AP833 | GCGTTCATCGAGGGCGGCGGCCTGTTGCAACATC |
| AP834 | GATGTTGCAACAGGCCGCCGCCCTCGATGAACGC |
| AP835 | CTTTCGCGTTCATCGGCGGCGTTGGCCTGTTG |
| AP836 | CAACAGGCCAACGCCGCCGATGAACGCGAAAG |
| AP837 | TTGATCGTCGCCGCAATAATCCCCTGATGCTGTTCCG |
| AP838 | CGGAACAGCATCAGGGGATTATTGCGGCGACGATCAA |
| AP839 | CTCATACAGCTCGTTATTGGCCGTCGCCTGAATAATCCCC |
| AP840 | GGGGATTATTCAGGCGACGGCCAATAACGAGCTGTATGAG |
| AP841 | GGATCTCATACAGCTCGTTAGCGATCGTCGCCTGAATAATCC |

| | |
|---|---|
| AP842 | GGATTATTCAGGCGACGATCGCTAACGAGCTGTATGAGATCC |
| AP843 | TCTCATACAGCTCGGCATTGATCGTCGCCTGAATAATCCC |
| AP844 | GGGATTATTCAGGCGACGATCAATGCCGAGCTGTATGAGA |
| AP845 | GCGGATCTCATACAGCGCGTTATTGATCGTCGC |
| AP846 | GCGACGATCAATAACGCGCTGTATGAGATCCGC |
| AP847 | GAACATGCGGATCTCATACGCCTCGTTATTGATCGTCGCC |
| AP848 | GGCGACGATCAATAACGAGGCGTATGAGATCCGCATGTTC- |
| AP857 | AGC AGG AGG AAT TCA ATG ACC CCG AAA AAA TTC TCC CTG ATC C |
| AP858 | AAA ACA GCC AAG CTT CTA CAG CAA GCT CTT GAC GTA AGC GTC |
| AP865 | GTACCATTACACCCATCGCCACCAGTAACACAATGAT |
| AP866 | ATCATTGTGTTACTGGTGGCGATGGGTGTAATGGTAC |
| AP867 | GGCGCTTTGTACCATTGCACCCATCGTCACCAG |
| AP868 | CTGGTGACGATGGGTGCAATGGTACAAAGCGCC |
| AP871 | TCTGCGCCCCACAATGCATCCAGATAAGTGGAC |
| AP872 | GTCCACTTATCTGGATGCATTGTGGGGCGCAGA |
| AP873 | CGGTACATTTTCTGCGGCCCACAATGTATCCAG |
| AP874 | CTGGATACATTGTGGGCCGCAGAAAATGTACCG |
| AP877 | GATGTTCAGCAGCAGTGGCGCCAGAATGTTTTGCAACGTA |
| AP878 | TACGTTGCAAAACATTCTGGCGCCACTGCTGCTGAACATC |
| AP879 | GATGTTCAGCAGCAGTGCCAGCAGAATGTTTTGCA |
| AP880 | TGCAAAACATTCTGCTGGCACTGCTGCTGAACATC |
| AP881 | CAGCAAACCGATGTTCGCCAGCAGTGGCAGCAGA |
| AP882 | TCTGCTGCCACTGCTGGCGAACATCGGTTTGCTG |
| AP883 | GCCAGCAAACCGATGGCCAGCAGCAGTGGCAG |
| AP884 | CTGCCACTGCTGCTGGCCATCGGTTTGCTGGC |
| AP885 | GGAGAAATGGCGGGCTGTGGTATAGCCAAATAACGCCAGC |
| AP886 | GCTGGCGTTATTTGGCTATACCACAGCCCGCCATTTCTCC |
| AP887 | CTGCCAGGTCACGGCACGCCAGTTATCGACGCG |
| AP888 | CGCGTCGATAACTGGCGTGCCGTGACCTGGCAG |
| AP891 | TAATAACACATTCACGGCATCGAGCTTCCCGGC |
| AP892 | GCCGGGAAGCTCGATGCCGTGAATGTGTTATTA |
| AP893 | AGAGCACTAACGCCGCTGCTCTGAACATGTAGCGCG |
| AP894 | CGCGCTACATGTTCAGAGCAGCGGCGTTAGTGCTCT |
| AP895 | AACGGATGAAAAAGCAATCGCCAGCCAGAGCACTAACGCC |
| AP896 | GGCGTTAGTGCTCTGGCTGGCGATTGCTTTTTCATCCGTT |
| AP897 | ACGATGTAAAAAACGGATGAAGCAGCAATCAACAGCCAGAGCAC |
| AP898 | GTGCTCTGGCTGTTGATTGCTGCTTCATCCGTTTTTTACATCGT |
| AP899 | CTGATGTAACGCATTAACGATGTAAGCAACGGATGAAAAAGCAATCAACAGC |
| AP900 | GCTGTTGATTGCTTTTTCATCCGTTGCTTACATCGTTAATGCGTTACATCAG |
| AP901 | CCGATTCTCGCTGAGCTAACGCATTAACGATGTAAAAAACGGA |
| AP902 | TCCGTTTTTTACATCGTTAATGCGTTAGCTCAGCGAGAATCGG |
| AP903 | CGAGATCGCGGAACGGTGCGCCGACTAATTCTTCTG |
| AP904 | CAGAAGAATTAGTCGGCGCACCGTTCCGCGATCTCG |

| | |
|---|---|
| AP905 | CTTCACCGAGATCGCGGGCCGGTTTGCCGACTAATT |
| AP906 | AATTAGTCGGCAAACCGGCCCGCGATCTCGGTGAAG |
| AP907 | ACTTCACCGAGATCGGCGAACGGTTTGCCGAC |
| AP908 | GTCGGCAAACCGTTCGCCGATCTCGGTGAAGT |
| AP909 | AGC AGG AGG AAT TCA ATG AAA TAT CTT GCC TCT TTT CAT ACA ACC CTG AAG |
| AP910 | AAA ACA GCC AAG CTT TTA TTC CCG CCC TTT ACG CAC CCG |
| AP913 | CGTTGCCAGATGTCCTGCCTGGTTAAAGGTAGTAC |
| AP914 | GTACTACCTTTAACCAGGCAGGACATCTGGCAACG |
| AP915 | ACCGTTGCCAGATGTGCTGGCTGGTTAAAGG |
| AP916 | CCTTTAACCAGCCAGCACATCTGGCAACGGT |
| AP917 | ACGACCGTTGCCAGAGCTCCTGGCTGGTTAAAGG |
| AP918 | CCTTTAACCAGCCAGGAGCTCTGGCAACGGTCGT |
| AP923 | AAAAGCAATCAACAGCCAGGCCACTAACGCCAATGCTCTG |
| AP924 | CAGAGCATTGGCGTTAGTGGCCTGGCTGTTGATTGCTTTT |
| AP939 | TTTACTGTTGCCGGCATTGACCAGCGCATTTTTCAGTCGC |
| AP940 | GCGACTGAAAAATGCGCTGGTCAATGCCGGCAACAGTAAA |
| EW3 | TAC CCG GGG ATC CTC TAG AGT CG |
| EW4 | TAA CTA AGA ATT CGG CCG TCG TTT TAC AAC GTC |
| EW55fix | TGA ATT CCT CCT GCT AGC CCA AAA AAA CGG GTA TGG AG |
| EW56fix | AAG CTT GGC TGT TTT GGC GGA TGA GAG AAG ATT TTC AGC |
| EW81 | GCT ATG ACC ATG ATT ACC CCG AAA AAA TTC TCC CTG ATC CCC |
| EW82 | GCA GGC ATG CAA GCT CAG CAA GCT CTT GAC GTA AGC GTC |
| EW213 | ACCACGCCTGACAGACTAAGTAAGATGGGGAAAGC ATG AGC ACC ATT GTG ATT TTT TTA GCT GCT TTG C |
| EW214 | GACAGGGTAGCATAACCTGCCGCGCAAACGTG TTA TTC GAT AAG GCT TTC TGA AGG GGT GAT CAG TTG |

**Table S6: List of synthetic gene fragments (gBlocks) used in this study**

| Name | Sequence (5'-3') |
|---|---|
| AP_FMF25kp | GAGGATCCCCGGGTAATGGATGTCATTAAAAAGAAACATTGGTGG CAAAGCGACGCGCTGAAATGGTCAGTGCTAGGTCTGCTCGGCCTG CTGGTGGGTTACCTTGTTGTTTAATGTACGCACAAGGGTCCATGC TAAGCAGATCTCCCGTTGAGCCTGCTCAAAGCACTGCAACCCCGCC GGTTAAATCTGAGCCGAGCAAACCGCGCGCCACCCGCCCGGCCCC AGTACGTATTTACACCGATGCGTCAGAGCTGGTTGGTAAACCGTTC CGCGATCTGGGTGAAGTATCTGGCGAGTCCTGCCAGGCTTCGAATC AGGATTCTCCGCCGAATATCCCCACCGCGCGCAAGCGCCTGCAAA TTAATGCCGCGCGTATGAAAGCTAATGCCGTCCTGCTTCACCGCTG CGAAGTGACCAGCGGTACGCCAGGCTGCTACCGTCAGGCCGTTTG CCTGGGTTCAGCGCTTAACGTCTCGGCGCAATAACTAAGAATTCGG |

| | |
|---|---|
| AP_GF33kp | AGCGAATTCGAGCTCAGCAGGAGGAATTCAATGCGTGCTTTACCG ATCTGTTTGTTAGCACTCATGTTAAGCGGTTGTTCCATGCTAAGCA GATCTCCCGTTGAGCCTGCTCAAAGCACTGCAACCCCGCCGGTTAA ATCTGAGCCGAGCAAACCGCGCGCCACCCGCCCGGCCCCAGTACG TATTTACACCGATGCGTCAGAGCTGGTTGGTAAACCGTTCCGCGAT CTGGGTGAAGTATCTGGCGAGTCCTGCCAGGCTTCGAATCAGGATT CTCCGCCGAATATCCCCACCGCGCGCAAGCGCCTGCAAATTAATG CCGCGCGTATGAAAGCTAATGCCGTCCTGCTTCACCGCTGCGAAGT GACCAGCGGTACGCCAGGCTGCTACCGTCAGGCCGTTTGCCTGGG TTCAGCGCTTAACGTCTCGGCGCAATGAAAGCTTGGCTGTTTT |

**Table S7: Software and algorithms**

| **Name** | **Description** | **Reference** |
|---|---|---|
| CryoSPARC | Cryo-EM data processing software | Punjani et. al 2017[9] |
| TOPAZ | Cryo-EM data particle picking software | Bepler et. al 2019[10] |
| PHENIX | Structure refinement and analysis software | Liebschner et. al 2019[11] |
| BioWulf Cluster | Data processing and storage | NIH HPC[12] |
| DeepEMhancer | Cryo-EM map improvement software | Sanchez-Garcia et. al 2021[13] |
| ChimeraX | Structure analysis and image preparation software | Meng et. al 2023[14] |
| Chimera | Structure analysis and image preparation software | Pettersen et. al 2004[15] |
| GraphPad Prism | Data processing software | GraphPad Prism[16] |
| EPU | Data collection software | ThermoFisher |
| SerialEM | Data collection software | Mastronarde et. al[17] |
| Coot | Model building software | Emsley et. al 2010[18] |
| AlphaFold 3 | Structure prediction | Abramson et. al 2024[19] |
| Isolde | Model building software | Croll[20] |
